# Single-cell multiomics atlas of organoid development uncovers longitudinal molecular programs of cellular diversification of the human cerebral cortex

**DOI:** 10.1101/2022.03.17.484798

**Authors:** Ana Uzquiano, Amanda J. Kedaigle, Martina Pigoni, Bruna Paulsen, Xian Adiconis, Kwanho Kim, Tyler Faits, Surya Nagaraja, Noelia Antón-Bolaños, Chiara Gerhardinger, Ashley Tucewicz, Evan Murray, Xin Jin, Jason Buenrostro, Fei Chen, Silvia Velasco, Aviv Regev, Joshua Z. Levin, Paola Arlotta

## Abstract

Realizing the full utility of brain organoids as experimental systems to study human cortical development requires understanding whether organoids replicate the cellular and molecular events of this complex process precisely, reproducibly, and with fidelity to the embryo. Here we present a comprehensive single-cell transcriptomic, epigenetic, and spatial atlas of human cortical organoid development, comprising over 610,000 cells, spanning initial generation of neural progenitors through production of differentiated neuronal and glial subtypes. We define the lineage relationships and longitudinal molecular trajectories of cortical cell types during development in organoids, and show that developmental processes of cellular diversification in organoids correlate closely to endogenous ones, irrespective of metabolic state. Using this data, we identify genes with predicted human-specific roles in lineage establishment, and discover a developmental origin for the transcriptional diversity of human callosal projection neurons, a population that has undergone dramatic expansion and diversification during human evolution. Our work provides a comprehensive, single-cell molecular map of human corticogenesis *in vitro*, identifying developmental trajectories and molecular mechanisms associated with human cellular diversification.

## Introduction

Development of the human cerebral cortex is a protracted process that spans long periods of *in utero* development and thus is largely experimentally inaccessible (Silbereis *et al*., 2016). Although studies using human fetal tissue have proven critical to our current knowledge of human corticogenesis (Fietz *et al*., 2010; Hansen *et al*., 2010; Nowakowski *et al*., 2017; de la Torre-Ubieta *et al*., 2018; Polioudakis *et al*., 2019; Markenscoff-Papadimitriou *et al*., 2020; Song *et al*., 2020; Trevino *et al*., 2021), there is an outstanding need for experimental models that can be used to dissect the molecular logic governing human cortical development.

Human brain organoids have emerged as a powerful experimental system to investigate human brain development and neurodevelopmental diseases (Lancaster *et al*., 2013; Mariani *et al*., 2015; Bershteyn *et al*., 2017; Birey *et al*., 2017; Kanton *et al*., 2019; Klaus *et al*., 2019; Esk *et al*., 2020; Khan *et al*., 2020; Paulsen *et al*., 2022). Organoids generate a large diversity of cell types resembling those that populate the human developing cortex (Camp *et al*., 2015; Quadrato *et al*., 2017; Velasco *et al*., 2019; Yoon *et al*., 2019), and they are capable of doing so in a reproducible fashion (Velasco *et al*., 2019). Additionally, organoids broadly recapitulate the transcriptional and epigenetic profile of the endogenous fetal cortex (Velasco *et al*., 2019; Trevino *et al*., 2020; Gordon *et al*., 2021), and show physiologically-relevant features such as the presence of neuronal activity (Quadrato *et al*., 2017; Trujillo *et al*., 2019, 2020; Miura *et al*., 2020) and the ability to form long-distance connections (Giandomenico *et al*., 2019; Andersen *et al*., 2020; Miura *et al*., 2020).

While these features highlight the promise of human brain organoids as models for human brain development, the lineage relationships and developmental trajectories that generate each cell type remain largely not understood. This requires a comprehensive, single-cell level analysis of cell-type specification and diversification across long timelines of organoid development, and investigation of their fidelity to the molecular programs driving endogenous human corticogenesis. Such a map of human cortical development *in vitro* would be an invaluable resource to study events of human cortical cellular diversification, in particular of populations that have undergone striking species-specific expansion and diversification during human evolution, such as outer radial glia (oRG) and callosal projection neurons (CPN) of the upper cortical layers. This understanding will empower the use of organoids for experimental investigation of the developmental origin of human-specific cortical features and human neurodevelopmental abnormalities.

Here, we have assembled an unprecedented atlas of 613,242 single-cell transcriptomes and epigenomes from a reproducible human cortical organoid model (Velasco *et al*., 2019), sampled through extended developmental trajectories spanning 8 timepoints over 6 months of *in vitro* development, integrated with spatial transcriptomic profiling to define topographic organization of organoid cell types over time. We show that longitudinal events of cortical development *in vitro* mirror developmental milestones of *in utero* development, including proper acquisition of cell identity of the great majority of cortical cell types populating the organoids. Importantly, in this model, expression of metabolism-associated genes does not affect fidelity of cell identity for most organoid cell types. We define a lineage map that reveals the molecular logic underlying cell specification across the main human cortical lineages, and identify species-specific differences in lineage-associated genes and in transcription factors mediating fate specification in the human cortex. Notably, we discover that the recently-described transcriptional diversity in adult human callosal projection neurons (CPN) (Berg *et al*., 2021), a population that has undergone unusual expansion in the human cerebral cortex, emerges at early stages of corticogenesis, providing first demonstration of an early developmental appearance of this human-specific cortical feature.

## Results

### Single-cell-resolution transcriptomic, epigenomic, and spatial atlas of human cortical organoid development

To investigate the molecular characteristics and reproducibility of corticogenesis in human brain organoids, we built a longitudinal single-cell atlas comprising 8 timepoints across 6 months of organoid development, spanning processes from early stages of progenitor amplification to later astroglia production (Figure 1A). We profiled a total of 532,414 cells by single-cell RNA sequencing (scRNA-seq), combining our previously published datasets (Velasco *et al*., 2019; Paulsen *et al*., 2022) with an additional 218,240 newly-profiled cells from multiple timepoints. We analyzed organoids derived from multiple stem cell lines (2-6 lines per stage) and differentiation batches (n = 2-7 per line and stage; Figures 1A and S1A). Each organoid was profiled individually in order to evaluate organoid-to-organoid variability, producing an atlas comprising the single-cell transcriptome of a total of 83 single organoids (Figures 1A and S1A). We provide this data as a resource for interactive exploration at the Single Cell Portal: https://singlecell.broadinstitute.org/single_cell/study/SCP1756.

**Fig. 1.**
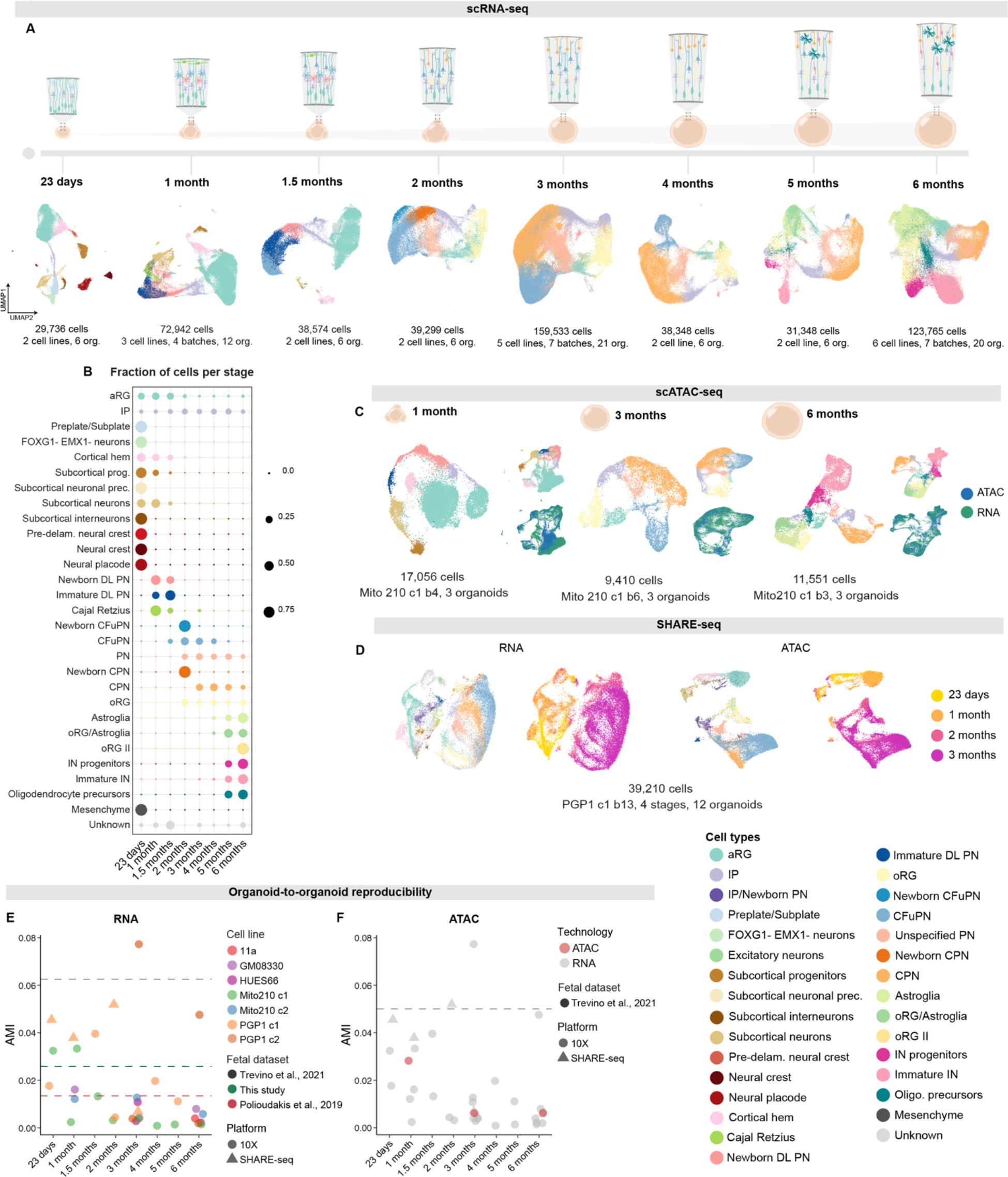
Single-cell transcriptomic and epigenetic landscape of developing cortical organoids. (A) scRNA-seq of organoids cultured for 23 days, 1 month, 1.5 months, 2 months, 3 months, 4 months, 5 months, and 6 months, represented by Uniform Manifold Approximation and Projections (UMAPs) of the data after Harmony batch correction. Cells are colored by cell type, and the number of cells per plot is indicated. Above, schematic of cell types during organoid development. (B) Fraction of cells belonging to each cell type at each timepoint. (C) scATAC-seq of organoids cultured for 1, 3, and 6 months, represented by UMAP. Cells are colored by cell type, and the number of cells per plot is indicated. Smaller inset UMAPs show scATAC-seq (blue) and scRNA-seq (green) data integration. (D) SHARE-seq of organoids cultured for 23 days, 1, 2 and 3 months, represented by UMAP. Left: cell types from all the times are shown in UMAP. Right: organoid stage is shown in UMAP. (E) Adjusted mutual information (AMI) scores between cell types and individual organoids for each scRNA-seq (10x) and SHARE-seq dataset, where lower scores indicate lower variability. Dotted lines represent the AMI scores of fetal cortex datasets. (F) Adjusted mutual information (AMI) scores between cell types and individual organoids in scATAC-seq (10x, green points). Gray points are repeated from Figure 1E and dotted lines represent the AMI scores of human fetal cortex datasets. Abbreviations: “c”, cell line clone; “b” differentiation batch; org., organoid; aRG: apical radial glia; IP: intermediate progenitor; prec., precursors; DL PN, deep-layer projection neurons; CFuPN, corticofugal projection neuron; CPN, callosal projection neuron; oRG, outer radial glia; IN: interneurons.

Notably, at the earliest timepoint characterized (23 days), the organoids contained not only diverse cell types characteristic of the telencephalon (expressing *FOXG1*), but also a large proportion (∼25-60%) of cells of other regions of the developing nervous system (Figures 1B, S1B, and S2). Cortical cell types included neural progenitors, primarily apical radial glia (aRG; *EMX1*, *SOX2*, *HES1*) and a smaller proportion of intermediate progenitors (IP; *EMX1*, *EOMES*, *INSM1*), as well as the first neurons populating the developing cortical plate (preplate/subplate; *EMX2*, *FOXP2*, *EOMES*). We also observed cells with a cortical hem identity (*WLS*, *RSPO1*, *RSPO2*), consistent with previous reports (Kadoshima *et al*., 2013). The non-cortical cell types included populations expressing markers of the developing midbrain (*LMX1B*, *TH*), hindbrain (*HOXB2*, *PHOX2A*, *PHOX2B*), thalamus (*NEFM*, *LHX9*), neural placode (*ISL1*, *SIX1*), and neural crest (*FOXD3*, *PAX3*) (Figures 1B and S1B). At this stage, organoids contained subcortical interneurons, expressing markers such as *LAMP5*, *LHX5*, and *GAD2* as well as markers characteristic of caudal regions of the neural tube, i.e., hindbrain (*PAX2, CRABP1, CRABP2, TFAP2A*) (Figure S2C, K). Thus, while later in development organoids have a definitive cortical identity, at the earliest stages of *in vitro* development organoids initiate production of multiple brain regions, similar to the early neural tube *in vivo*.

Over the course of organoid development, from 1 to 6 months *in vitro,* non-cortical cell types were progressively depleted, and cellular composition followed differentiation transitions mirroring those in the human cerebral cortex (Figures 1A, 1B, S1, and S2). A few non-cortical cells (∼8.1%) were still present at 1 month, but from 2 months onwards organoids were exclusively populated by cell types of the cerebral cortex (Figures 1A and 1B). As organoid differentiation progressed, the dominant neuronal progenitor types changed from aRG, to IP, to outer radial glia (oRG; *SOX2, HOPX, PEA15, LGALS3BP, MOXD1*) (Figures 1A, 1B, S1, and S2), as previously described in the fetal developing human cortex (Fietz *et al*., 2010; Hansen *et al*., 2010). The main neuronal types of the developing cortex appeared in the order observed *in vivo* (Angevine and Sidman, 1961; Rakic, 1974; Shen *et al*., 2006; Yue Huang *et al*., 2020): immature deep-layer projection neurons (DL PN; *FEZF1, NEUROD2, NEUROD6, TBR1, PDE1A*) appeared at 1 month, followed by deep-layer corticofugal projection neurons (CFuPN; *SOX5, LDB2, CRYM, TLE4*) by 1.5 months *in vitro*. Neurogenesis of callosal projection neurons (CPN; *UNC5A, EPHA4, BHLHE22, PLXNA4, SATB2*) began at approximately 2 months *in vitro*, concomitant with the emergence of oRG, and consistent with the production of CPN from oRG in the endogenous cortex (Lukaszewicz *et al*., 2005; Betizeau *et al*., 2013; Nowakowski *et al*., 2016).

At two months we also observed emergence of an excitatory neuron population marked by expression of *TBR1*, *NEUROD2*, and *NEUROD6*, but which lacked other canonical markers of subtype identity (labeled here as ‘unspecified PN’). By three months in culture, CPN were the most abundant excitatory neuron population. At four months, we observed a small population of glial cells with molecular features characteristic of astroglia (*GJA1*, *S100B, SPARC*). By five months *in vitro*, oligodendrocyte precursors (*OLIG1*, *OLIG2*, *PDGFRA*) and cycling interneuron progenitors (*BIRC5*, *DLX2*, *DLX5*) developed, concomitant with the appearance of immature interneurons (*GAD2*, *DLX2*, *DLX5*). Finally, at six months, organoids contained a great abundance of astroglia and immature interneurons (Figures 1A, 1B, S1, and S2). This interneuron population was molecularly distinct from the *GAD2*-positive population observed at 23 days, and, importantly, expressed *FOXG1,* indicating that this later-emerging population has a telencephalic origin. This interneuron population expressed *SP8*, *SCGN*, *PROX1* and *CALB2* (Figure S2L), suggesting that these may be olfactory bulb-like interneurons derived from cortical progenitors, as previously described in mice (Kohwi *et al*., 2007; Young *et al*., 2007; Fuentealba *et al*., 2015); alternatively, they may constitute a recently described dorsally-derived interneuron population, which molecularly resembles interneurons derived from the caudal ganglionic eminence (Delgado *et al*., 2022).

We next assessed if these longitudinal events of cellular production each emerged reproducibly across development in all organoids. We calculated the adjusted mutual information (AMI) score for each of the timepoints sampled, measuring the dependence between the proportion of cell types present and the individual organoid of origin (n = 3 single organoids per set, except for PGP1 replicate 2 at 6 months, for which n = 2, for a total of 83 organoids). Most replicates, across all ages and cell lines, had AMI scores comparable to AMI scores for two published endogenous human cortex datasets (Polioudakis *et al*., 2019; Trevino *et al*., 2021) and a newly-generated fetal dataset (see below, Figures 1E and S1B-I). This indicates that for each timepoint, organoid-to-organoid variation in the proportion of the different cell types is at the same low level as that of endogenous brains.

To begin to understand the regulatory logic that underlies this cellular diversification, we investigated gene-regulatory changes across organoid development. We generated single-cell Assay for Transposase Accessible Chromatin Using Sequencing (scATAC-seq) (Corces *et al*., 2017; Satpathy *et al*., 2019) profiles of 38,017 nuclei, combining our previously published dataset (Paulsen *et al*., 2022) with an additional 11,551 newly-profiled cells from 1, 3, and 6 month organoids, spanning amplification of neural progenitors, the peak of excitatory neuron diversity, and the emergence of astroglia and interneurons, respectively (Figure 1C and S3, n = 3 individual organoids per timepoint). To classify cell types, we integrated the expression profiles from scRNA-seq with inferred gene activity scores from scATAC-seq clusters by label transfer from scRNA-seq (Figure S3A-F), and refined assignments by looking at genes near differentially accessible regions (DARs). We provide the full epigenomic data for further exploration at https://singlecell.broadinstitute.org/single_cell/study/SCP1756.

Overall, we found good agreement between cell types as defined by scRNA-seq and scATAC-seq (Figures 1C, S3D-F). Moreover, AMI scores indicated that epigenetic variability between individual organoids falls into the same low range as transcriptional variability (Figure 1F).

We found that cell type-specificity of accessible chromatin increased with time, consistent with cell differentiation. At 1 and 3 months, approximately 4.5% of peaks were cell type-specific (i.e., called as differentially accessible in at least one cell type), which increased to 12.5% at 6 months (Figure S3G), suggesting increased specialization of epigenetic state over time.

To further investigate the relationship between RNA profiles and underlying chromatin organization in organoids, we also applied the multimodal single-cell assay SHARE-seq, which captures RNA-seq and ATAC-seq profiles simultaneously from the same single cell (Ma *et al*., 2020). We used SHARE-seq to profile a total of 42,810 cells across 4 timepoints, from 23 days to 3 months. Because the non-cortical cell types present at 23 days may have more distinct epigenomic profiles that might not be fully captured in the relatively low number of single cells at this timepoint (1,281 scATAC-seq profiles), we also performed bulk ATAC-seq on organoids at 23 days to generate a comprehensive reference peak set for analysis (Methods). These data identified the same cell types at each timepoint as the other modalities (Figure 1D), and showed the same high organoid-to-organoid reproducibility (Figure 1E).

Finally, to investigate the relationship between gene expression and the spatial position of different cell types in organoids, we applied spatial transcriptomics using Slide-seqV2 (Stickels *et al*., 2021) at 1, 2, and 3 months *in vitro* (Figures 2 and S4), spanning the emergence and expansion of excitatory neurons, which achieve their highest diversity in organoids at 3 months *in vitro*. Due to the size of the beads (10 μm), each data point may represent transcripts from multiple closely-adjacent cells. To resolve cell types from the per-bead spatial measurements, we applied RCTD (Robust Cell Type Decomposition; (Cable *et al*., 2021)) to transfer cell type annotations from our scRNA-seq data to the Slide-seqV2 data. In parallel, we also plotted the expression of the 50 most significantly differentially expressed genes (DEGs) in the scRNA-seq signatures (Table S1-3) for each cell type (Figure 2, Figure S4).

**Figure 2.**
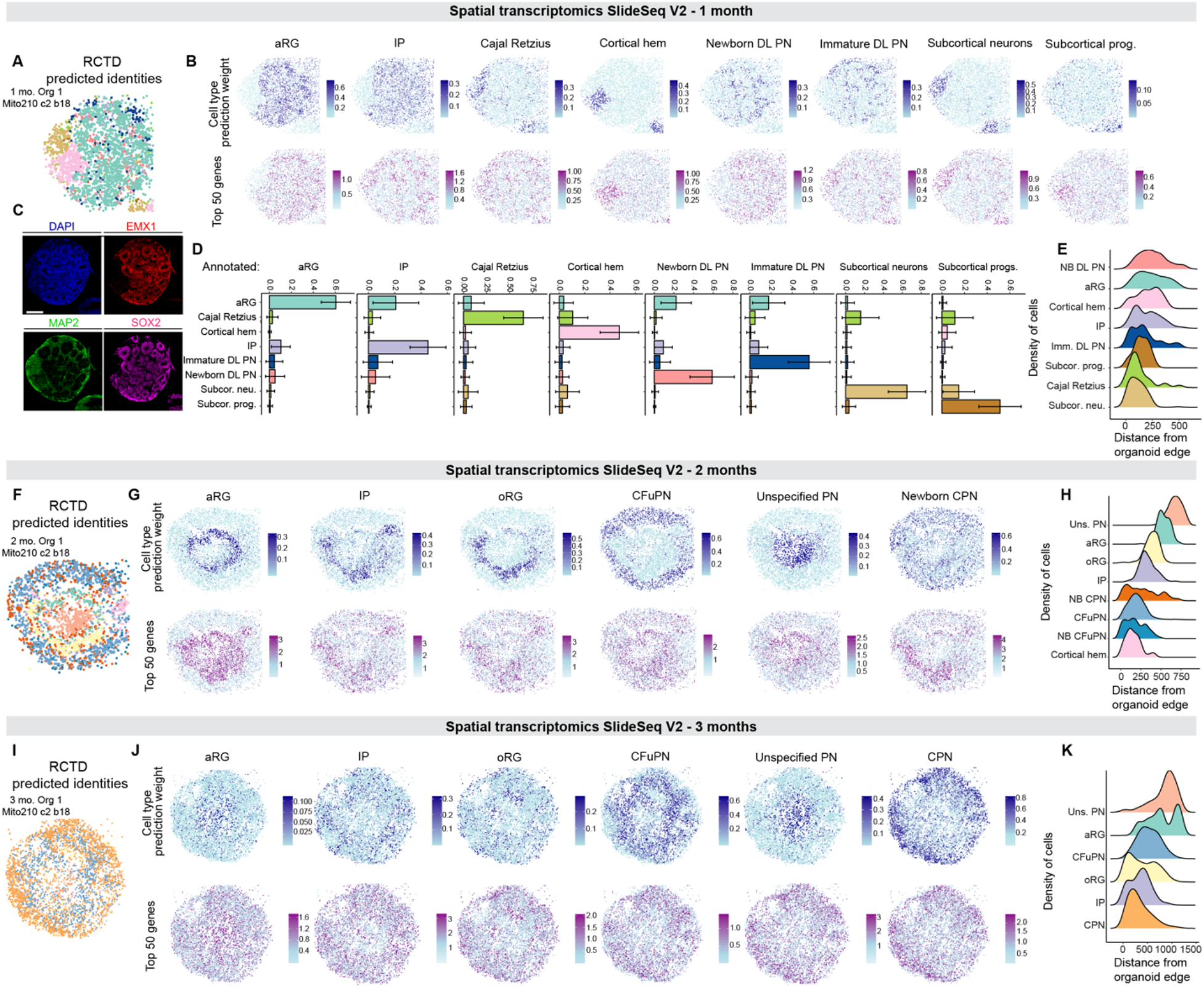
Spatial transcriptomic landscape of developing cortical organoids. (A) Spatial plot of Slide-seqV2 data from 1-month Organoid 1, colored by RCTD-assigned cell type. (B) Spatial plots showing Robust Cell Type Decomposition (RCTD) prediction weights (top row) and the summed, normalized expression of the top 50 marker genes (bottom row) for each cell type in 1-month Organoid 1. (C) Immunohistochemistry for neuronal (MAP2) and dorsal forebrain progenitor (EMX1, SOX2) cell types in 1-month Organoid 1, in a section adjacent (posterior) to the one used for Slide-seqV2. Scale bar 200 μm. (D) The mean ± standard deviation of the cell type weights given by RCTD, for beads in 1-month Organoid 1 that were annotated as each cell type. (E) The distribution of each annotated cell type over the beads’ calculated distance from the edge of the organoid, ordered from top to bottom by median distance. (F-H) Spatial plots and distributions of cell types from 2-month Organoid 1, as in A, B, and E. (I-K) Spatial plots of distributions of cell types from 3-month Organoid 1, as in A, B, and E. Abbreviations: “c”, cell line clone; “b” differentiation batch; org., organoid; NB, newborn; DL for deep-layer; PN, projection neurons; Subcor., subcortical; prog., progenitors; neu., neurons; aRG, apical radial glia; oRG, outer radial glia; IP, intermediate progenitor; CPN, callosal projection neuron; CFuPN, corticofugal projection neuron; Uns., unspecified.

This dual approach revealed spatial patterning of cell types in 1-, 2-, and 3-month organoids. At 1 month, we found clusters of aRG, occupying both central and peripheral regions of the organoid (Figures 2A-E, S4A, and S4D, n = 4 organoids). Cajal Retzius cells, IP, newborn DL PN, and immature DL PN were superficially located with respect to aRG. Interestingly, cells of the cortical hem and subcortical regions largely did not intermix with cortical cell types, but rather formed patches located next to each other, suggesting that cells of different regional origin self-segregate in developing organoids. At 2 months, aRG formed a ring of rosette-like structures around a core which consisted primarily of the ‘unspecified PN’ population (Figures 2F-H, S4B, and S4E, n = 3 organoids). The remaining cortical cell types were located largely superficial to the aRG rosettes. Strikingly, oRG were superficially positioned with respect to aRG (Figures 2H and S4E), reflecting the relative location of these cells in endogenous tissue (Fietz *et al*., 2010; Hansen *et al*., 2010). By 3 months there was a loss of recognizable structures such as rosettes, consistent with other studies showing that as organoid development progresses, the rudimentary structure of the developing cortical wall is lost (Eiraku *et al*., 2008; Velasco *et al*., 2019; Qian *et al*., 2020) (Figures 2I-K, S4C, and S4F, n = 3 organoids). Some degree of organization could still be observed: for example, most CPN were superficial to CFuPN, reflecting their relative positioning *in vivo* (Figures 2K and S4F). The inner core of the organoids was mainly populated by ‘unspecified PN’ and aRG, similar to the 2-month organoids (Figures 2K and S4F). These data show dynamic changes and preferential positioning of cell types in cortical organoids that reflect processes observed *in vivo*.

Overall, these data provide a comprehensive multiomics molecular map of the development of human cortical organoids.

### Longitudinal fetal programs of cell identity acquisition are established in cortical organoids with cell-type and temporal specificity

Previous work has shown resemblance between cortical organoids and endogenous human fetal tissue (Camp *et al*., 2015; Velasco *et al*., 2019; Bhaduri *et al*., 2020; Gordon *et al*., 2021). However, it remains unexplored whether these similarities extend equally to all cell types and across all steps of development. For this purpose, we first sought to understand the dynamic molecular programs that define each cell type across organoid development. We determined gene expression signatures for all cortical cell types at each stage (Table S2-8, Figures 3A-B and S5), using a pseudo-bulk analysis with a model that controls for variability between organoids (see Methods). We confirmed that the most significant DEGs in these signatures reflected developmental-stage and cell-type-appropriate expression of known endogenous markers (see Supplementary Text and Figure S5). Importantly, this analysis showed that the transcriptional state of individual cell types was highly reproducible across organoids (Figures 3A-B, S5; one-way ANOVA, p > 0.05).

**Figure 3.**
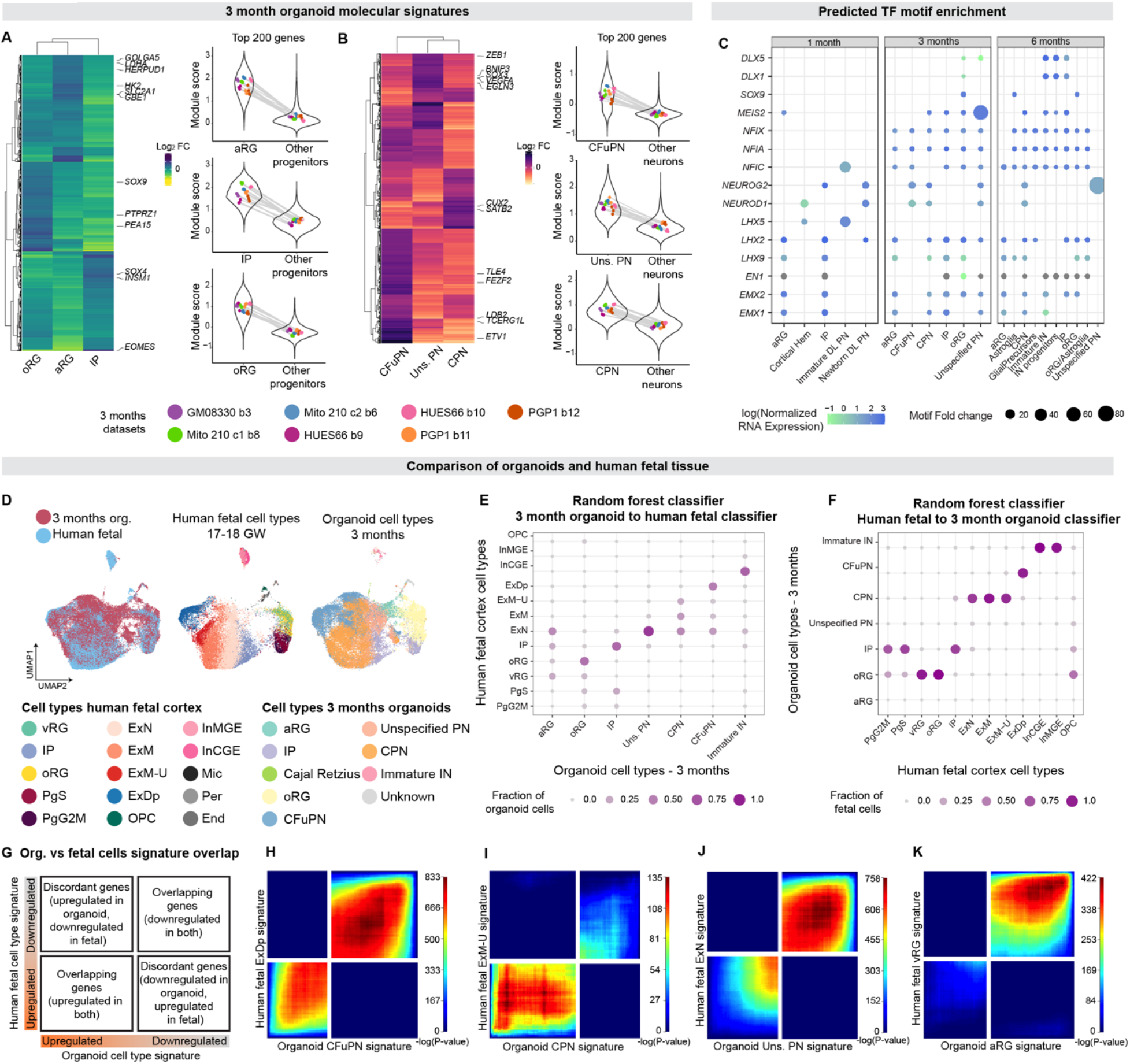
Longitudinal fetal programs of cell identity acquisition are established in cortical organoids with cell type specificity. (A, B) Left, heatmaps comparing the upregulated genes in each progenitor (A) or neuronal (B) population (showing genes with adjusted p value < 0.0015, log_2_ fold change > 1.5, Table S2-8). Right, gene set expression of the top 200 genes in the molecular signatures of each cell type. Violin plots show the module score distribution across single cells, points show the mean module score for each individual organoid, with lines connecting points representing cells from the same organoid. The means within each distribution (separately, within the cell type for that signature, left, or the background cells, right, for each plot) showed no significant difference across individual organoids in all cases (one-way ANOVA, p > 0.05). (C) Transcription factor (TF) motifs enriched (p < 1e-10) in peaks with increased accessibility (Bonferroni adjusted p < 0.1) in each cell type, per timepoint. Points are sized by fold change enrichment of the motif in accessible peaks versus GC-content-matched background regions. Points are colored by normalized mRNA expression for each TF in the corresponding cell type, from corresponding scRNA-seq. Gray points indicate zero expression. (D) UMAPs representing data from all 3-month organoids, integrated via Harmony batch correction with human fetal scRNA-seq data (Polioudakis *et al*., 2019). Left, cells are colored by dataset. Middle, fetal cells are colored by cell type as assigned in (Polioudakis *et al*., 2019). Right, organoid cells are colored by cell type. (E) Assignments of 3-month organoid cells to human cortical fetal cell types, as classified by random forest. Points are sized and colored by the fraction of each organoid cell type assigned to the fetal cell types. (F) Assignments of human fetal cells to 3-month organoid cell types, as classified by random forest. Points are sized and colored by the fraction of each human fetal cell type assigned to the organoid cell types. (G) Schematic explaining the rank-rank hypergeometric overlap (RRHO2) output plot. Genes from each signature are ordered from most upregulated to most downregulated, with the most upregulated gene of each signature in the lower left corner. (H-K) RRHO2 plots from comparing the signatures of organoid CFuPN to fetal ExDp (H), organoid CPN to fetal ExM-U (I), organoid unspecified PN to fetal ExN (J), and organoid aRG to fetal vRG (K). Points in the plot are colored by the p value of hypergeometric tests measuring the significance of overlap of gene lists up to that point in the lists. Cell type abbreviations as in previous figures, with the addition of those in (Polioudakis *et al*., 2019): vRG, ventricular radial glia; PgS, progenitors in S phase; PgG2M, progenitors in G2M phase; ExN, migrating excitatory neurons; ExM, maturing excitatory neurons; ExM-U, maturing excitatory neurons-upper layer enriched; ExDp, deep layer excitatory neurons; OPC oligodendrocyte precursors; InMGE, interneurons from the medial ganglionic eminence; InCGE, interneurons from the caudal ganglionic eminence; Mic, microglia; Per, pericyte; End, endothelial.

To identify the regulatory programs mediating cell diversification, we leveraged our chromatin landscape atlas to define the epigenetic signatures of each cell type, and identify transcription factors (TFs) associated with each cell type using motif enrichment analysis (see Methods; (Heinz *et al*., 2010)). We found motif enrichment and gene expression of known developmental-stage and cell-type appropriate TFs for different cell populations (Figure 3C, Supplementary Text, and Table S9). To examine how the expression of a TF within a single cell was associated with the chromatin accessibility of its motif, we analyzed the SHARE-seq timecourse spanning the day 23 to 3-month timepoints. We found that for many of these TFs, such as *EMX1*, *LHX2*, and *NFIA* (Figure S3I), gene expression correlated positively with motif accessibility across the genome, suggesting that these TFs are activators that promote opening of chromatin at their binding sites.

Together, this work provides longitudinal transcriptional and epigenetic signatures of cell types across multiple stages of human cortical organoid development, and demonstrates that cell diversification can occur *in vitro* through robust and reproducible developmental programs.

Having defined the molecular signatures identifying cortical cell types in our human cortical organoids, we sought to determine how well these signatures corresponded to the ones defining cortical cell types in the endogenous human fetal cortex. We integrated our organoid data with two published scRNA-seq datasets (Figure 3D, (Polioudakis *et al*., 2019; Trevino *et al*., 2021)) as well as with 60,806 newly profiled single nuclei of the human fetal somatosensory cortex at PCW 14, 15, 16, and 18 generated in this study (Figure S6). We used two separate, complementary approaches to assess similarity: **1)** A classification-based approach, where we trained a random forest classifier (Shekhar *et al*., 2016) on fetal cortical cell types, and applied it to assign organoid cortical cells to the endogenous human cell types, and vice versa; and **2)** a pairwise comparison of DEGs, using molecular signatures identified for each cortical cell type in organoid and fetal cells separately (see Methods). Both approaches indicated high agreement between organoid and fetal cortical cell types.

For the classifier-based approach, we integrated our organoid data (at each stage from 1 to 6 months) with two published scRNA-seq datasets (Polioudakis *et al*., 2019; Trevino *et al*., 2021). We found that the vast majority of organoid cells at each stage were predominantly assigned to the corresponding endogenous cell types by the classifier (Figures 3E, and S7A-B). In parallel, we trained the classifier on the organoid cells from each timepoint (1 to 6 months) and applied it to classify fetal cells; this analysis can reveal whether organoid cell populations are separated by factors that differ from those distinguishing endogenous populations. In this approach as well, most fetal cell types were assigned to their expected counterparts in organoids (Figure 3F and S7C-D). For instance, with respect to the classifier trained with cells from 3 months organoids, the timepoint with the highest excitatory neuron diversity in organoids, we observed that fetal deep-layer neurons were matched almost exclusively to organoid CFuPN; other fetal excitatory neurons (i.e., migrating, maturing, and upper-layer enriched in (Polioudakis *et al*., 2019), Glutamatergic neurons 1-8, in (Trevino *et al*., 2021)) to CPN; fetal IP/nIPC to organoid IP; and both subtypes of fetal interneurons (InMGE/MGE-IN and InCGE/CGE-IN) to organoid IN. Fetal cycling progenitors from both datasets were split between organoid oRG and IP, while fetal oRG were assigned almost exclusively to organoid oRG. While the vast majority of cell types were assigned to their expected endogenous counterparts, very few fetal cells were assigned to the aRG and ‘unspecified PN’ organoid cells (Figure 3F), suggesting that these populations are less closely related to endogenous cell types.

Analysis with classifiers trained on 2, 4, 5, or 6-month organoid cells yielded similar results (Figure S7C-D). Notably, however, analysis with classifiers trained on 1 or 1.5-month organoid cells showed multiple fetal progenitor cell types assigned to organoid aRG (Figure S7C-D), likely pointing to stage-specific differences in the transcriptional profile of the organoid aRG populations.

As a second, independent approach, we compared the molecular signatures for all pairs of fetal and organoid cortical cell populations (1 to 6 months), using rank-rank hypergeometric overlap (RRHO2), a method that is designed to compare ranked DEGs lists between two independent gene profiling experiments (Plaisier *et al*., 2010; Cahill *et al*., 2018). Each such pairwise comparison results in a plot showing the strength and pattern of the correlation between the DEGs in each cell type, highlighting both concordant and discordant patterns for up- and down-regulated genes (Figure 3G). As in the classifier analysis, most organoid cell types showed a high agreement of gene expression signatures with the corresponding endogenous fetal cell types (Figures 3H-K, S8-14). For example, at 3 months, organoid CFuPN and CPN strongly agreed with fetal ExDp (deep layer) and ExM-U (upper-layer enriched) neurons in the (Polioudakis *et al*., 2019) dataset, and with SP (which express CFuPN markers) and GluN 2-4 and 7-8 (expressing markers of CPN) clusters of the (Trevino *et al*., 2021) dataset, respectively (Figures 3H-I, the bottom left quadrant of each plot, and Figure S11). Similarly, organoid oRG and IP signatures matched with the endogenous oRG and IP signatures from (Polioudakis *et al*., 2019), with RG and nIPC signatures from (Trevino *et al*., 2021), as well as with oRG and IP from our newly-generated fetal dataset (Figures S8-S14).

Importantly, we found that the molecular signature of the ‘unspecified PN’ population of organoids, while most closely matching the fetal migrating ExNs from (Polioudakis *et al*., 2019), GluN 1 and 5 clusters from (Trevino *et al*., 2021), or CPN from our fetal dataset, showed a comparatively weak agreement overall, driven more by shared down-regulated genes (Figure 3J and S10-S14). Similarly, the signature of organoid aRG did not show a strong correlation to any fetal signature at any timepoint from 2 months *in vitro* onwards (Figure 3K and Figure S10-S14). Thus, although these two cell types showed some overall transcriptional resemblance to endogenous cells (as they could be assigned to appropriate cell types via the random forest classifier trained on fetal cells, Figure 3E), their transcriptional signatures had comparatively weak matching to cell type-specific fetal signatures from this gestational age range, compared to all other classes of organoid cells.

Overall, using two independent methods we find that the vast majority of organoid cell types closely transcriptionally resemble cells of the endogenous fetal cortex, with only two notable exceptions, aRG and a type of mis-specified PN that does not seem to exist *in vivo*.

We next sought to assess the concordance of the epigenomic landscape of cell types in brain organoids to that of their endogenous counterparts. We compared scATAC-seq data from 1, 3, and 6 months organoids to a recently published dataset of scATAC-seq in human cortex at PCW 16-24 (Trevino *et al*., 2021). For each of the three organoid timepoints, the fetal and organoid datasets were projected to a shared peak set, and molecular signatures were calculated and compared using RRHO2, as for RNA-seq (see Methods). Again, we found that corresponding cell types had highly similar cell type signatures (Figure S7E-G, Figure S15, and Supplementary Text). Notably, we found substantial overlap between the scATAC-seq peaks called in each dataset, with higher concordance at the older organoid timepoints: 73% of peaks called in 1-month organoids overlap with peaks called in fetal cells, while 88% of peaks called in 3-month organoids and 85% of peaks called in 6-month organoids overlap with peaks called in fetal cells. This suggests that 1-month organoids probably represent an earlier developmental stage than the PCW 16-24 cortex profiled in the (Trevino *et al*., 2021) dataset. In addition, certain organoid cell types had a particularly high proportion of overlapping peaks with their corresponding cell types in the (Trevino *et al*., 2021) dataset: 94.8% of peaks called in aRG at 6 months overlap peaks called in fetal RG, and 94.6% of peaks called in CPN at 3 months overlap with the fetal glutamatergic neuron populations (largely CPN). Overall, these data show that organoid cell types have highly similar epigenomic profiles to their endogenous fetal counterparts.

Finally, we applied the epigenomic atlas to query cell type-specific dynamics of gene regulatory elements across organoid development. As a test case to compare mechanisms of gene regulation in organoids compared to endogenous processes, we examined regulation of *FEZF2,* a TF that regulates the fate specification of a subset of layer 5 CFuPN (Chen, Schaevitz and McConnell, 2005; Molyneaux *et al*., 2005; Eckler *et al*., 2014), previously reported to have a CFuPN-specific cis-regulatory interaction with a neighboring highly-conserved enhancer (Eckler *et al*., 2014; Markenscoff-Papadimitriou *et al*., 2020). Our data showed that the cell type-specific epigenetic regulation of this gene in cortical organoids is similar to that observed *in vivo* (Figure S7H-L and Supplementary text).

Overall, our analysis demonstrates that the majority of organoid cell types closely replicate the cell type-specific signatures and epigenetic states of the corresponding human fetal cortical cell types, indicating a high degree of fidelity in cell identity.

### Human cortical organoid cell type identity is largely unaffected by metabolic state

To examine biological processes unfolding as development progresses in organoids, we applied Weighted Gene Correlation Network Analysis (WGCNA; (Langfelder and Horvath, 2008)) to identify modules of genes whose expression co-varies across organoid cells at each age (1 to 6 months). Overall, modules reflected the unfolding of appropriate developmental programs for each stage (Figure S16, Table S11). However, modules including glycolysis-associated genes (e.g., *ENO1*, *PGK1,* Table S11) were detected starting at 1.5 months *in vitro*, and were specifically enriched in the two populations of cells with the weakest resemblance to fetal cells, namely aRG and ‘unspecified PN’, from 2 months *in vitro* onwards (Figures 4A-B). At 1 month, no module representing glycolysis was found (Figure S16, Table S11), but at 1.5 months a module of glycolysis genes was enriched in subsets of the aRG and deep-layer PN populations, suggesting that the glycolysis metabolic signature emerges at this stage (Figure S16B).

**Figure 4.**
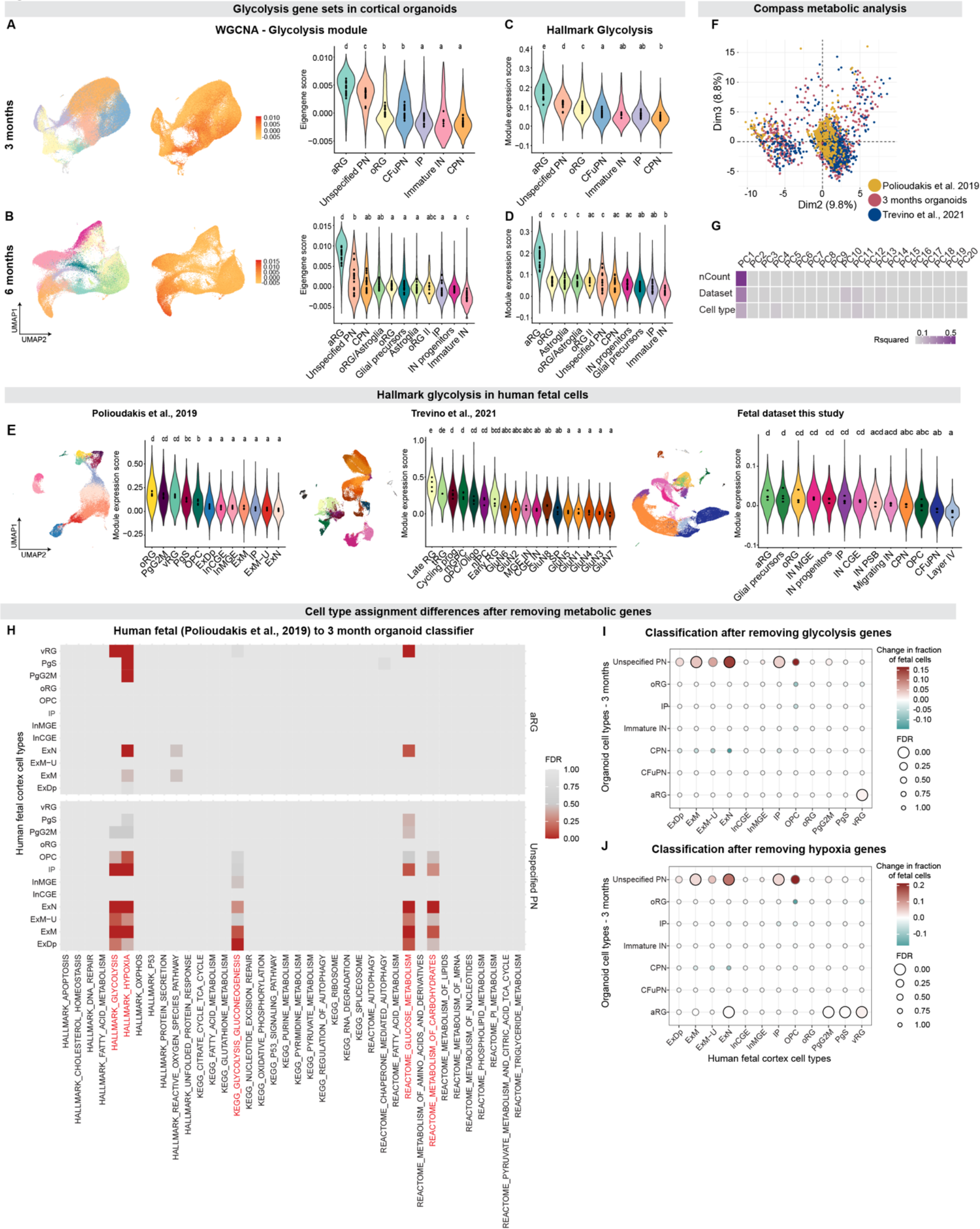
Human cortical organoid cell type identity is largely unaffected by metabolic state. (A-B) Enrichment of a Weighted Gene Correlation Network Analysis (WGCNA) gene module containing glycolysis genes in 3- and 6-month organoids. Left, UMAP of cells from all 3-month (A) and 6-month (B) organoids, downsampled to have an equal number of cells per organoid. Cells are colored according to cell type (left) and to eigengene score for the “glycolysis” module (middle). Right, violin plots showing the distribution of eigengene scores for this module across cell types. Points indicate average scores for each individual organoid. Letters above violin plots indicate the results of a one-way ANOVA followed by pairwise TukeyHSD comparisons; cell types with the same letter indicate there is no significant difference between organoid averages of those cell types. (C-D) Violin plots showing the distribution of module scores for the MSigDB Hallmark Glycolysis gene set across cell types at 3 months (C) and 6 months (D), in the same downsampled data. Points and letters as in A. (E) UMAP of the human fetal cells (Right: (Polioudakis *et al*., 2019), middle: (Trevino *et al*., 2021), right: novel fetal dataset) colored by their assigned cell type (top), and violin plot showing the distribution of module scores for the MSigDB Hallmark Glycolysis gene set across cell types (bottom). Points and letters as in A. (F) Principal component analysis (PCA) of the Compass matrix of metabolic reaction potential activity scores. Cells are colored by the dataset that they are derived from. (G) Correlation of the top 20 principal components of the Compass PCA with three metadata values. Tiles colored by R squared values from linear regressions of the principal component loadings with each metadata value separately: nCount (the number of UMIs per cell), the dataset that each cell is derived from, and the cell type label of each cell. (H) Significance (False Discovery Rate by Benjamini-Yekutieli) of the increase in (Polioudakis *et al*., 2019) fetal cell assignments to the 3-month organoid aRG label (top) and unspecified PN cell types (bottom) after removing genes from the model in each of the 38 gene sets listed along x axis. (I-J) Change in the assignments of (Polioudakis *et al*., 2019) fetal cell types to 3-month organoid cell types, after removing genes from the model that belong to the MSigDB Hallmark Glycolysis gene set (I) and Hypoxia gene set (J). Points are colored by the change in the fraction of cells assigned to each label by the model with removed genes, compared to the full model. Points are sized by the significance of the increase in that fraction (FDR by Benjamini-Yekutieli), compared to a background distribution of 10,000 models trained after removing random expression-matched gene lists. Points with FDR < 0.05 are outlined in black. As in the text, changes were considered significant if they showed an FDR<0.05 and changed at least 1% of assigned cells. Cell type abbreviations as in previous figures, with the addition of those in (Trevino *et al*., 2021): tRG: truncated radial glia; mGPC, multipotent glial progenitor cell; OPC/Oligo, oligodendrocyte progenitor cell/oligodendrocyte; nIPC, neuronal intermediate progenitor cell; GluN, glutamatergic neuron; CGE IN, caudal ganglionic eminence interneuron; MGE IN, medial ganglionic eminence interneuron; SP, subplate.

These findings prompted us to examine cell type-specific variation in metabolic processes implicated in organoid biology (Pollen *et al*., 2019; Bhaduri *et al*., 2020; Qian *et al*., 2020; Tanaka *et al*., 2020) using curated gene sets from the Molecular Signatures Database Hallmark collection (MsigDB; (Liberzon *et al*., 2011)). We found that the MSigDB glycolysis gene set was enriched in ‘unspecified PN’ and in aRG, along with other progenitor types, at multiple stages of organoid development (Figure 4C-D and S17). This is comparable, however, to a mild enrichment of this gene set observed in a subset of endogenous human fetal progenitors (oRG, vRG, and PgG2M from (Polioudakis *et al*., 2019); Late RG, tRG, Cycling progenitors, and mGPC from (Trevino *et al*., 2021); aRG, Oligodendrocyte precursors, and oRG from the novel fetal dataset); Figure 4E and S18A). The MSigDB hypoxia gene set was also enriched in both organoid aRG (2 months onwards) and in endogenous fetal oRG and vRG (Polioudakis *et al*., 2019), fetal Late RG, tRG, and mGPC (Trevino *et al*., 2021) and fetal Glial precursors, aRG and oRG (novel fetal dataset) (Figure S18B), consistent with the high molecular cross-talk and genes shared between the hypoxia and glycolysis pathways (Semenza, 2012; Zheng *et al*., 2016). We also confirmed that upregulation of metabolic states in organoid cells could be detected on an epigenetic level in the SHARE-seq data (Table S14, Figure S18D-E, and Supplementary Text). The data indicate that metabolic states observed during normal development of cortical progenitors are also observed in the corresponding cell types in organoids.

To understand whether these metabolic differences depended on the culture protocol employed, we profiled 69,333 cells from a different organoid model, whole-brain organoids (Quadrato *et al*., 2017), at 3.5 months *in vitro*, and found a similar specific enrichment of glycolysis and hypoxia gene sets in aRG and unspecified PN cell populations (Figure S19 and Supplementary Text), showing that multiple organoid models have similarly restricted differences in metabolic gene expression across similar cell types.

In contrast, while oxidative phosphorylation has been reported to be abnormal in organoids (Bhaduri *et al*., 2020), the MSigDB oxidative phosphorylation signature did not show strong enrichment in any particular cortical cell type in either of our organoid models or in fetal tissue (Figure S17C, S18C, S19G and Supplementary Text).

Collectively, these data demonstrate that neural progenitors, especially aRG, in both organoids and endogenous fetal tissue show an enrichment of glycolysis and hypoxia genes, possibly due to the previously reported energetic requirements of these cells (Khacho *et al*., 2016; Zheng *et al*., 2016; Namba *et al*., 2020).

Given these findings, we sought to examine the degree to which organoids differed from endogenous fetal cells in expression of metabolic genes. For this, we applied three complementary approaches: Compass, an algorithm that uses flux balance analysis to model the metabolic state of single cells from scRNA-seq data (Wagner *et al*., 2021); RRHO2, to identify processes enriched among the genes up-regulated in organoids but downregulated in fetal cells; and differential expression of MSigDB gene sets for a broad range of metabolic processes (Methods and Supplementary Text). Across all three analyses, organoids and fetal cells showed overall similar results, with only a few metabolic pathways, in particular glycolysis and oxidative phosphorylation, being enriched in organoids (Figures S18F-H and S20). Notably, the Compass results showed that organoid and endogenous fetal cells clustered together well, and could not be discriminated based on their overall metabolic flux (Figure 4F-G).

The enrichment of metabolic genes in the aRG and ‘unspecified PN’ populations combined with the comparatively weak transcriptional matching between these two organoid cell types and endogenous fetal cells led us to wonder whether divergent metabolic processes could be interfering with the alignment of organoid cell types to their endogenous counterparts, as has been suggested for other cortical organoid models (Bhaduri *et al*., 2020). We therefore examined the effect of removing the 38 MSigDB metabolic gene sets described above on cell-type classification. For this, we systematically removed each metabolic gene set from the variable genes used to train the random forest classifiers for both fetal and organoid cells, and assessed the change in the proportion of fetal cells assigned to each fetal and organoid cell type, respectively, for multiple organoid timepoints and fetal datasets. Changes to cell type assignments were considered significant if >1% of cells were re-assigned to a new label, and if the proportion of cells reassigned was higher than expected by chance (FDR<0.05). The classifier trained with fetal cells did not show any significant change in the number of cells assigned to each fetal cell type (Table S12), consistent with the good matching of organoid cells to fetal ones observed already with the full set of variable genes. Similarly, for the organoid-trained classifier, for the vast majority of processes, removing the gene sets did not significantly change the number of cells assigned to each organoid cell type (Figures 4H-J and S21A-C, Table S12). Only two categories of metabolic processes affected classification: in the classifier trained with 3 month organoid cells, removing gene sets related to glucose metabolism and hypoxia (Hallmark Glycolysis, Hallmark Hypoxia, KEGG Glycolysis and Gluconeogenesis, Reactome Glucose Metabolism, and Reactome Metabolism of Carbohydrates) significantly increased the number of fetal cells assigned to the ‘unspecified PN’ and aRG cell types (FDR < 0.0001, Figures 4H-J), and in the classifier trained with 6-month organoids cells, removing genes related to the Hallmark Glycolysis gene set increased the proportion of cells assigned to aRG in the (Polioudakis *et al*., 2019) dataset (FDR < 0.0001, Figure S21A, D, G, Table S12), while no significant changes (FDR <0.05, >1% of cells re-assigned) were observed in assignments of the (Trevino *et al*., 2021) dataset (Figure S21B, C,E-F, H-I). No other cell type showed an increase in assignments at any of these timepoints. Importantly, genes related to other fundamental subcellular processes such as apoptosis and oxidative phosphorylation did not significantly change the assignment of any fetal cell type to organoid cell types (Figure 4H, Figure S21A-C, Table S12).

Overall, our analysis shows that specification of the vast majority of the cortical cell types generated in the present organoid model is not affected by diverging metabolic states, which affect the identity of only two cell types (aRG and ‘unspecified PN’). Furthermore, only glycolysis and hypoxia affected assignment in these two cell types, pointing at both cell-type- and pathway-specific associations between cell identity acquisition and metabolic susceptibility.

### Metabolically-compromised cells reside in a restricted and central region of human cortical organoids

To understand the relationship between the topographical location of organoid cells and the expression of metabolic pathways, we first performed immunohistochemistry for SLC16A3, a member of the Hallmark glycolysis gene set and also part of the cell-type signatures for organoid aRG and ‘unspecified PN’, and GORASP2, previously reported as a marker for cellular stress in organoids (Bhaduri *et al*., 2020). At 1 month, both proteins were expressed in SOX2^+^ neural progenitors (aRG) located in rosette-like structures (Figure S22A-B). However, at later stages (2 and 3 months *in vitro*), these proteins did not co-localize with canonical markers of RG, CFuPN, or CPN (Figure S22C-F), confirming that glycolysis genes are excluded from major cortical cell types. Of note, cells expressing these proteins were predominantly located in the inner core of the organoids.

We therefore leveraged our spatial transcriptomics developmental atlas to investigate whether there was an association between topographical location and metabolic state of cortical organoid cells. The inner regions of both 2- and 3-month organoids were predominantly composed of aRG and unspecified PN (Figures 2F-K, S4B-C, E-F, and 5A), the two populations whose metabolic state affected assignment of cell identity. Notably, aRG became more centrally located between 1 and 2 months, consistent with our finding that aRG identity becomes affected starting at 1.5 months (Figures S9 and S17), presumably as organoid size begins to create a more hypoxic environment at the tissue core.

To test for topographic association of metabolic pathways, we plotted expression of the 38 MSigDB metabolic pathway gene sets in the spatial transcriptomic data. Most pathways (e.g., apoptosis) remained constant across the organoid diameter (Figure 5C-D and S22G-I). However, pathways associated with hypoxia and glycolysis, which we previously identified as altering aRG and unspecified PN identity, were enriched in cells towards the center of the organoid (Figures 5D and S22G-I). Besides the aRG and unspecified PN populations, cells located in the inner regions of the organoids showed higher expression of glycolysis and hypoxia genes, independent of cell identity (Figure 5E and S22J-L). We confirmed these results with the hypoxia detection reagent pimonidazole, which also showed signal restricted to the organoid core from 2 months *in vitro* onwards (Figure S23).

**Figure 5.**
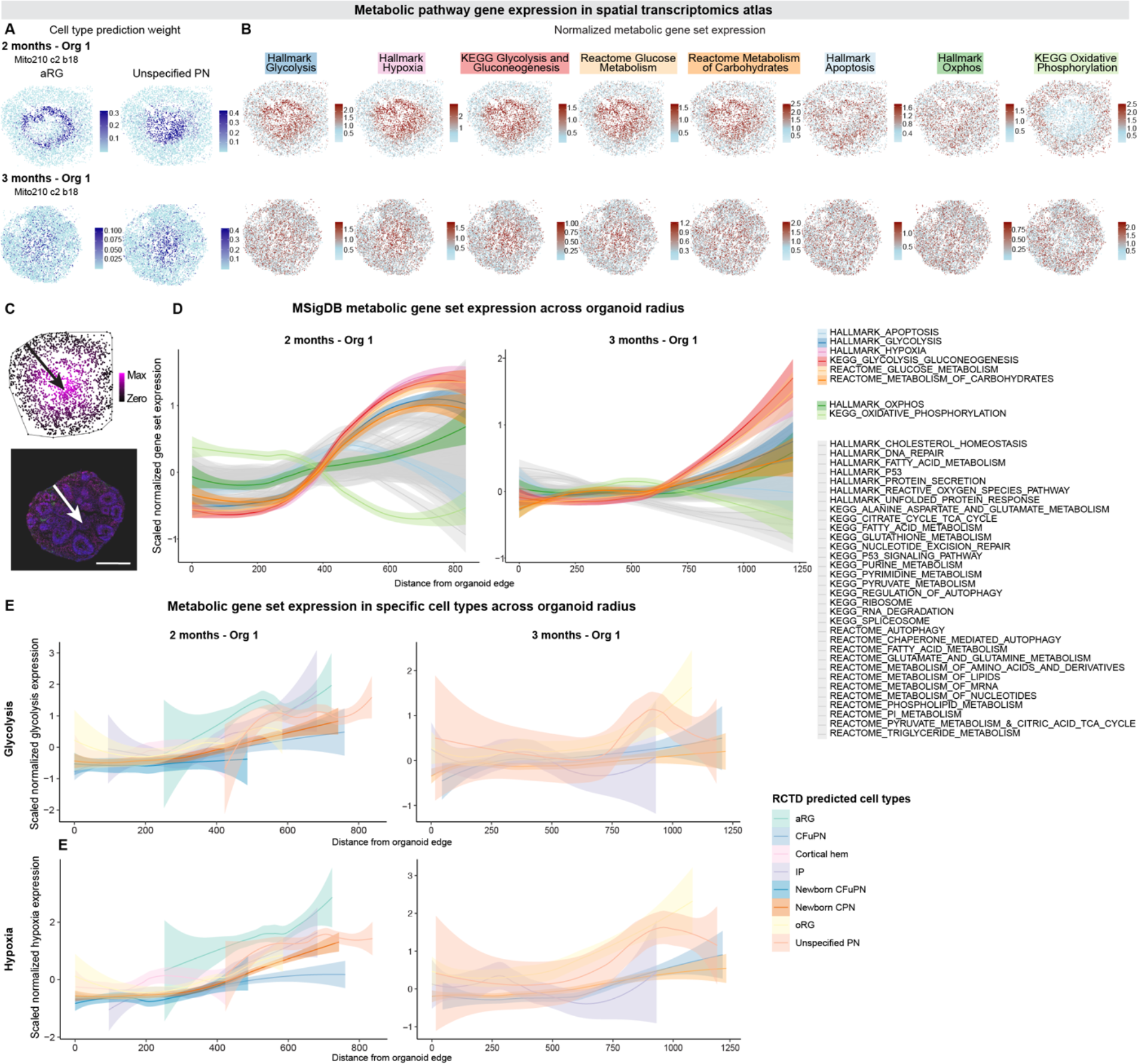
Metabolically-compromised cells reside in a restricted and central region of human cortical organoids. (A) Spatial plots colored by RCTD prediction weights for aRG (left) and unspecified PN (right), for 2-months Organoid 1 (top) and 3-months Organoid 1 (bottom), repeated from Figures 3G and 3J for ease of comparison. (B) Spatial plots showing the summed, normalized expression of genes in the indicated MSigDB gene sets for 2-months Organoid 1 (top) and 3-months Organoid 1 (bottom). (C) Schematic showing how beads’ distance from the edge of the organoid was calculated. Top, Slide-seqV2 data from 2-months Organoid 1, with beads colored by distance from organoid edge. Line shows convex hull around organoid. Bottom, IHC of a 1.5-month organoid, showing DAPI (blue), *CTIP2* (magenta), *HOPX* (green), and *SOX2* (red). Scale bar 200 μm. Arrows point from edge of organoid to points most distant from the edge. (D) Distribution of the beads’ scaled expression for each gene set compared to their distance from the edge of the organoid. Solid line shows the smoothed conditional mean values across beads, with transparent backgrounds showing the 95% confidence interval. Distributions are shown separately for 2-month Organoid 1 (left) and 3-month Organoid 1 (right). (E) Distributions as in C for the Hallmark Glycolysis (top) and Hallmark Hypoxia (bottom) gene sets, graphed separately for beads assigned to each cell type. Cell types assigned to more than 8 beads per organoid are shown. Distributions are shown separately for 2-month Organoid 1 (left) and 3-month Organoid 1 (right). Cell type abbreviations as in previous figures.

Altogether, these data show that only cells located within inner regions of cortical organoids (primarily aRG and ‘unspecified PN’) become metabolically challenged and express higher levels of specific metabolic gene sets, i.e., glycolysis and hypoxia, compared to cells populating more superficial position (the majority of cell types). The data point at position-specific and metabolic pathway-specific effects that are restricted to cells in the organoid core, rather than broad metabolic susceptibility of cells generated in organoids compared to endogenous cortical tissue.

### Molecular logic of development of individual human cortical cell types

We next sought to apply our molecular map of organoid development to understand how cellular lineages are established in the human cortex, and the transcriptional events associated with lineage decisions. We first used the longitudinal transcriptional dataset to infer the cells’ developmental trajectories, focusing on the 459,711 cortical cells (excluding unspecified PN). To highlight relationships between cell types across time, we connected cell clusters from the same and consecutive timepoints by transcriptional similarity (see Methods; (Ko *et al*., 2020)) (Figures 6A and S24A). When visualized as a force-directed graph, aRG were connected to other progenitor types (IP, oRG), as well as to different types of excitatory neurons, both early- (Preplate/Subplate, newborn DL PN, Immature DL PN) and late-produced (CFuPN, CPN), consistent with the existence of both direct and indirect (i.e., through other progenitor cell types) neurogenesis in organoids. At 5 and 6 months, aRG were also connected to astroglia, oligodendrocyte precursors, and interneurons. The generation of these cell types occurs during perinatal (late) stages of cortical development *in vivo* (Kriegstein and Alvarez-Buylla, 2009; Obernier and Alvarez-Buylla, 2019). The majority of IPs were found within the excitatory neuron lineage (66% of first-neighbor clusters to IP clusters contain majority excitatory projection neurons or other IPs, whereas only 33% of all clusters belong to those types), in agreement with the commitment of this progenitor cell type to glutamatergic neuron production *in vivo* (Hevner, 2019).

**Figure 6.**
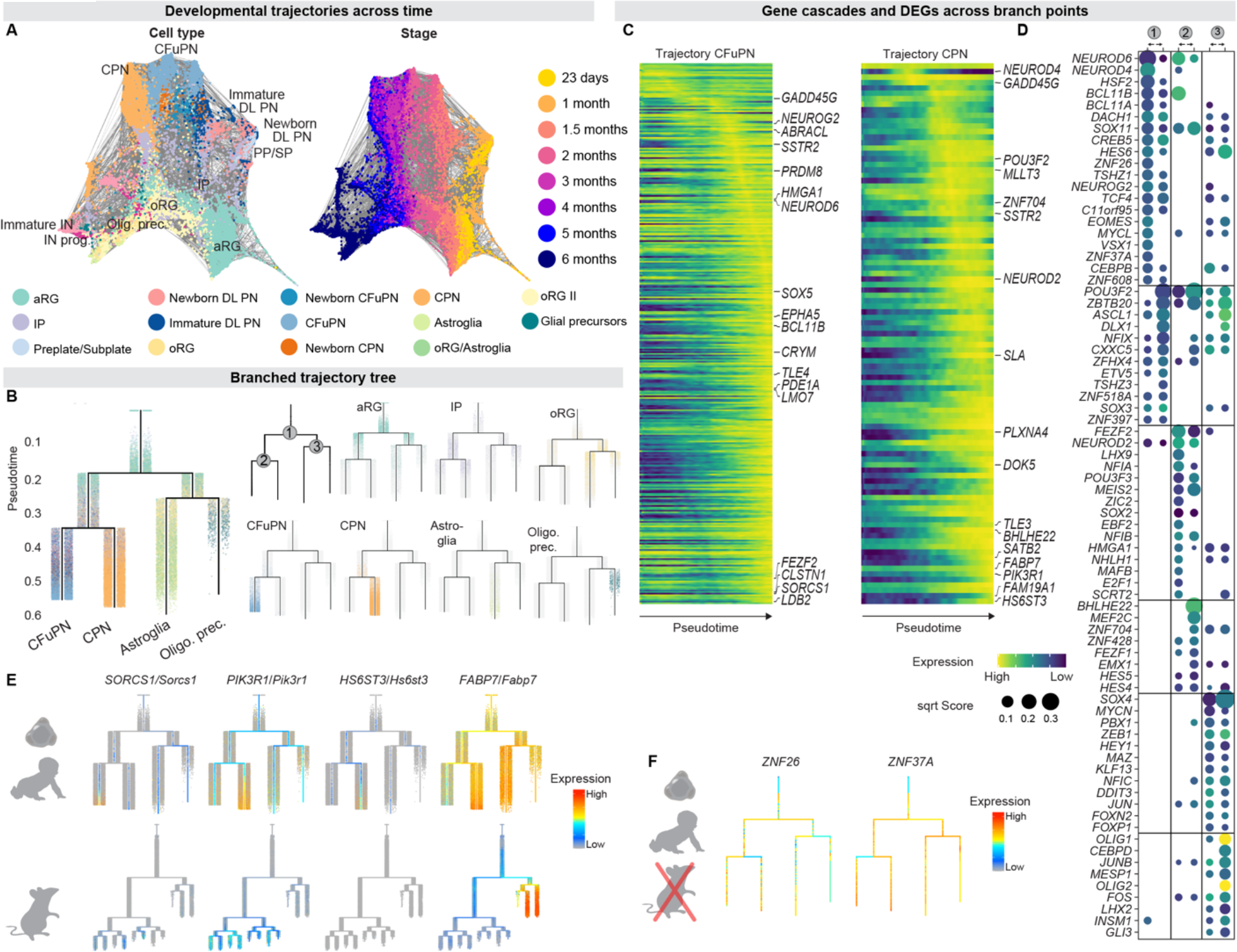
Molecular logic of development of individual human cortical cell types. (A) Integrated trajectory of scRNA-seq datasets across time. Weighted graph is created using the FLOWMAP algorithm and visualized using a force-directed layout. Cell clusters are colored by majority cell type (left, colors as in Figure 1) and by timepoint (right). Gray lines connect cell clusters with high transcriptional similarity. (B) URD branching tree. The root (top of the tree) was set to day 23 aRG, tips (terminal states at the bottom of the tree) are CFuPN, CPN, astroglia and oligodendrocyte precursors. Cells are colored by identity. Branch points are labeled 1-3. (C) Heatmap of lineage-specific gene cascades for CFuPN and CPN. Gene expression in each row is scaled to maximum observed expression. Genes are ordered on the basis of their onset and peak expression timings (Methods). Full cascade is shown in Table S15. (D) Top 20 transcription factors per branch predicted to be involved in cell type divergence. Points are sized by their score, defined as their feature importance in a gradient boosting-classifier distinguishing cells in each segment of the branch point, and are colored by their average expression in the corresponding cells (row-scaled, color). Branch points numbered as in B. DEGs: differentially expressed genes (E) Expression of human lineage-specific genes in human cortical organoid tree (top) and developing mouse cortex tree (bottom, (Di Bella *et al*., 2021)). (F) Expression of two human ZNF genes in the simplified branching tree, both of which have a high feature importance score in the first branch point. Cell type abbreviations as in previous figures.

oRG were connected to postmitotic neurons, in agreement with the neurogenic potential of this progenitor cell type (Fietz *et al*., 2010; Hansen *et al*., 2010). We also observed a connection between oRG and IPs (2.7% of IP neighbors are oRG), suggesting that oRG can give rise to IP in organoids, as described in the endogenous human and non-human primate cortex (Betizeau *et al*., 2013; Coquand *et al*., 2022). At late timepoints, oRG were also connected to interneuron progenitors and astroglia (Figure 6A). This is interesting because although oRGs were initially described as neurogenic progenitors (Fietz *et al*., 2010; Hansen *et al*., 2010), recent work *in vivo* has pointed to their possible contribution to astrocyte and oligodendrocyte production in the human and non-human primate brain (Rash *et al*., 2019; Yue Huang *et al*., 2020). To further explore the extent to which organoids can model the oRG-to-oligodendrocyte lineage, we searched for a recently-described pre-oligodendrocyte precursor cell (pre-OPC) present in human endogenous cortex at late gestational stages and characterized by the co-expression of oRG (*HES1*, *HOPX*, *MOXD1*) and early oligodendrocyte lineage markers (*EGFR*, *PDGFRA*, *OLIG1*, *OLIG2*). At 5 and 6 months *in vitro*, but not at earlier stages, a population expressing these markers was indeed present in organoids (Figure S24B-C). These data support initial findings in the field that human oRG contribute to glial lineages in addition to excitatory neurons during human corticogenesis. Together, these findings demonstrate the value of organoids to investigate human-specific developmental processes that would be otherwise difficult to mechanistically dissect *in vivo*.

To further investigate how progenitors’ transcriptional profile changes as they develop, we examined differential gene expression with each cell type over time (see Methods). Overall, we found that progenitors acquire an expression profile reflective of the progeny they produce (Figure S24D). This was also reflected by gene ontology terms whose expression is up-regulated through organoid development in specific progenitor types, such as “modulation of chemical synaptic transmission” and “gliogenesis” in oRG, and “gliogenesis” in aRGs and IP (Figure S24E). The scATAC-seq data showed a similar pattern, such as the increased accessibility of *APOE* and *PDGFRA* in oRG at 6 months (Figure S24F).

To define the molecular sequence involved in the specification of each cell lineage, we applied the trajectory-inference algorithm URD to generate a branched trajectory tree based on the transcriptional similarity of pseudotime-ordered cells (Figure 6B and S25A-C) (Farrell *et al*., 2018). We defined the root as aRG present at the earliest stage profiled, 23 days *in vitro*, and the tips as the terminal cell types of a given lineage present on the organoids across the stages examined. Because organoids were grown with a protocol to pattern dorsal telencephalon, we excluded cell types known to originate in other regions of the developing neural tube. The order established in the tree reflects the cells’ stage and differentiation status, with basal progenitors following aRG, and then diverging into neuronal (CFuPN, CPN) and glial lineages (astroglia, oligodendrocyte precursors).

We used the tree to map dynamic expression changes over development (Figure 6C and S25D). The early portion of the neuronal trajectories showed expression of cell cycle related and migration genes (*GADD45G*, *SSTR2*, respectively), and upregulation of pan-neuronal genes (*NEUROD2*, *NEUROD6*), in agreement with the molecular trajectories described for cortical neurons of the mouse cortex (Di Bella *et al*., 2021). Later cell-type specific programs involved known lineage-specific genes such as *SOX5*, *BCL11B,* and *FEZF2* for CFuPN, and *CUX2*, *PLXNA4,* and *SATB2* for CPN (Figure 6C, Figure S25C, Table S15).

We then applied this tree to identify lineage-specific genes associated with human cortical neuron subtypes. To this end, we compared the molecular cascades emerging from the human cortical organoid tree with the molecular cascades associated with mouse projection neurons (Di Bella *et al*., 2021). This analysis revealed novel lineage-specific genes in the human organoid tree that showed a different expression pattern in the mouse tree (Figure 6E). For instance, *SORCS1* is associated with the CFuPN lineage in the human organoid tree, while it is expressed across multiple cortical projection neurons in the mouse. Similarly, *PIK3R1* is associated with the CPN lineage in humans, but more broadly expressed in mouse. Additionally, we observed genes associated with human projection neuron lineages which are entirely absent in the mouse developing cortex, such as *HS6ST3* for CPN. Finally, this analysis revealed *FABP7* as a gene associated with the CPN lineage in humans, while in rodents it is exclusively associated with progenitors and glial cells. To confirm the involvement of these genes in the associated lineages, we verified their cell-type-specific expression patterns in the human endogenous fetal datasets used in this study (Figure S25E), and in two additional datasets of the mouse developing cortex (Yuzwa *et al*., 2017; Telley *et al*., 2019).

There is limited knowledge of how human cellular diversification in the cortex is achieved, and the molecular program associated with human lineage bifurcation decisions remain undefined. In order to address these gaps in knowledge, we applied the URD tree to identify genes associated with lineage bifurcations in organoids. We defined differential gene expression among the parent and daughter branches at each branch point, trained a gradient boosting decision tree to assign an importance score to each gene, and selected the 20 highest-scoring genes for each branch (Figure S25F), as well as the top 20 TFs (Figure 6D). The top TFs for each daughter branch included genes with known functions in neurogenesis, gliogenesis, and/or cell identity acquisition (e.g., Neuronal daughter branch: *NEUROD6*, *NEUROD4*, *NEUROG2*; Glial daughter branch: *ZBTB20*, *ASCL1*; CFuPN daughter branch: *BCL11B*, *FEZF2*; CPN daughter branch: *POU3F2*, *MEF2C*).

To identify human genetic programs governing these bifurcations, we compared the most important TFs in the organoid tree branch points with TFs and genes in a recently-published mouse developing cortex URD tree (Figure S25G, (Di Bella *et al*., 2021)). This resulted in the discovery of new TFs, expressed both in the human and mouse developing cortex, potentially governing neurogenesis and/or gliogenesis in the mammalian cortex (*DACH1*, *TSHZ1*, with higher importance score in the neuronal branch; *CXXC5*, *ZFHX4* with higher importance score in the glial branch, Figure S25H), as well as new candidate regulators of identity acquisition for specific cortical projections neurons (*ZNF704* for CPN, Figure S25H). Although all of these genes were expressed in the mouse developing cortex, some of them showed a different expression pattern across the mouse pseudotime tree (*ZNF704*/*Zfp704*, Figure S25H) suggesting divergent species-specific functions of certain TFs. In the human organoid tree, we also identified TFs putatively mediating astrocyte vs oligodendrocyte lineage commitment (e.g., *CEBPD*, *MESP1* with high importance scores in the oligodendrocyte lineage daughter branch, Figure S25I), a molecular process poorly explored in human corticogenesis. Finally, by comparing with the mouse tree, we discovered novel candidate regulators with no 1:1 mouse ortholog (Ruan *et al*., 2008) that may have a role in neuronal commitment in human corticogenesis: e.g., ZNF26 and ZNF37A (Figure 6F).

In order to have a complete picture of the molecular logic underlying human cortical projection neuron specification, we next built a new branched trajectory tree using the SHARE-seq map, containing the transcriptome and epigenome from the same individual cells (Figure S25J-L and Supplementary Text). The URD algorithm was applied to the RNA-seq data to generate a tree, and then the corresponding epigenetic data was projected onto the tree to identify TFs predicted to act in each of the projection neuron lineages represented in this tree, i.e., CFuPN and CPN (Methods and Supplementary Text). This analysis identified known and novel TFs that may act at different stages of neuronal development (Figure S25M). Of interest, the motif for *ZNF264*, which has no mouse ortholog (Ruan *et al*., 2008), was enriched in the early steps of the CFuPN lineage. We confirmed the expression of this gene in projection neurons from three human fetal datasets (the novel fetal dataset as well as (Polioudakis *et al*., 2019; Trevino *et al*., 2021)). This data suggests a species-specific role for *ZNF264* in human projection neuron development and more broadly highlight newly-emerging human-specific regulatory processes of neurodevelopment in the neocortex.

### Human callosal projection neuron diversity emerges during embryonic development

Recent work has described a higher gene-expression heterogeneity in the CPN population in the adult human cortex when compared to the mouse, suggesting that the expansion of this neuronal population in humans has been accompanied by increased diversification (Berg *et al*., 2021). However, how and when these different CPN types arise remains unknown. We examined the terminal neuronal cell types of our tree to investigate if the gene-expression CPN heterogeneity described in adulthood (Hodge *et al*., 2019; Berg *et al*., 2021) was already detectable during early stages of development. We first examined the expression of marker genes of the five different CPN types (Hodge *et al*., 2019) in fetal and organoid CPN (Figure 7A). We observed that some of these markers were expressed from early stages of development, and that fetal and organoid CPN showed a similar pattern of expression, i.e., those markers with a high expression in fetal CPN also showed a high expression in organoid cells, and vice versa. To assess if the molecular diversity found in adulthood could also be observed in early CPN, we applied consensus Non-negative Matrix Factorization (cNMF, (Kotliar *et al*., 2019)) to identify joint activity of gene programs (gene modules) in adult CPN (Table S16). We found five gene modules broadly representing each of the adult CPN types (Figure 7B-C, Figure S26A-B). Then, we assessed the expression of these gene modules both in fetal and organoid CPN. We observed that the modules were differentially expressed across cell populations both *in vivo* and *in vitro*, although expression was not completely discrete, with module 5 showing the strongest divergence from the others (Figure 7D-E and S26C-I). Therefore, although the full gene-expression diversity observed in human adult CPN is not yet present, our data indicate that CPN divergence begins at early stages of development, providing a model to study this process.

**Figure 7.**
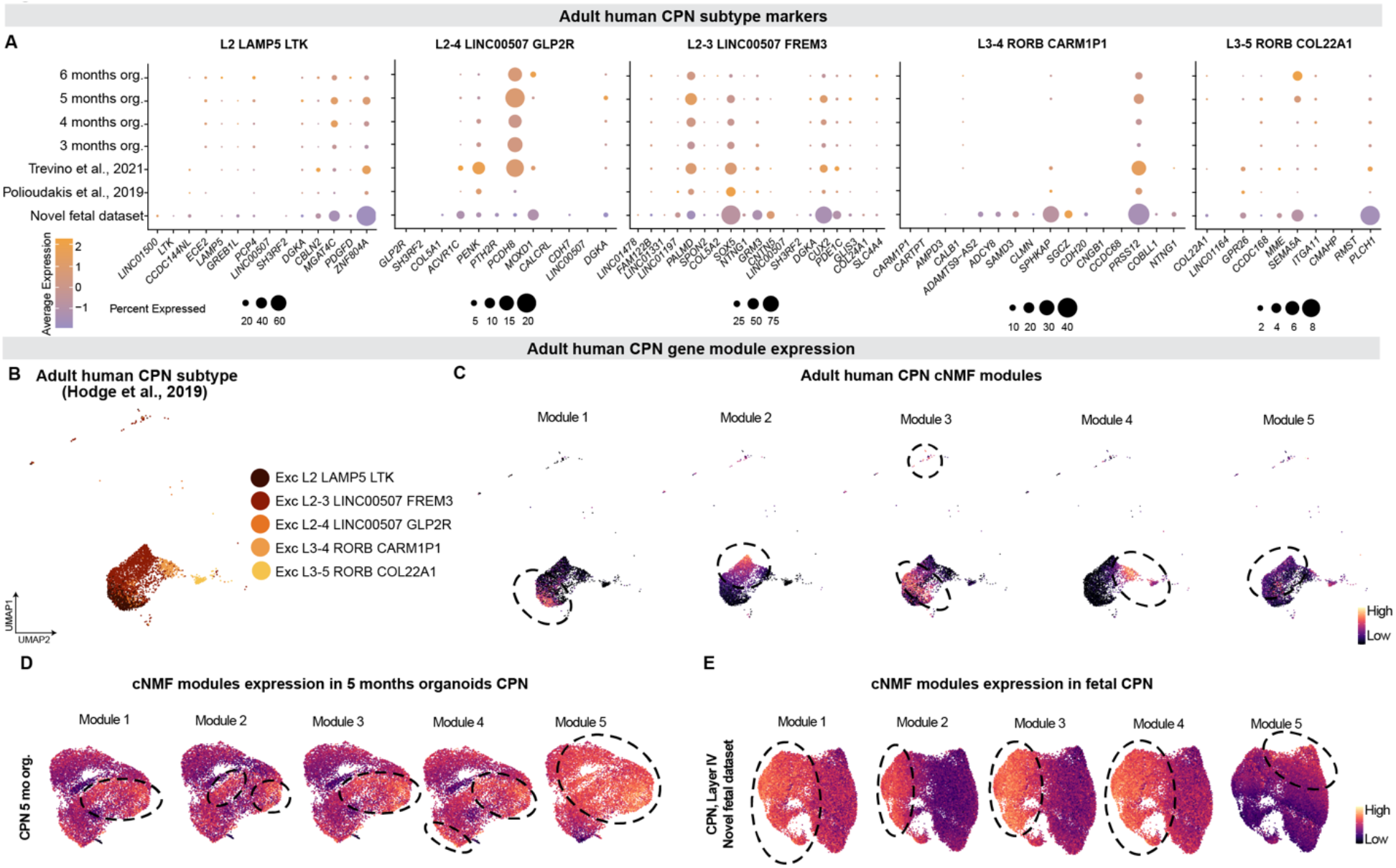
Callosal projection neuron transcriptional diversity emerges at early stages of cortical development. (A) Dotplot showing the percentage of cells (dot size) and average expression (color) of adult CPN subtypes markers (Hodge *et al*., 2019) in human organoid and fetal CPN. (B) UMAP displaying the five transcriptional subtypes of adult CPN described in (Hodge *et al*., 2019; Berg *et al*., 2021). (C) UMAP feature plot of adult CPN showing the expression of cNMF (consensus Non-negative Matrix Factorization) gene modules. Each cNMF module corresponds to a CPN transcriptional subtype. Dotted circles show areas of high gene module expression. (D-E) UMAP of 5 month organoid CPN (D) and human fetal CPN (novel fetal dataset, E) generated using as variable features the top 100 genes used in each of the cNMF modules. The UMAPs are colored by the module scores of genes in each cNMF module. Dotted circles show areas of high gene module expression.

## Discussion

Understanding the mechanisms that govern the generation of cell diversity and overall development of the human brain are major unmet goals, largely impeded by the ethical and practical limitations associated with profiling the fetal human brain over extended periods of *in utero* development. Brain organoids offer the opportunity to experimentally investigate aspects of human brain development that would not otherwise be amenable to research. However, in order to fulfill this promise, a high-resolution understanding of how organoids develop relative to endogenous fetal brain is necessary. Here, we have generated a comprehensive, single-cell resolution transcriptional and epigenetic developmental map of a human organoid model of the cerebral cortex (Velasco *et al*., 2019), across an extended timeline of *in vitro* development. We find that this model displays high levels of developmental fidelity and reproducibility across longitudinal processes of cellular diversification, displaying consistent acquisition of cell identity in each organoid, implemented through reliable molecular programs.

Previous work has indicated that the conditions under which organoids are cultured can affect their metabolic state (Pollen *et al*., 2019; Bhaduri *et al*., 2020; Tanaka *et al*., 2020), and that a change in growth environment, such as transplantation into endogenous tissue, can improve culture-related metabolic abnormalities (Bhaduri *et al*., 2020). Extending these findings, our work here shows that it is indeed possible to consistently generate accurate cell identities independent of metabolic state in organoids, under specific culture conditions.

It is interesting that while the molecular identities of the vast majority of cortical cell types in organoids compared to their fetal counterparts was not affected by metabolic gene sets, the identities of two cell types (aRG and ‘unspecified PN’) were affected by glucose metabolism and hypoxia. This may be related to the localization of these cell types disproportionately towards the more hypoxic center of the organoids, supported by our finding that these metabolic signatures were more prominent at advanced stages of organoid maturation, when organoid size is larger. Procedures to increase organoid oxygenation and nutrient diffusion, for example by vascularization and perfusion (Del Dosso *et al*., 2020) may in the future improve these cell states. However, the involvement of glucose metabolism is notable, as several prior studies have reported that neuronal progenitors have a predominantly glycolytic metabolism during normal brain development (Llorens-Bobadilla *et al*., 2015; Khacho *et al*., 2016; Zheng *et al*., 2016; Beckervordersandforth *et al*., 2017; Knobloch and Jessberger, 2017; Namba *et al*., 2020), and that the transition to a more differentiated state (i.e., from neural progenitor to neurons) is indeed accompanied *in vivo* by a metabolic switch away from glycolysis and towards mitochondrial oxidative phosphorylation (Knobloch and Jessberger, 2017; Khacho, Harris and Slack, 2019). In agreement, we find that fetal progenitors also show an enrichment of glycolytic genes relative to other endogenous fetal cell types. It is therefore possible that progenitors in organoids may exhibit an exacerbated alteration of metabolic pathways that are normally more mildly enriched in these same cells during endogenous developmental process in the embryo.

Our comparison of reconstructed developmental trajectories of individual cell lineages in human organoids versus the corresponding lineage trees in developing mouse neocortex (Di Bella *et al*., 2021) predicted novel regulators of cell identity acquisition in humans. This analysis identified two genes of the ZNF family with no mouse ortholog that are associated with the neurogenic vs. gliogenic switch (ZNF26 and ZNF37A), as well as TFs associated with projection neuron class diversification in humans (ZNF704). This finding is consistent with prior reports that the developing human cortex is enriched for genes of recent evolutionary origin (Florio *et al*., 2018; Suzuki *et al*., 2018) and points at the value of organoids to identify human-specific mechanisms of development for further functional investigation.

During evolution, callosal projection neurons of the cortical upper layers have undergone a remarkable expansion and diversification in the human cortex relative to other species (Sousa *et al*., 2017; Hodge *et al*., 2019; Miller *et al*., 2019; Berg *et al*., 2021). Here, we discover that the transcriptional diversity observed for this neuronal population in the adult human cortex (Hodge *et al*., 2019; Berg *et al*., 2021) begins to emerge during early stages of cortical development in both organoids and fetal tissue, suggesting that this diversity reflects specific developmental events rather than depending on later factors such as neuronal activity. This finding points at developing organoids as systems to explore the mechanistic underpinning of this critical, human-specific process, and further reinforces the value of organoids as experimental models to investigate otherwise-inaccessible mechanisms of human cortical development, evolution, and disease.

## Author contributions

P.A., J.Z.L., A.J.K., and A.U. conceived the experiments. A.U., S.V., B.P., M.P., and A.T. generated, cultured, and characterized all the organoids in this study and P.A. supervised their work and contributed to data interpretation. X.A. performed scRNA-seq and scATAC-seq experiments, with help from A.U., M.P., B.P., and S.V.; A.J.K., K.K., T.F., A.U., and J.Z.L. performed scRNA-seq and scATAC-seq analysis and J.Z.L and A.R. supervised the computational work, with input from A.R.; S.N. performed and analyzed SHARE-Seq experiments, supervised by J.B. N.A.B. and C.G. generated the human fetal cortex snRNAseq data. A.U., S.V., B.P., M.P., and A.J.K. worked on cell type assignments and data analysis and P.A. supervised the work. X.J., E.M., and F.C. performed and analyzed Slide-seqV2 experiments. P.A., A.U., and A.J.K. wrote the manuscript with contributions from all authors. All authors read and approved the final manuscript.

## Acknowledgments

We thank J. R. Brown and S. Simmons for editing the manuscript, and the entire Arlotta Lab for support and insightful discussions; A. Shetty and D. Di Bella for help with scRNA-seq cell type classification; V. Vuong, C. Abbate, R. Sartore for technical support in organoid culture and characterization; the Broad Genomics Platform for sequencing; the Talkowski laboratory for the GM08330 line; the Cohen laboratory for the Mito210 line. This work was supported by grants from the Stanley Center for Psychiatric Research, the Broad Institute of MIT and Harvard, the National Institutes of Health (R01MH112940 to P.A. and J.Z.L., and P50MH094271, U01MH115727, and RF1MH123977 to P.A.), the Klarman Cell Observatory to J.Z.L. and A.R., and the Howard Hughes Medical Institute to A.R. AR was a Howard Hughes Medical Institute and a Koch Institute extramural member while conducting this study.

## Supplemental Figures

**Figure S1, related to Figure 1:**
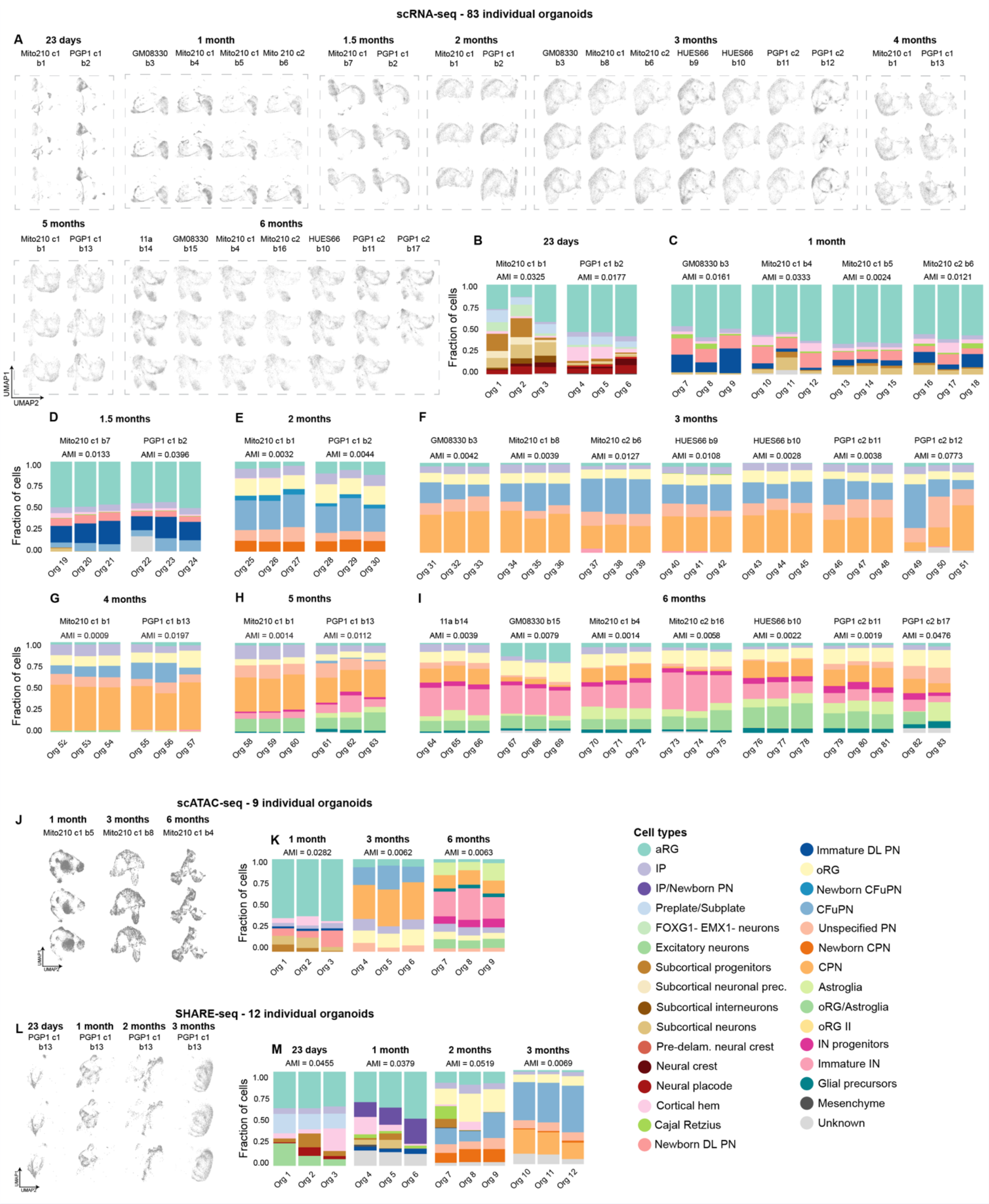
Cell-type proportions across individual organoids. (A) Contributions to UMAPs from individual organoids from scRNA-seq datasets across all timepoints and differentiation batches sequenced, plotted on the same UMAPs from Figure 1A. (B-I) Proportion of cells belonging to each cell type from each individual organoid from scRNA-seq datasets. Adjusted mutual information (AMI) between cell type assignments and organoid identity is given per batch, where lower AMI indicates lower organoid-to-organoid variability. AMI measures the dependence between two variables (cell type and organoid of origin), adjusted for agreement due to chance; low scores indicate high independence between cell type and the identity of the specific organoid a cell came from. (J) Contributions to UMAPs from individual organoids from scATAC-seq datasets across all timepoints and differentiation batches sequenced, plotted on the same UMAPs from Figure 1C. (K) Proportion of cells belonging to each cell type from each individual organoid from scATAC-seq datasets. AMI as in B. (L) Contributions to UMAPs from individual organoids from SHARE-seq datasets across all timepoints and differentiation batches sequenced, plotted on the same UMAPs from Figure 1D, in which the cells for a given timepoint were subsetted. (M) Proportion of cells belonging to each cell type from each individual organoid from SHARE-seq datasets. AMI as in B.

**Figure S2, related to Figure 1:**
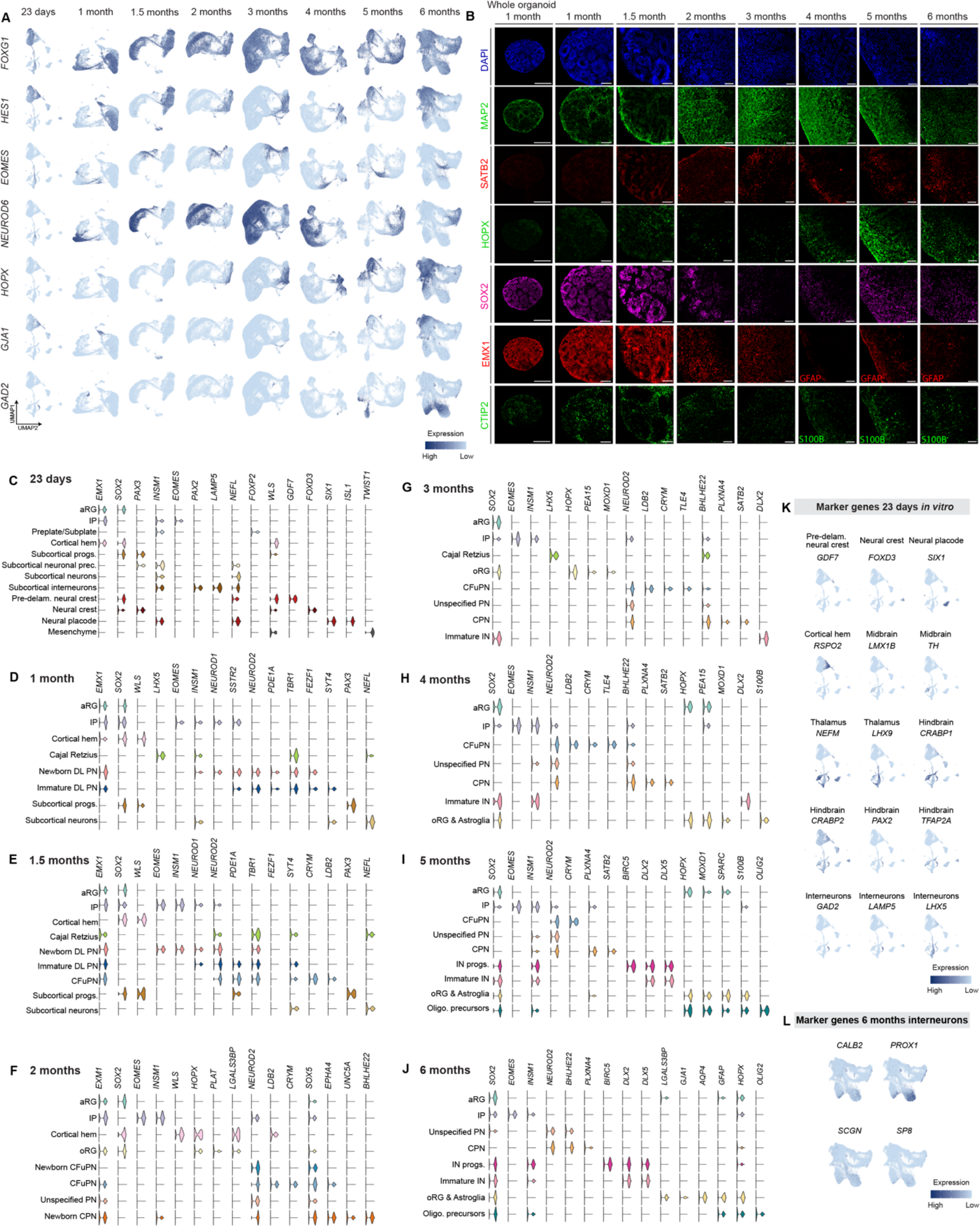
Marker gene expression for major cell types. (A) Feature plots showing normalized expression of marker genes of cortical cell types across organoids after 23 days to 6 months in culture, plotted on the same UMAPs from Figure 1A: telencephalon (*FOXG1*), aRG and oRG (*HES1*), IP (*EOMES*), dorsal excitatory neurons (*NEUROD6*), oRG *(HOPX*), astroglia (*GJA1*), interneurons (*GAD2*). (B) Immunohistochemistry for neuronal (MAP2), dorsal forebrain progenitor (EMX1, SOX2), CFuPN (CTIP2), CPN (SATB2), oRG (HOPX) and astroglia (GFAP, S100B) cell types, across organoids after 1-6 months in culture. Scale bar 100 μm, except first column (whole organoid), 300 μm. 1-, 2-, 3-, 4-, 5-, and 6-month organoids correspond to Mito210 c1 line b1. 1.5-month organoid corresponds to Mito210 c1 b7. 6-month organoid corresponds to Mito210 c1 b4. (C-J) Violin plots showing marker gene expression across cell types in organoids from 23 days to 6 months in culture. For this plot, oRG and astroglia cells were merged into “oRG & Astroglia” at 4-6 months. (K) Feature plots showing normalized expression of marker genes for non-cortical cell types in organoids after 23 days in culture. The cell type or brain region for which each gene is a marker is indicated. (L) Feature plots showing normalized expression of marker genes for late-stage immature interneurons in 6 month organoids harmonized UMAP.

**Figure S3, related to Figure 1:**
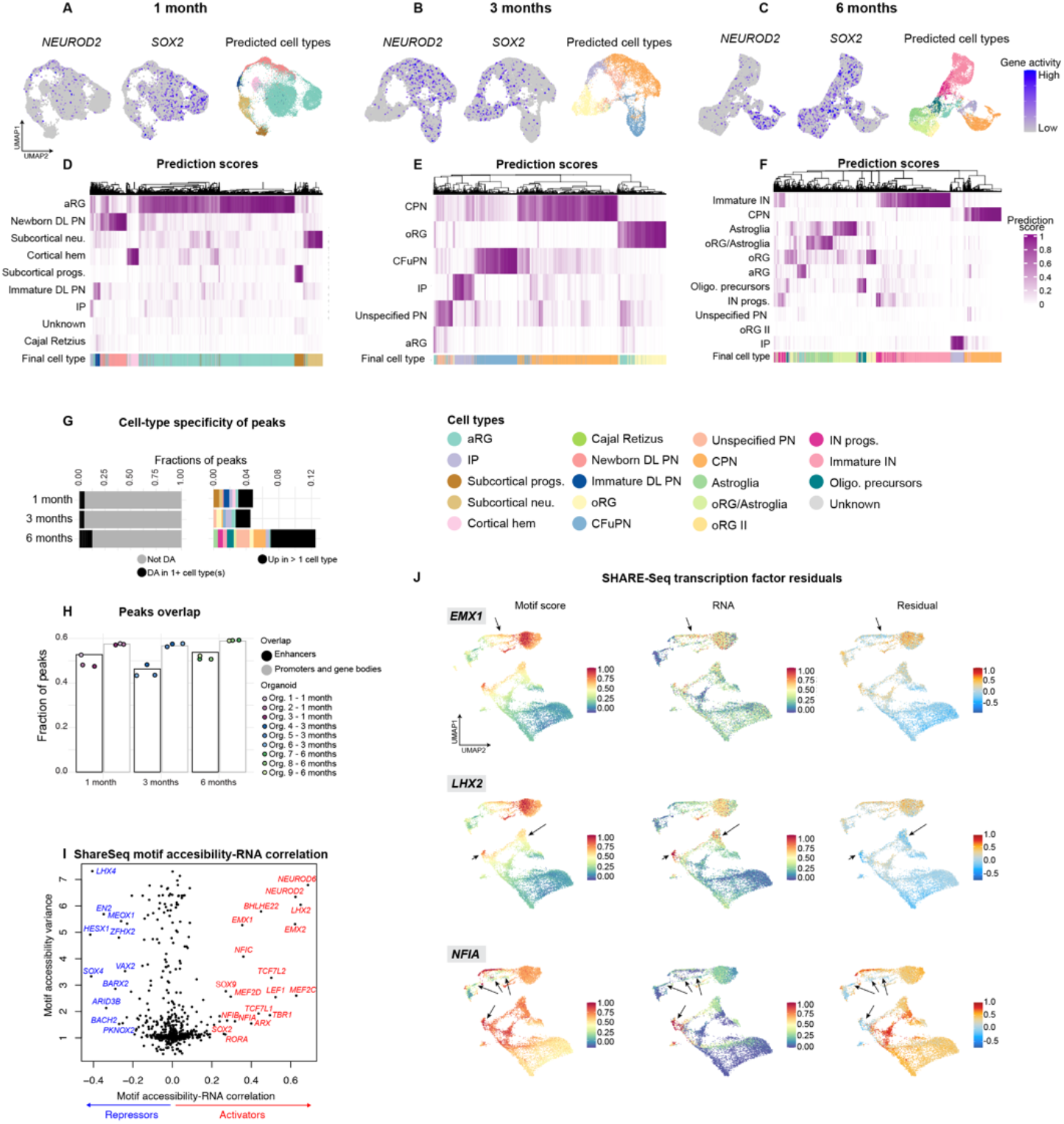
Cell types and peak characteristics from epigenetic data. (A-C) Feature plots showing the gene activity (calculated from reads in the promoter and gene bodies) of neuronal marker gene *NEUROD2* and progenitor marker gene *SOX2* in scATAC-seq data of 1-, 3-, and 6-month organoids, as well as cell types predicted after integration with scRNA-seq data from matching timepoints. (D-F) Heatmaps showing the prediction scores for each cell corresponding to each scRNA-seq label. (G) Fraction of total peaks called in each scATAC-seq dataset that were called as differentially accessible (DA) in at least one cell type versus all other cells in that timepoint. (H) Fraction of total peaks called in each scATAC-seq dataset that overlap with enhancers (black outline) and with gene bodies and promoters (gray outline). Bar chart represents the overlap of the peak set after merging three organoids from each timepoint (these peak sets were used in downstream analyses), and individual points represent peaks called from an individual organoid. The proportion of peaks overlapping promoter and gene body regions (57-59% per timepoint) was modestly higher than the fraction overlapping annotated enhancer regions (46-54%), and did not change significantly over time. (I) Transcription factor motif (TF) accessibility related to the gene expression of the cognate TF, from SHARE-seq data in day 23 through 3 month organoids. (J) Difference (residuals) between chromatin accessibility and gene expression for the transcription factors *EMX1*, *LHX2*, and within the SHARE-seq data. Arrows point to regions where there is early RNA expression without high motif accessibility.

**Figure S4, related to Figure 2:**
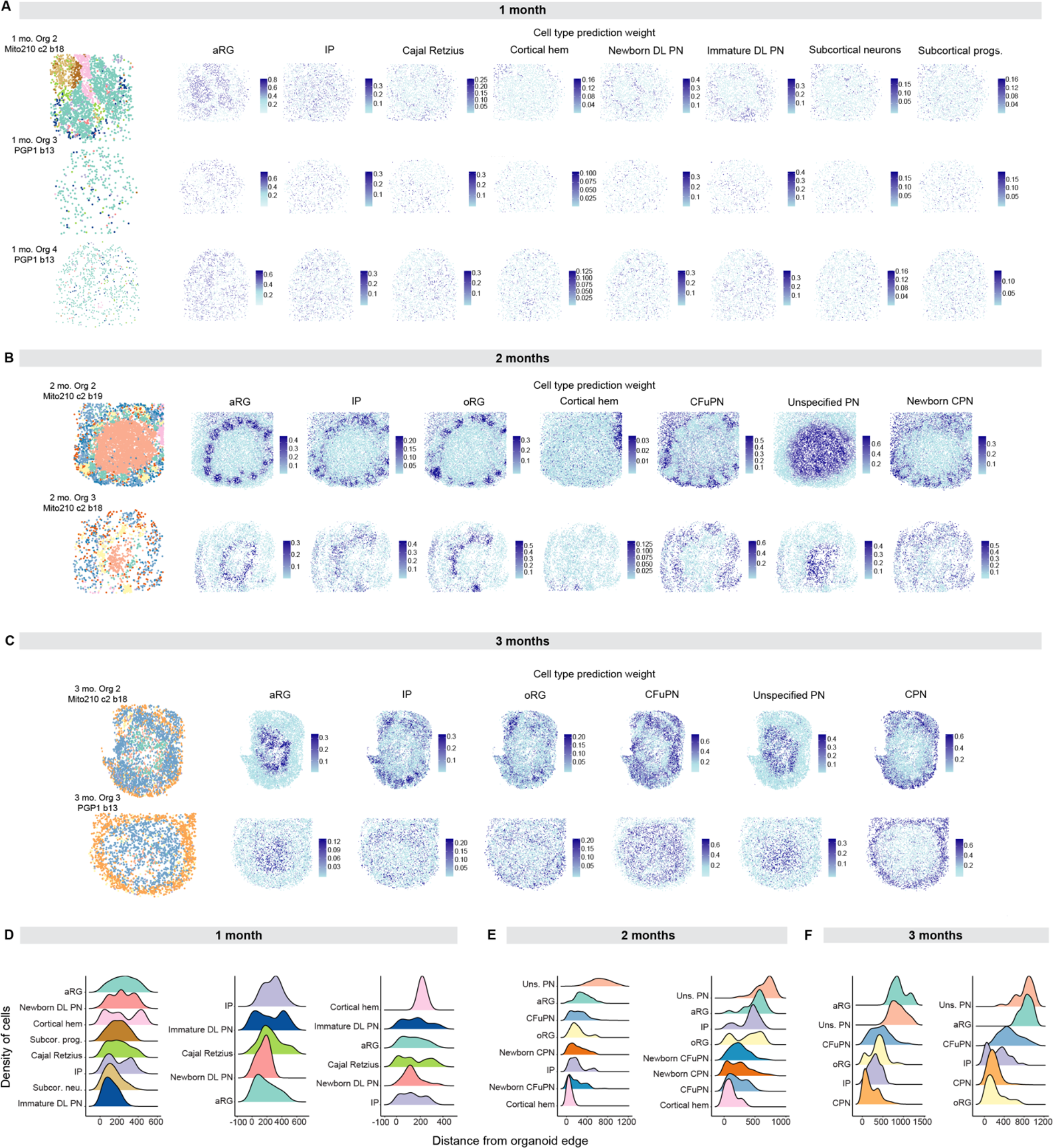
Spatial transcriptomic landscape of additional organoids. (A-C) Left, spatial plot of Slide-seqV2 data from 1-month Organoids 2-4 (A), 2-month Organoids 2-3 (B), and 3-month Organoids 2-3 (C), colored by RCTD-assigned cell type (Colors as in D). Right, spatial plots showing RCTD prediction weights. (D-F) The distribution of each annotated cell type over the beads’ calculated distance from the edge of the organoid, ordered from top to bottom by median distance.

**Figure S5, related to Figure 3:**
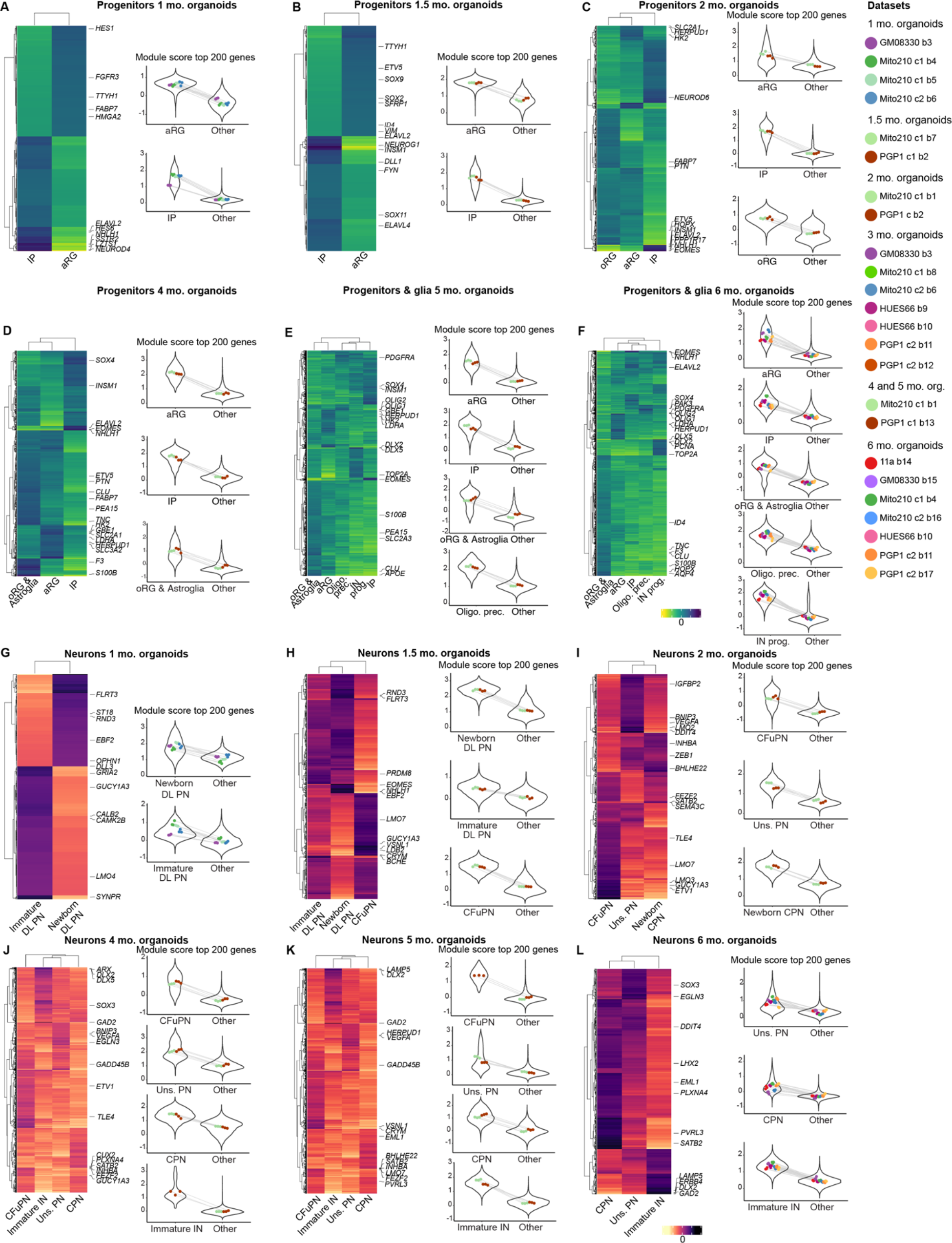
Gene signatures in progenitor and neuron populations over time. (A-L) Left, heatmap comparing the upregulated genes in each progenitor or neuronal population at 1 month (A and G, Table S4), 1.5 months (B and H, Table S5), 2 months (C and I, Table S2), 4 months (D and J, Table S6), 5 months (E and K, Table S7), and 6 months (F and L, Table S8), showing genes with adjusted p value < 0.0015 and log2 fold change > 1.5. At 4, 5 and 6 months, oRG, oRG/astroglia, and astroglia were merged in one oRG & Astroglia cell category. At 2 months, newborn CFuPN and CFuPN were merged in one CFuPN cell category. Right, gene set expression of the top 200 genes in the molecular signatures of each cell type. Violin plots show the module score distribution across single cells, points show the mean module score for each individual organoid, with lines connecting points representing cells from the same organoid. The means within each distribution (separately, within the cell type for that signature, left, or the background cells, right, for each plot) showed no significant difference across individual organoids in all cases (one-way ANOVA, p > 0.05).

**Figure S6, related to Figure 3:**
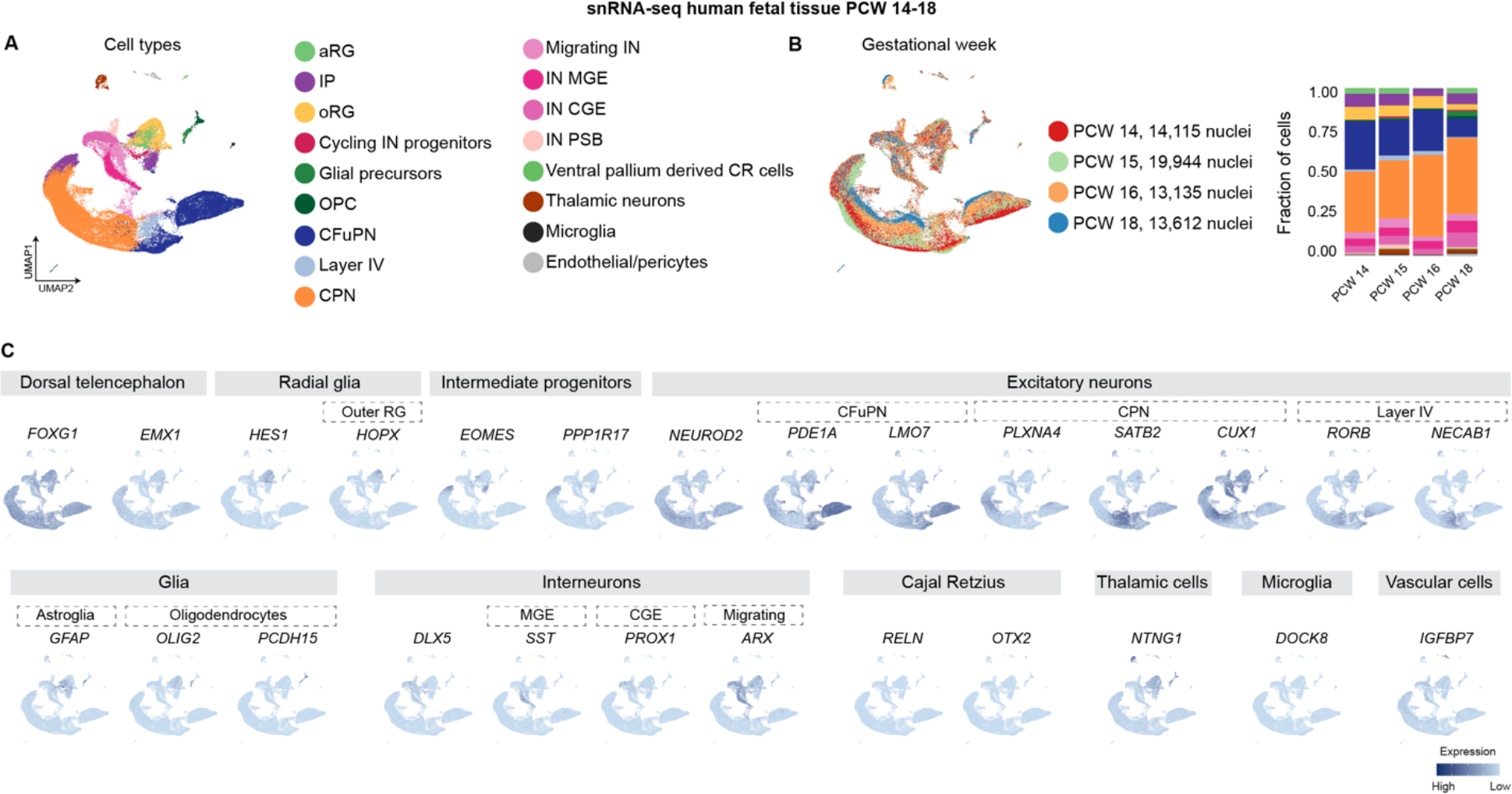
Novel human fetal cortex snRNA-seq dataset spanning PCW 14, 15, 16 and 18. (A) UMAP of snRNA-seq of human fetal cortical tissue from 14, 15, 16 and 18 post conception weeks. Cells are colored by cell type. (B) Left, UMAP with cells colored by PCW. Right, the fraction of cell types per PCW. The number of cells per PCW is indicated. (C) Feature plots showing normalized expression of marker genes of cortical cell types. The cell type or region for which each gene is a marker is indicated above the plots.

**Figure S7, related to Figure 3:**
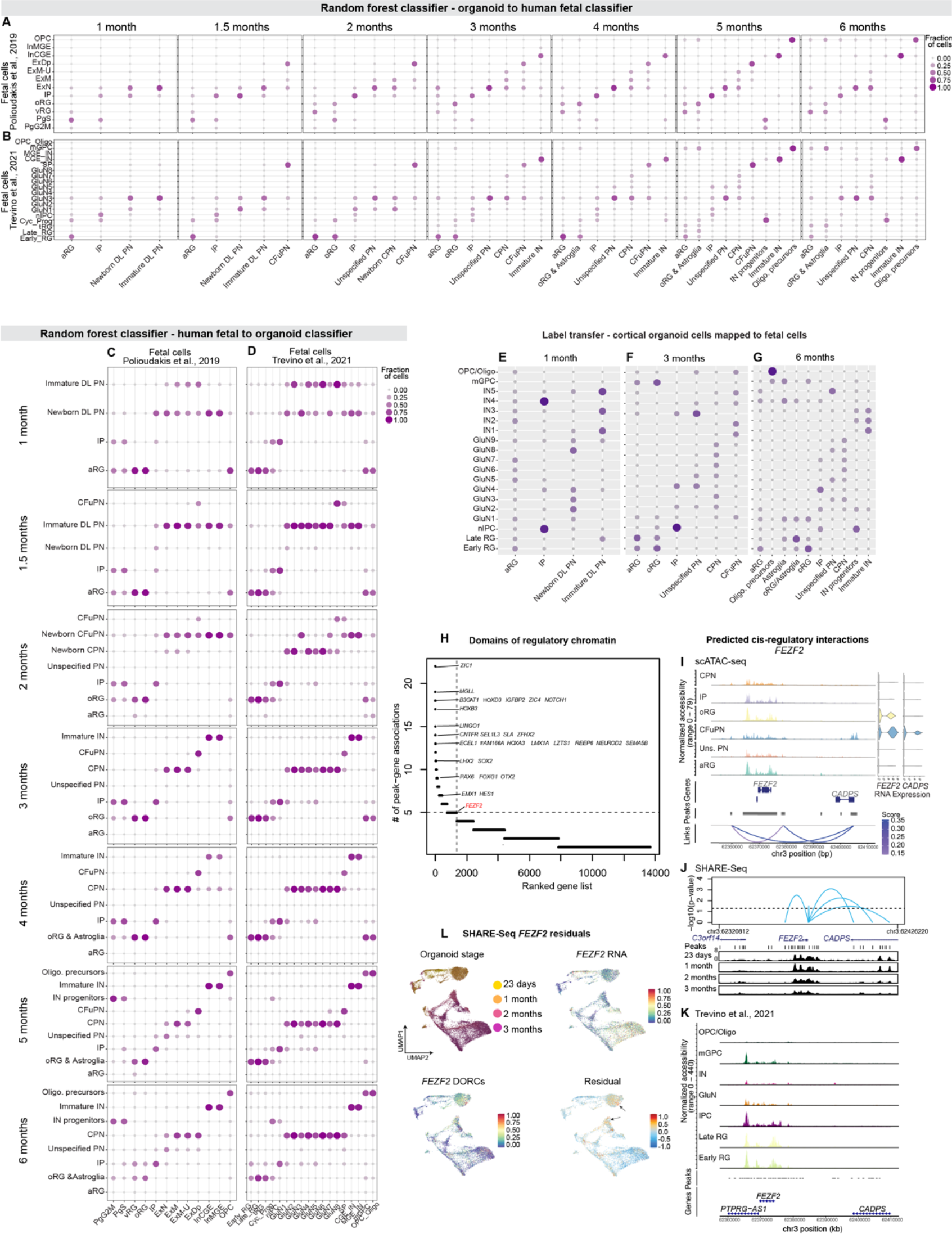
Transcriptional and epigenetic matching of human cortical organoid and fetal cell types. (A-B) Assignments of 1 to 6 months organoid cell types to human cortical fetal cell types (A: (Polioudakis *et al*., 2019), B: (Trevino *et al*., 2021)), as classified by random forest. Points are sized and colored by the fraction of each organoid cell type assigned to the fetal cell types. At 1 and 1.5 months, Cajal Retzius, Cortical hem, and subcortical cells, which are not expected to have corresponding cells in the fetal data, were excluded. (C-D) Assignments of human fetal cell types (C: (Polioudakis *et al*., 2019), D: (Trevino *et al*., 2021)) to 1-6 months organoid cell types, as classified by random forest. Points are sized and colored by the fraction of each human fetal cell type assigned to the organoid cell types. At 1 and 1.5 months, Cajal Retzius, Cortical hem, and subcortical cells, which are not expected to have corresponding cells in the fetal data, were excluded. Classifiers including 3 months organoids cells and (Polioudakis *et al*., 2019) are repeated from Figure 3 for ease of comparison. (E-G) Assignments of 1, 3, and 6 months organoid cell types to human cortical fetal cell types (Trevino *et al*., 2021) using scATAC-seq data, as classified by Seurat’s reference mapping and label transfer. (H) Genes with the greatest number of significantly correlated peaks using SHARE-seq data (p < 0.05, +/- 50 kb from transcription starting site), identifying domains of regulatory chromatin (DORCs). *FEZF2* is highlighted in red. (I) Genome tracks showing scATAC-seq reads from 3-month organoids in the region of *FEZF2*, along with detected peaks (gray bars) and Cicero co-accessibility between peaks (purple curves). Curves with a Cicero score >0.1 are shown. On the right, violin plots showing normalized mRNA expression for *FEZF2* and *CADPS* from scRNA-seq from corresponding Mito210 c1 b6 organoids after 3 months in culture. (J) Genome tracks showing SHARE-seq ATAC reads from 23 days, 1-, 2-, and 3-month organoids in the region of *FEZF2* and *CADPS.* Blue curves indicate peaks significantly correlated with *FEZF2* expression from SHARE-seq RNA data. (K) Genome tracks showing scATAC-seq reads from human fetal tissue (Trevino *et al*., 2021) in the region of *FEZF2* and *CADPS*. (L) Difference (residuals) for *FEZF2* between chromatin accessibility of its corresponding DORCs and gene expression within the SHARE-seq data.

**Figure S8-S14, related to Figure 3:**
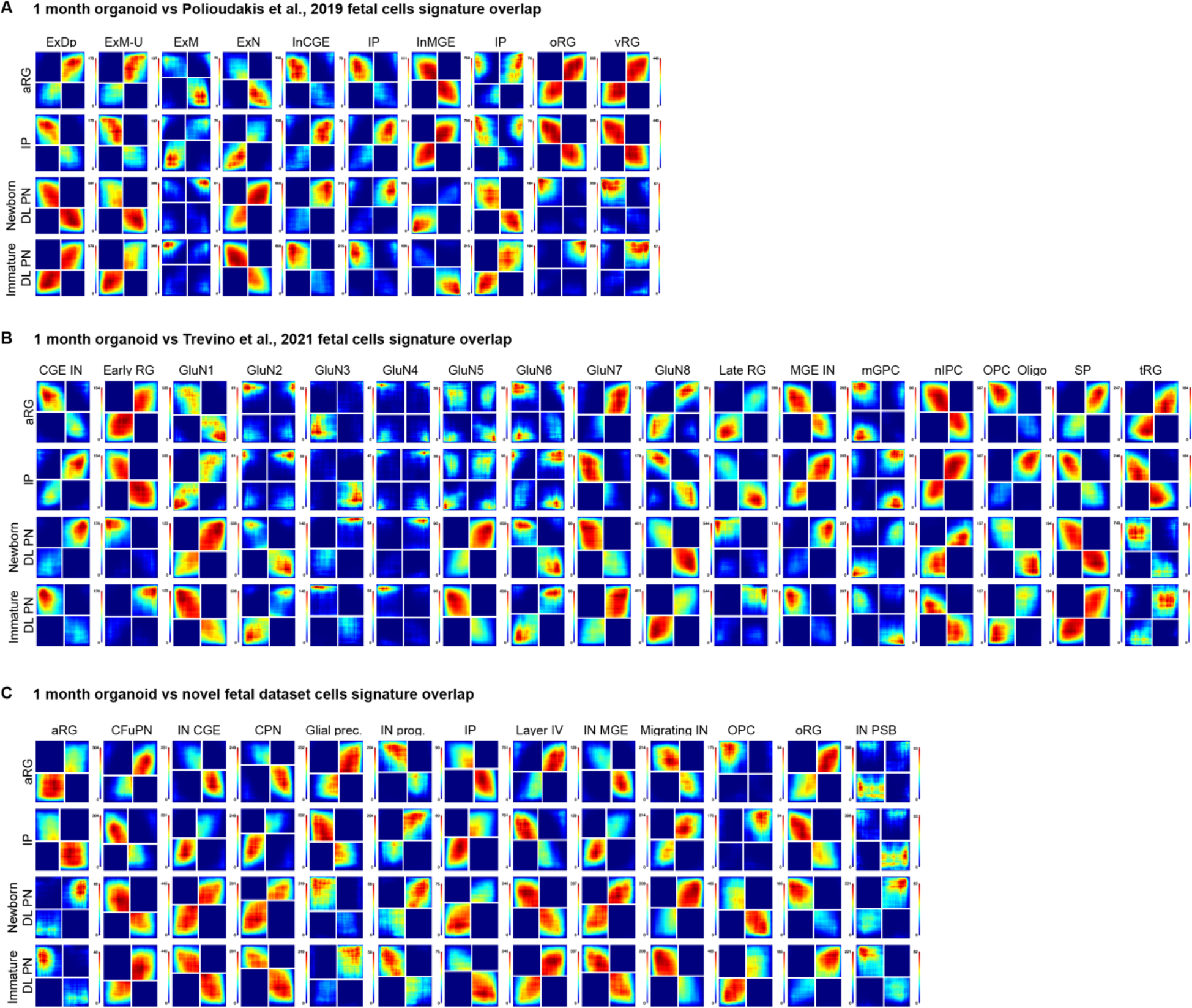
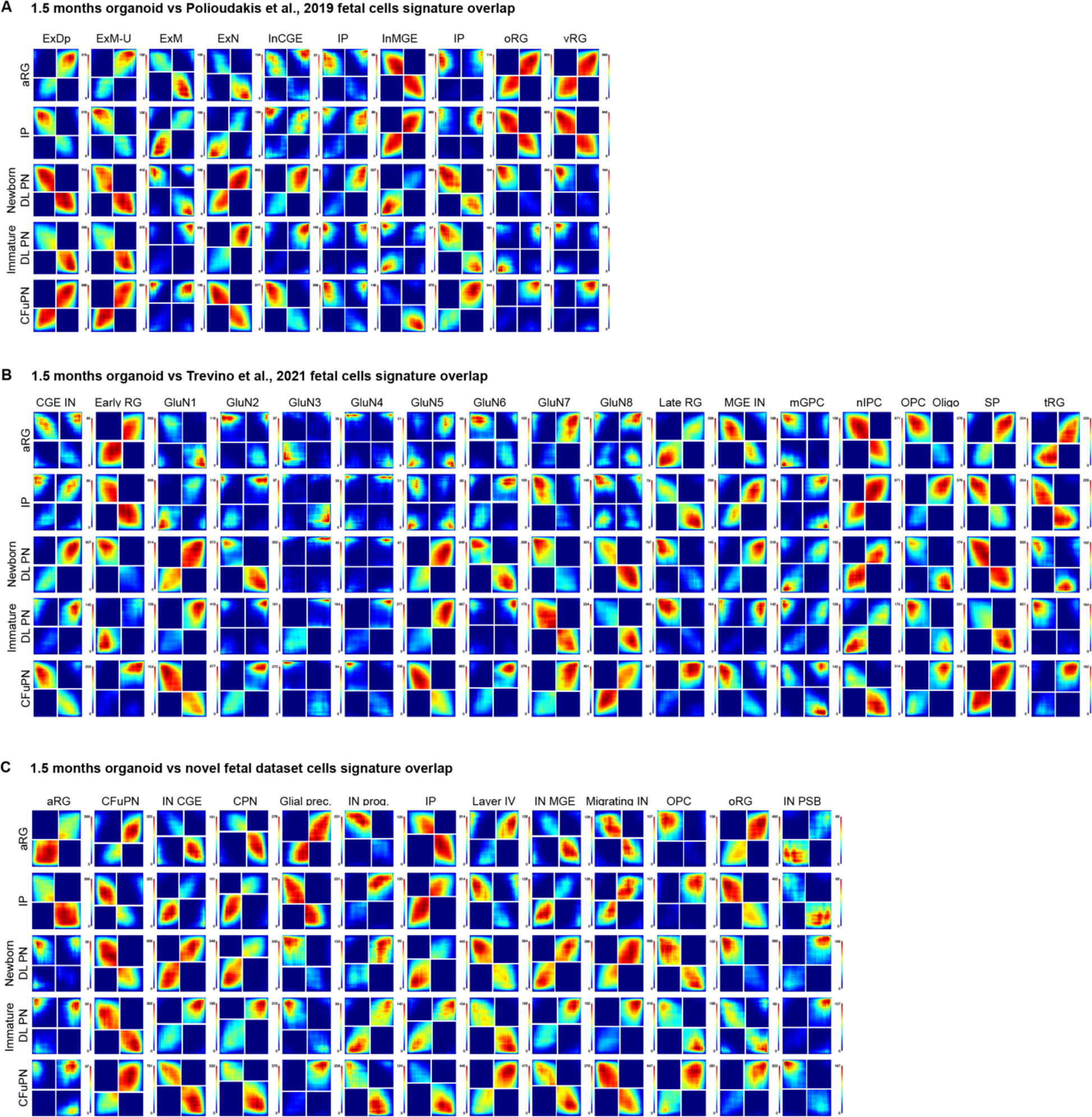
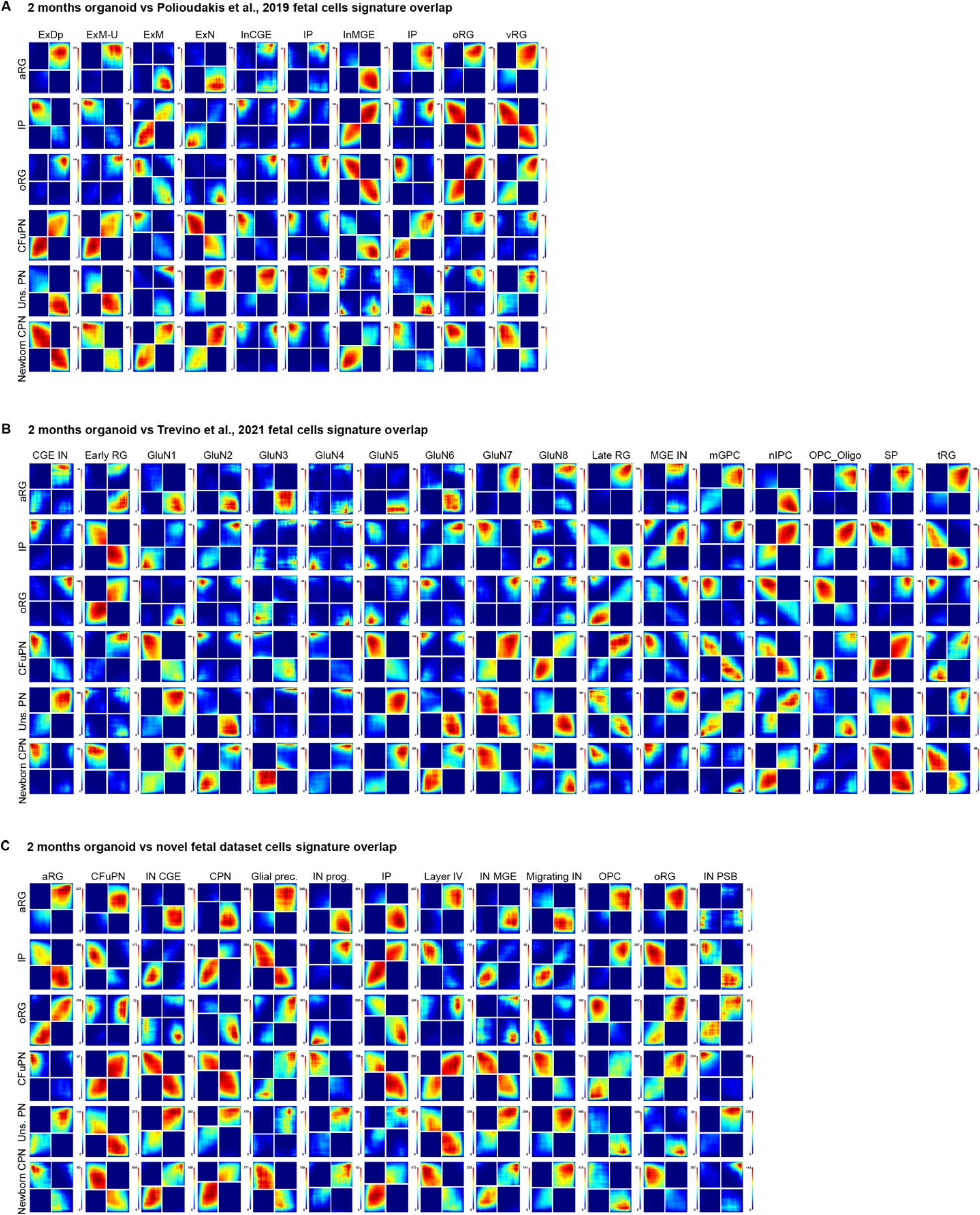
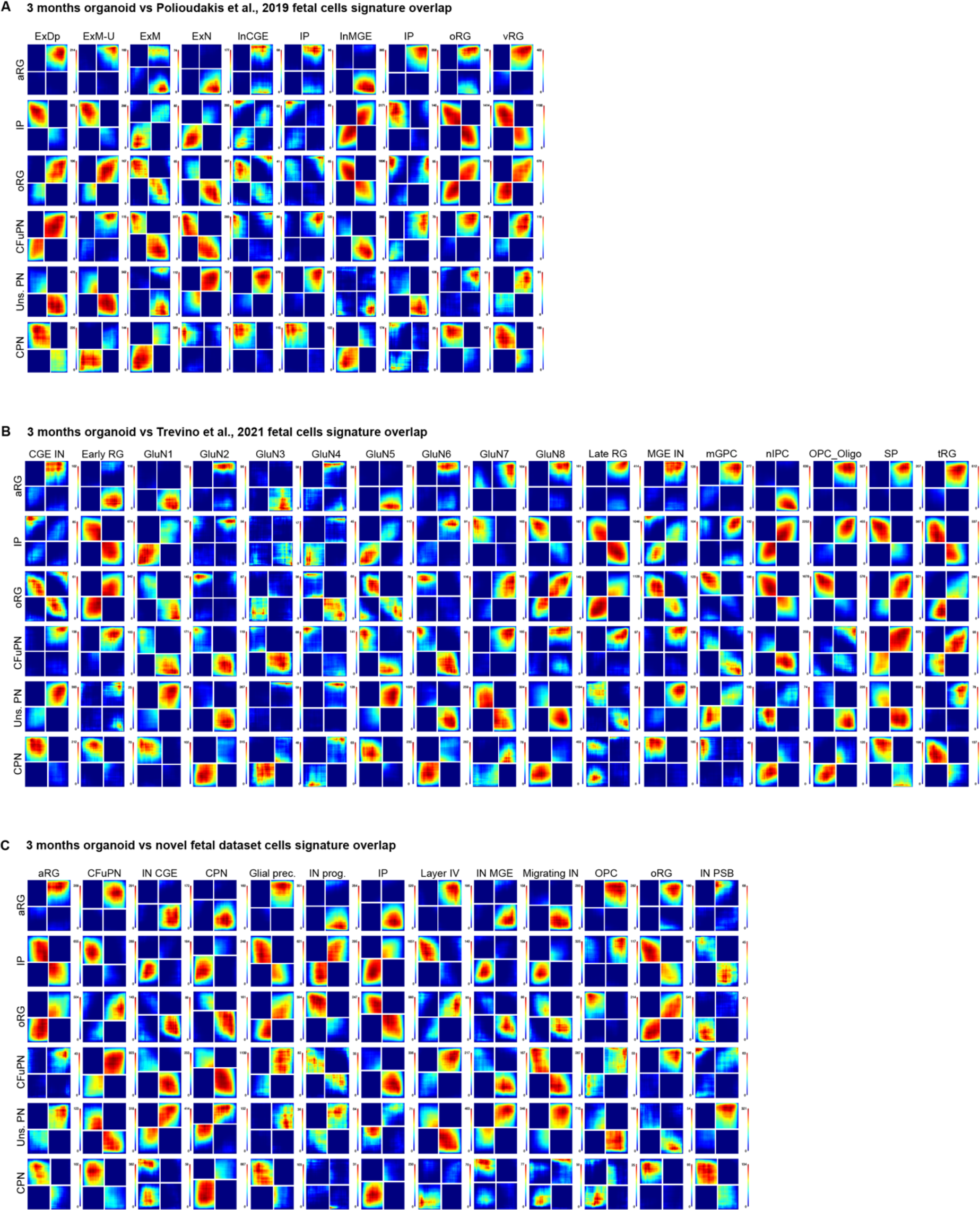
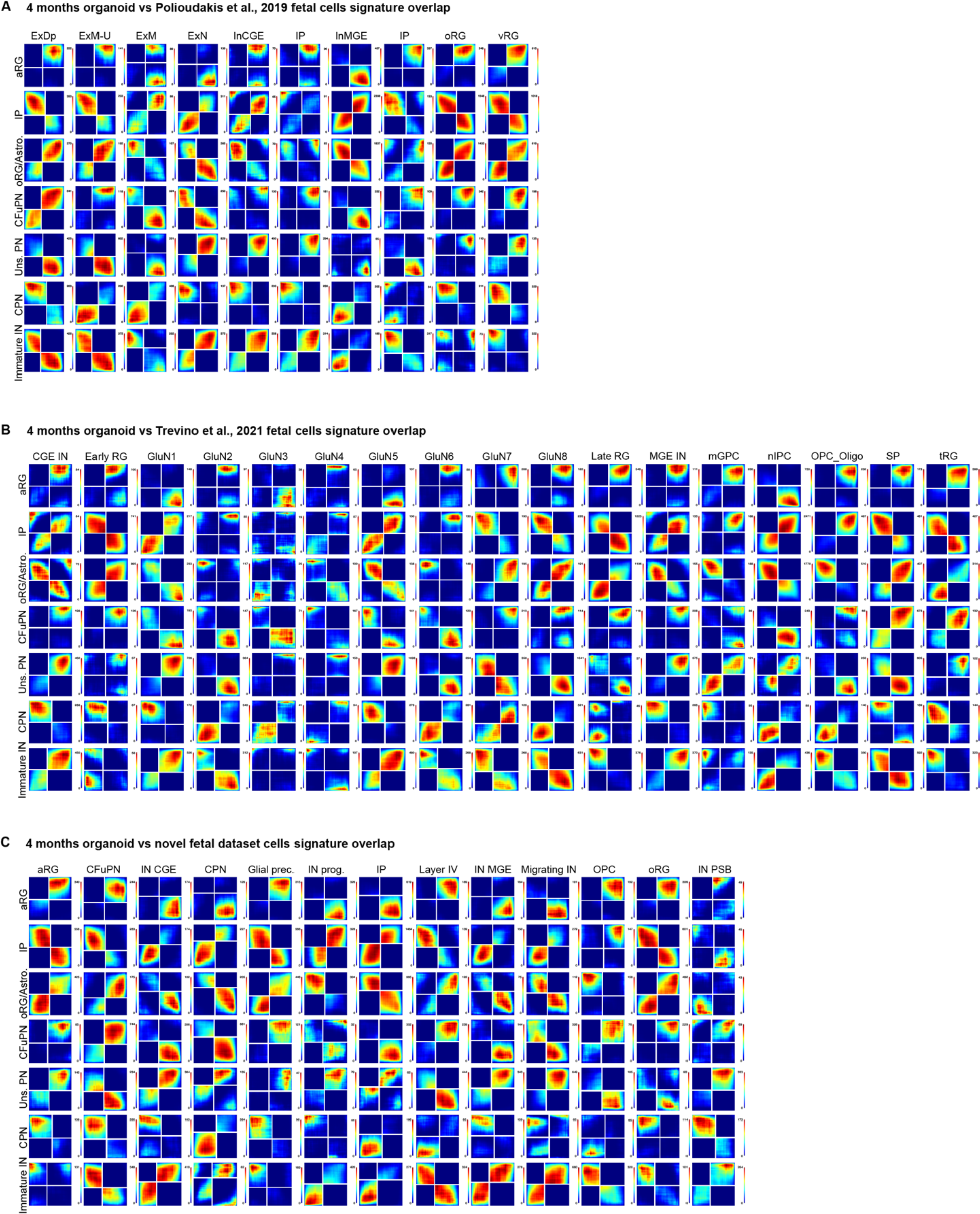
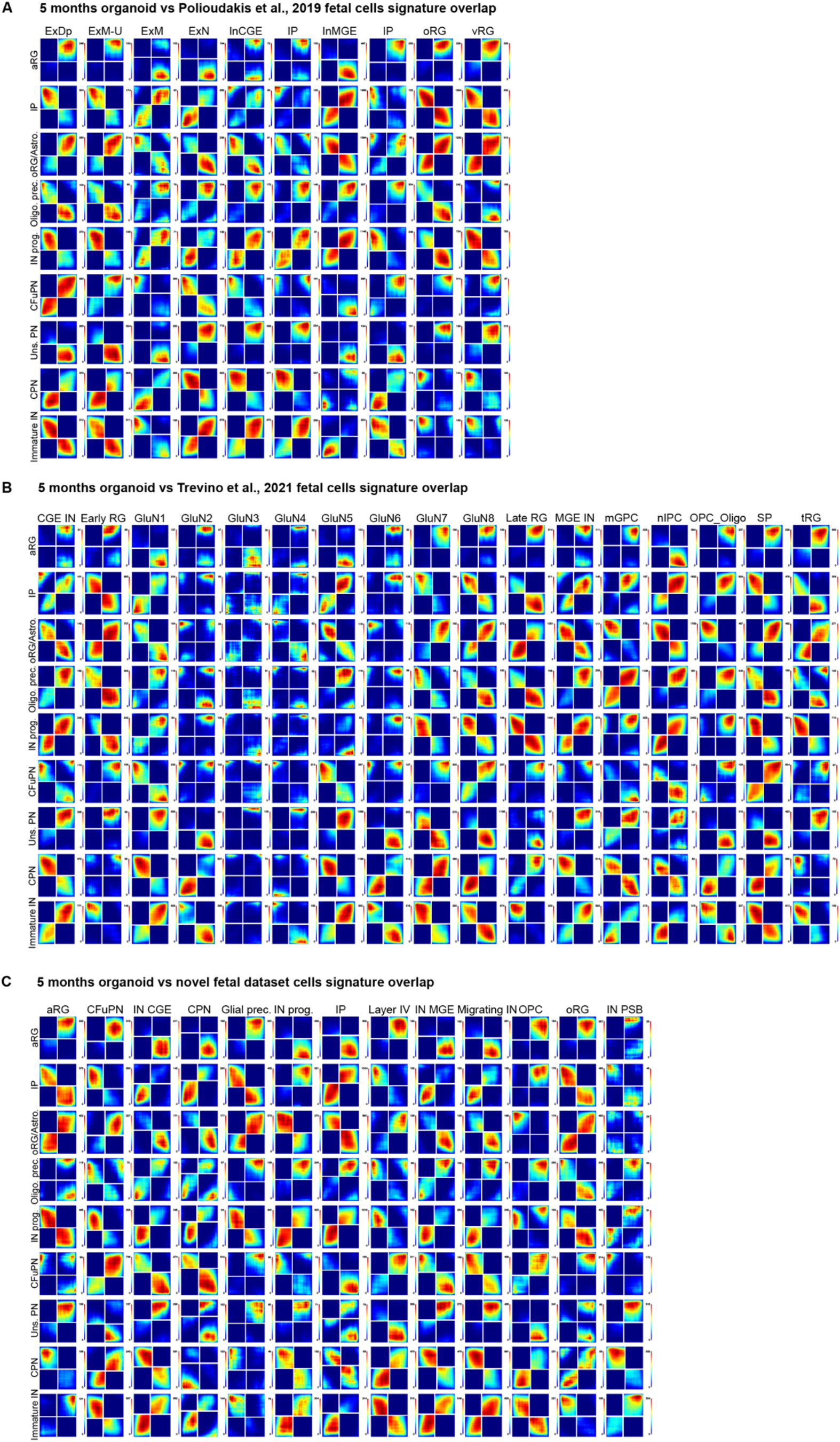
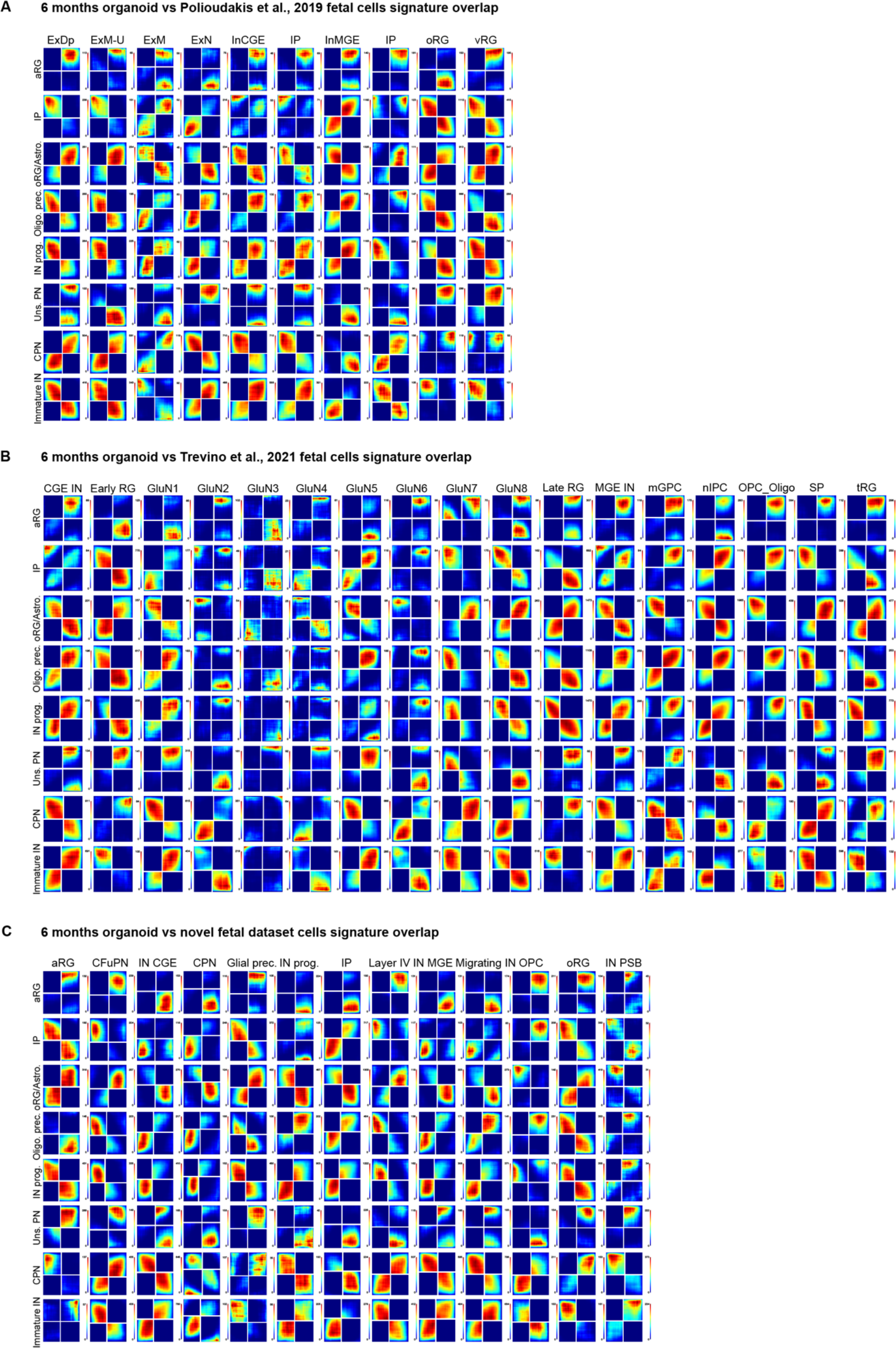
Gene signature concordance of 1-to 6-month organoid cortical cell types and fetal cells. (A-C) Rank-rank hypergeometric overlap (RRHO2) plots from comparing the signatures of organoid cortical cell types (along the x axis) to equivalent signatures from fetal cell types (along the y axis) from (Polioudakis *et al*., 2019) (A), (Trevino *et al*., 2021) (B), and novel fetal dataset (C). Points in the plot are colored by the p value of hypergeometric tests measuring the significance of overlap of gene lists up to that point in the lists.

**Figure S15, related to Figure 3:**
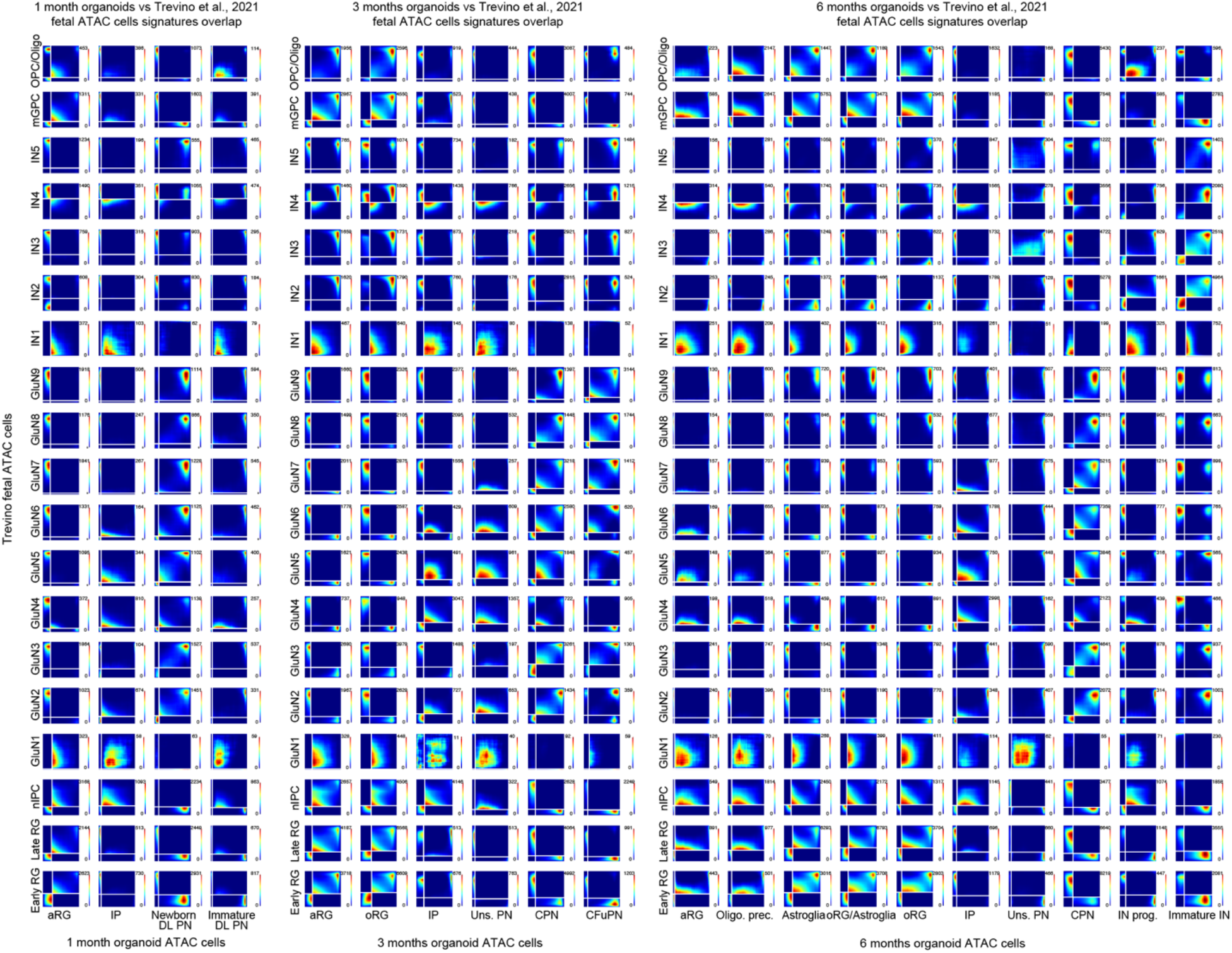
ATAC signature concordance of 1-, 3- and 6-month organoid cortical cell types and Trevino et al. fetal cells. (A-C) Rank-rank hypergeometric overlap (RRHO2) plots from comparing the scATAC-seq signatures of 1- (A), 3- (B), and 6-month(C) organoid cortical cell types (along the x axis) to equivalent signatures from fetal cell types (along the y axis) from (Trevino *et al*., 2021). Signatures were calculated as lists of chromatin regions from a jointly-calculated set of peaks for each timepoint. Points in the plot are colored by the p value of hypergeometric tests measuring the significance of overlap of peak lists up to that point in the lists.

**Figure S16, related to Figure 4:**
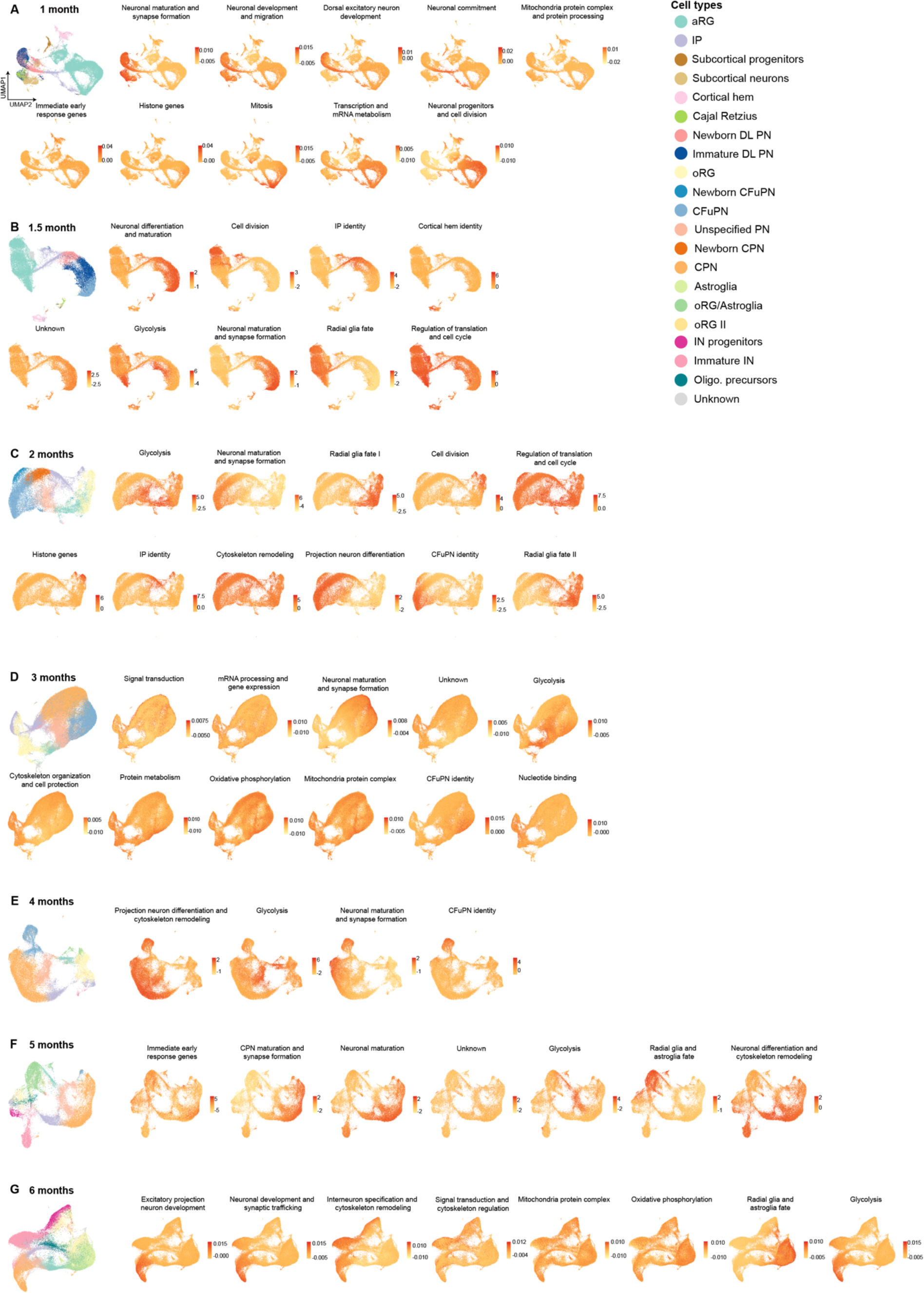
WGCNA modules in organoid scRNA-seq datasets and cell-type-specific metabolic gene set enrichment. (A-G) Feature plots showing eigengene scores for each module resulting from Weighted Gene Correlation Network Analysis (WGCNA; Table S11) on the scRNA-seq datasets from organoids after 1 month (A), 1.5 months (B), 2 months (C), 3 months (D), 4 months (E), 5 months (F), and 6 months (G) in culture. Each module was assigned a name based on manual curation of the gene lists. On the left, cell type UMAPs are repeated from Figure 1A for ease of comparison. 3 and 6 months glycolysis module Feature Plot is repeated from Figure 4A, B for ease of comparison.

**Figure S17, related to Figure 4:**
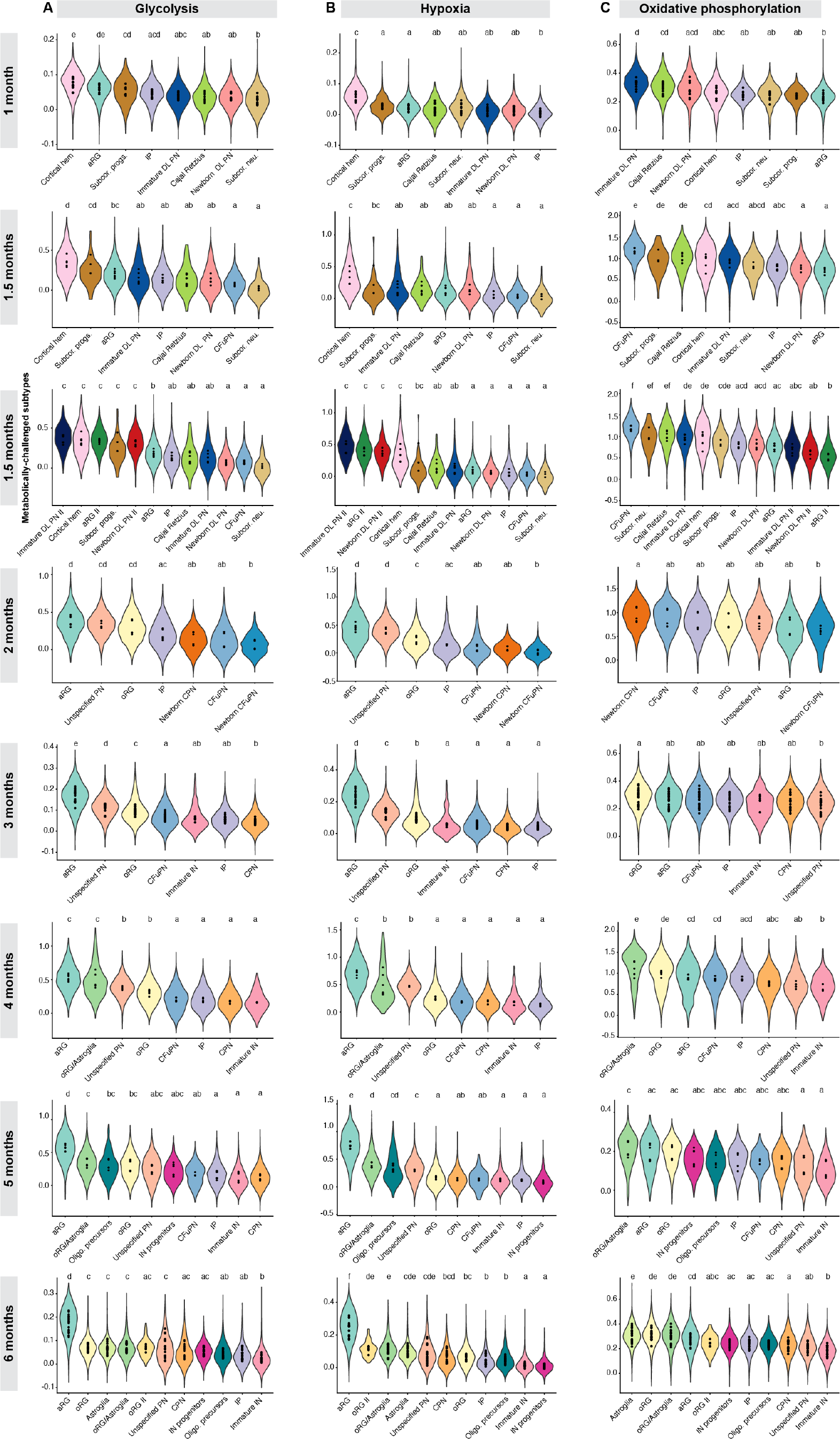
Human cortical organoid cell-type-specific metabolic gene set enrichment. (A) Violin plots showing the distribution of module scores for the MSigDB Hallmark Glycolysis gene set across cell types in 1 to 6 months organoids. Points indicate average scores for each individual organoid. Letters above violin plots indicate the results of a one-way ANOVA followed by pairwise TukeyHSD comparisons; cell types with the same letter indicate there is no significant difference between organoid averages of those cell types. 3 and 6 months organoids Violin Plots are repeated from Figure 4 C-D for ease of comparison. (B-C) Violin plots showing the distribution of module scores for the MSigDB Hallmark Hypoxia (B) and Oxidative phosphorylation (C) gene set across cell types at each timepoint. Letters and points as in A.

**Figure S18, related to Figure 4:**
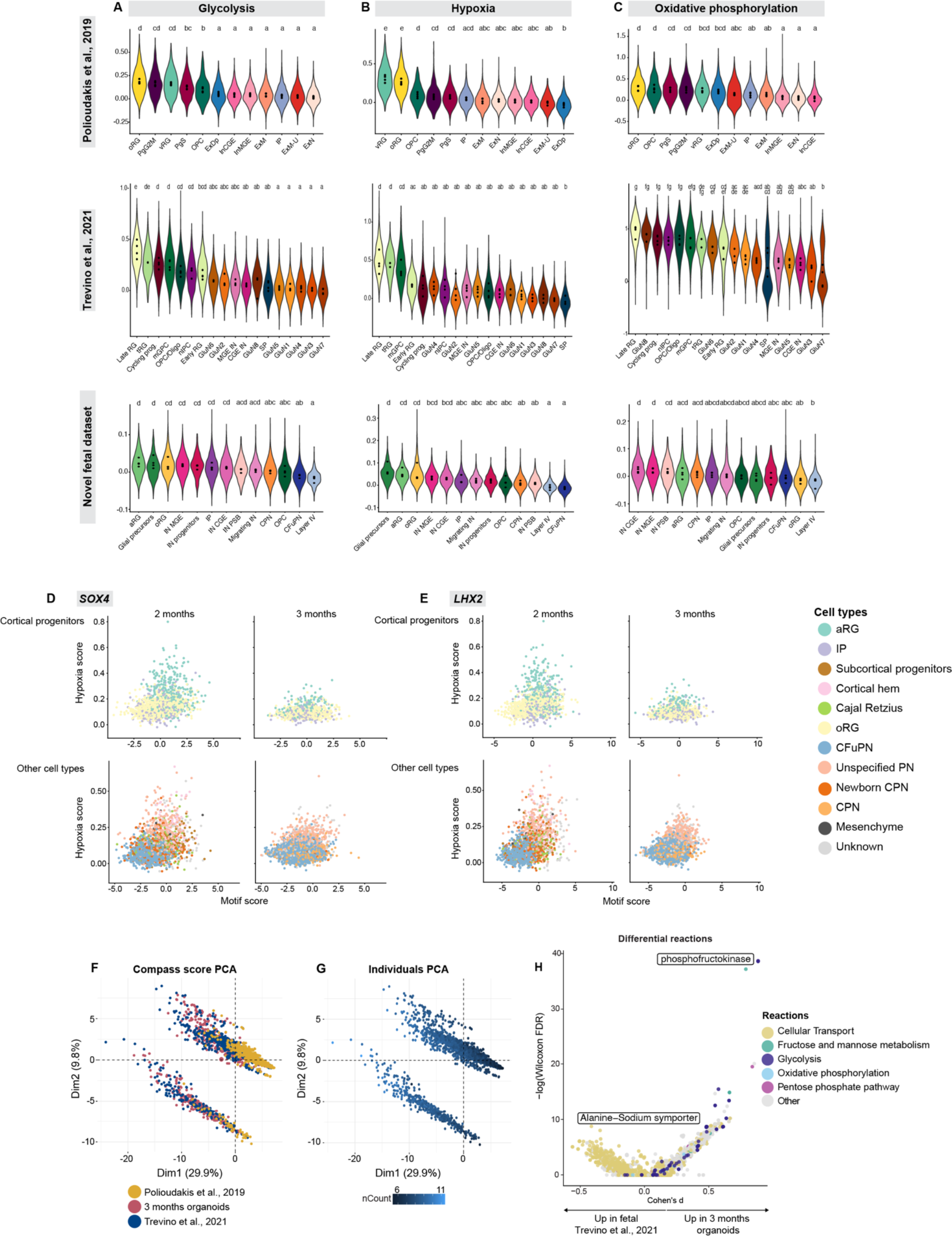
Human fetal cell-type-specific metabolic gene set enrichment and metabolic flux analysis on organoids and human fetal cells. (A) Violin plots showing the distribution of module scores for the MSigDB Hallmark Glycolysis gene set across cell types from (Polioudakis *et al*., 2019) (top), (Trevino *et al*., 2021) (middle), and novel fetal dataset (bottom). Points indicate average scores for each individual organoid. Letters above violin plots indicate the results of a one-way ANOVA followed by pairwise TukeyHSD comparisons; cell types with the same letter indicate there is no significant difference between organoid averages of those cell types. Hallmark Glycolysis Violin Plots are repeated from Figure 4E for ease of comparison. (B-C) Violin plots showing the distribution of module scores for the MSigDB Hallmark Hypoxia (B) and Oxidative phosphorylation (C) gene set across cell types at each timepoint. Letters and points as in A. (D-E) Correlation of *SOX4* (D) and *LHX2* (E) transcription factors’ motif accessibility with module scores from the MSigDB Hallmark Glycolysis and Hypoxia gene sets, from the SHARE-seq data on 2- (left) and 3-month (right) organoids. See Table S14. (F) Principal component analysis (PCA) of the Compass matrix of metabolic reaction potential activity scores, for principal components 1 and 2. Cells are colored by the dataset that they are derived from. (G) PCA as in D, cells are colored by the amount of UMIs sequenced in each cell. (H) Compass differential metabolic activity test between 3-month organoid cells and human fetal cells from (Trevino *et al*., 2021). Cells from each dataset were downsampled to 50 cells of each broad cell type (see Methods) and Compass scores were submitted to a Wilcoxon differential test. Reactions are colored by the subsystem they are assigned in the RECON2 database (Swainston *et al*., 2016).

**Figure S19, related to Figure 4:**
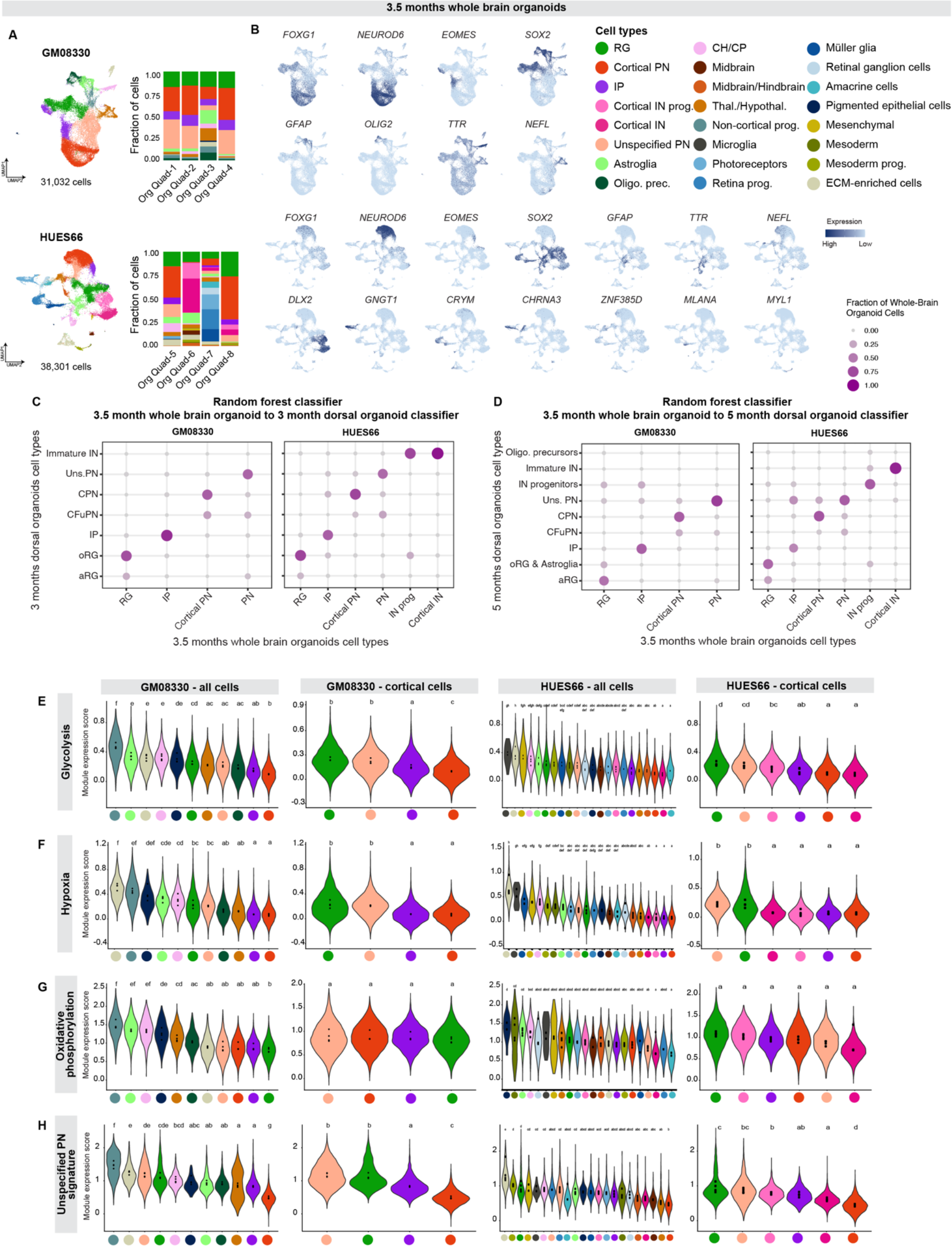
Whole brain organoids classification and cell-type-specific metabolic gene set enrichment. (A) scRNA-seq of whole brain organoids grown with (Quadrato *et al*., 2017) protocol and cultured for 3.5 months. On the top, differentiation batch grown with the GM08330 line. On the bottom, differentiation batch grown with the HUES66 line. Data is represented by UMAPs (left). Cells are colored by cell type, and the number of cells per plot is indicated. Right, proportion of cells belonging to each cell type from each individual organoid. (B) Feature plots showing normalized expression of marker genes of cell types across organoids, plotted on the same UMAPs from A: telencephalon (*FOXG1*), dorsal excitatory neurons (*NEUROD6*), IP (*EOMES*), progenitors (*SOX2*), astroglia (*GFAP*), cortical hem/choroid plexus (CH/CP, *TTR*), thalamus (Thal., *NEFL*), interneurons (*DLX2*), retina photoreceptors (*GNGT1*), retinal Müller glia (*CRYM*), retinal ganglion cells (*CHRNA3*), amacrine cells (*ZNF385D*), mesoderm (*MYL1*). (C) Assignments of 3.5 months whole brain organoid cortical cell types to 3-month dorsal organoid cell types, as classified by random forest. Points are sized and colored by the fraction of each whole brain organoid cell type assigned to the dorsal organoid cell types. (D) Assignments of 3.5 months whole brain organoid cortical cell types to 5-month dorsal organoid cell types, as classified by random forest. Points are sized and colored by the fraction of each whole brain organoid cell type assigned to the dorsal organoid cell types. (E) Violin plots showing the distribution of module scores for the MSigDB Hallmark Glycolysis gene set across all whole brain organoid cell types (left column), or cortical cell types (right column), for the GM08330 and HUES66 differentiation batches. Points indicate average scores for each individual organoid. Letters above violin plots indicate the results of a one-way ANOVA followed by pairwise TukeyHSD comparisons; cell types with the same letter indicate there is no significant difference between organoid averages of those cell types. (F-G) Violin plots showing the distribution of module scores for the MSigDB Hallmark Hypoxia (F) and Oxidative phosphorylation (G) gene set across cell types at each timepoint. Letters and points as in E. (H) Violin plots showing the distribution of module scores for the 3 months dorsal organoids ‘unspecified PN’ molecular signature (Table S3) across whole brain organoid cell types at each timepoint. Letters and points as in E.

**Figure S20, related to Figure 4:**
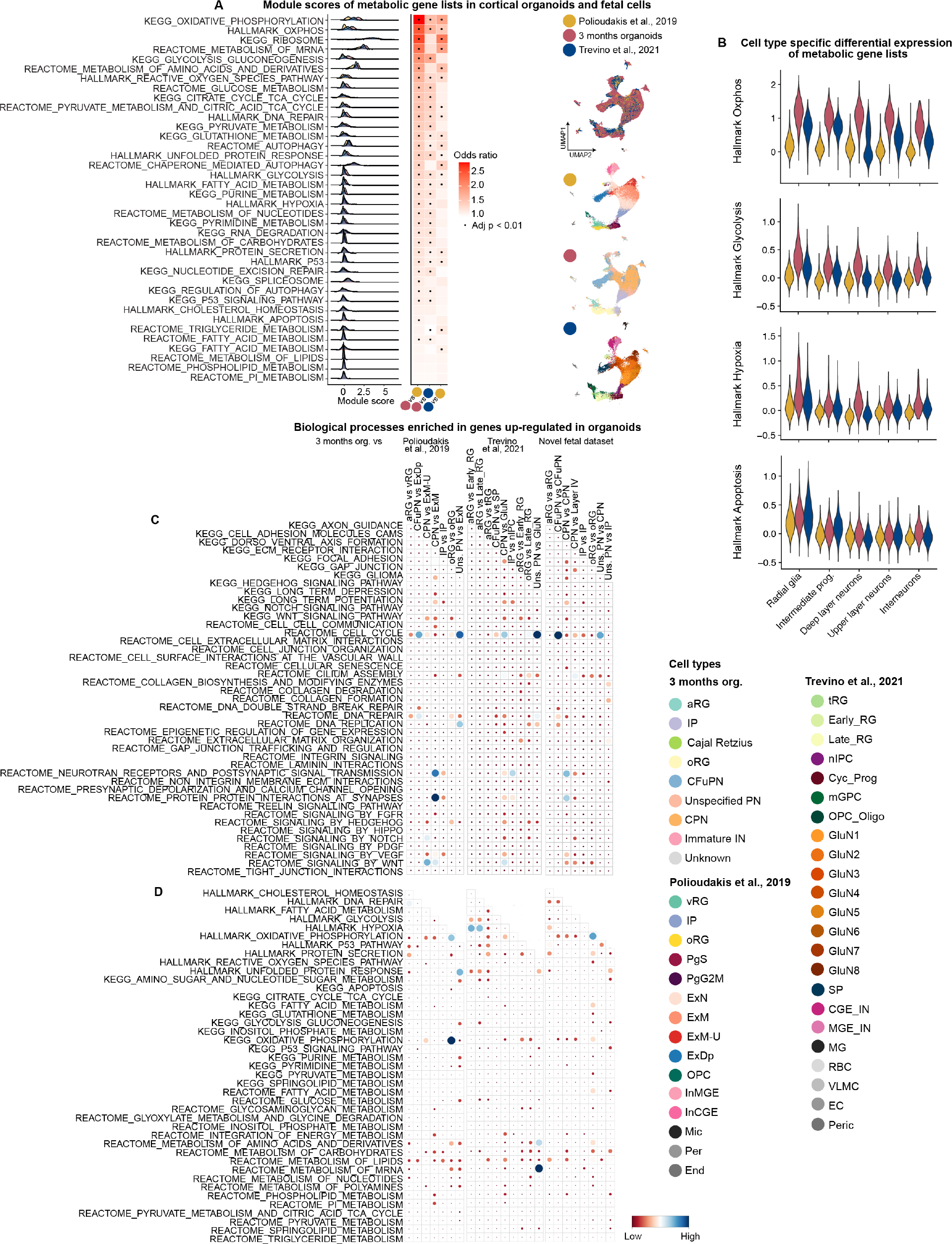
Differential metabolic pathway expression in human organoids and fetal cells. (A) Module scores of 38 metabolic gene lists in 3 months organoids (red) and fetal cells ((Polioudakis *et al*., 2019) yellow, (Trevino *et al*., 2021) blue). Left: ridge plot showing the distribution of module scores of the metabolic gene lists in the three datasets. Middle: heatmap indicating the odds ratio between 3 months organoids versus (Polioudakis *et al*., 2019), 3 months organoids versus (Trevino *et al*., 2021), and (Trevino *et al*., 2021) versus (Polioudakis *et al*., 2019) The odds ratio was calculated by computing a linear mixed-effect model, modeling the module scores in the two datasets being compared, with dataset as a fixed effect and individual sample as a random effect. The odds ratio is the exponential of the dataset coefficient of the resulting model. Tiles are outlined in black if the p value of an ANOVA between this model and a model without the dataset coefficient was less than 0.01. Right: UMAP of the three datasets after running Harmony batch correction. (B) Module score distributions for 4 metabolic gene lists by cell type in 3 months organoids (red) and fetal cells ((Polioudakis *et al*., 2019) yellow, (Trevino *et al*., 2021) blue). For the purpose of this analysis, cells were re-grouped into equivalent broad cell categories (see Methods). (C-D) Overlap of genes up-regulated in organoids at 3 months according to RRHO2 analysis (upper left quadrant) with published gene sets. 43 biological processes were queried (C) along with 38 metabolic gene lists (D). Dots are sized and colored to -log_10_(adjusted p value) of a hypergeometric test comparing the two lists.

**Figure S21, related to Figure 4:**
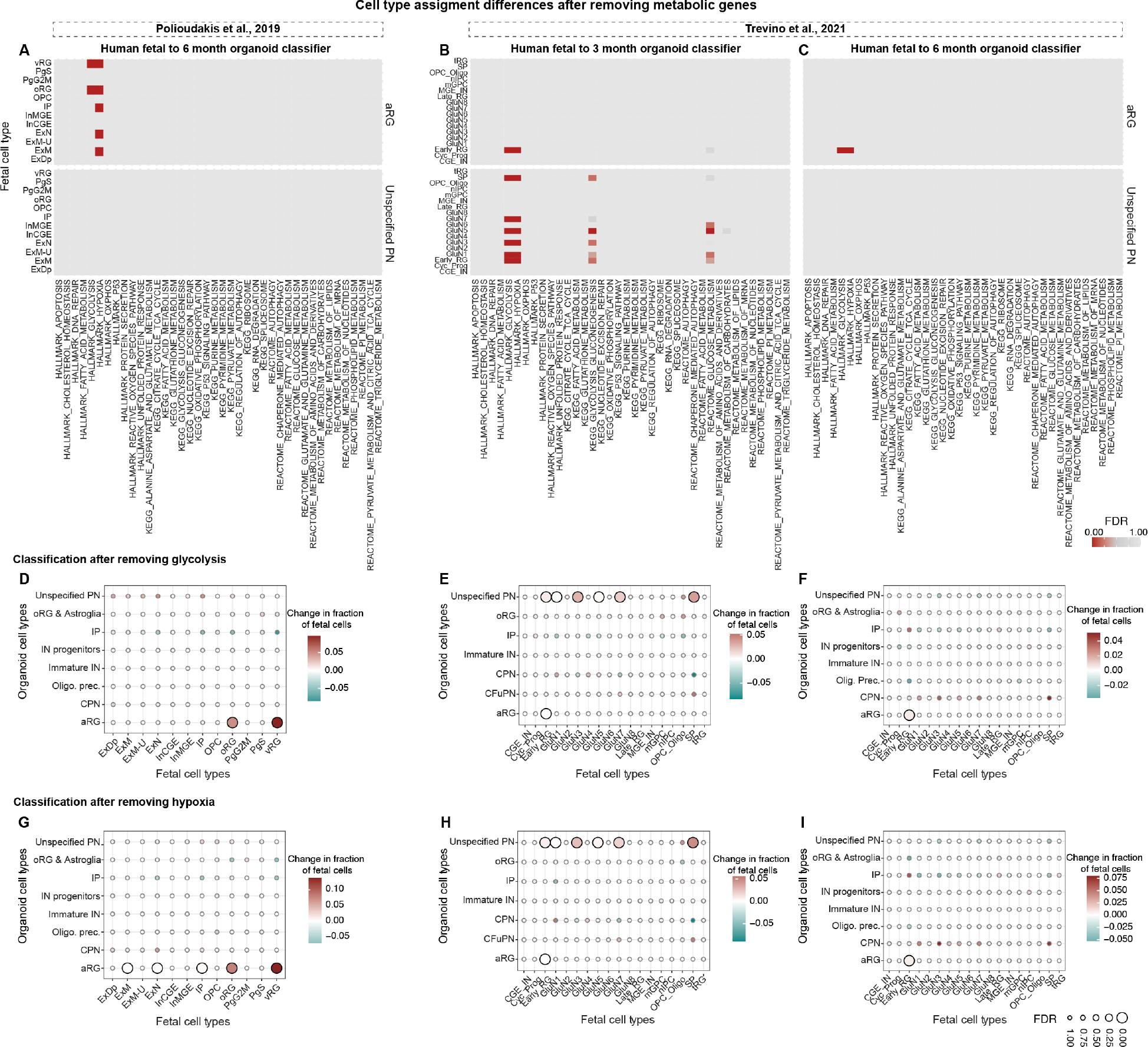
Effects of metabolic pathways on transcriptional matching. (A-C) Significance (FDR by Benjamini-Yekutieli) of an increase in (Polioudakis *et al*., 2019) fetal cell (A) and (Trevino *et al*., 2021) fetal cell (B-C) assignments to the 6-month (A,C) and 3-month (B) organoid aRG label (top) and unspecified PN label (bottom) after removing genes from the model in the 38 MSigDB metabolic gene sets listed along x axis. (D-F) Change in the assignments of (Polioudakis *et al*., 2019) fetal cell types to 6-month organoid cell types (D) and (Trevino *et al*., 2021) fetal cell types to 3 and 6-month organoid cell types (E, F), after removing genes from the model that belong to the MSigDB Hallmark Glycolysis gene set. Points are colored by the change in the fraction of cells assigned to each label by the model with removed genes, compared to the full model. Points are sized by the significance of an increase in that fraction (FDR by Benjamini-Yekutieli), compared to a background distribution of 10,000 models trained after removing random gene lists. Points with FDR < 0.05 are outlined in black. As in text, changes were considered significant if they had an FDR < 0.05 and changed at least 1% of assigned cells. (G-I) Change in the assignments of (Polioudakis *et al*., 2019) fetal cell types to 6-month organoid cell types (G) and (Trevino *et al*., 2021) fetal cell types to 3 and 6-month organoid cell types (H, I), after removing genes from the model that belong to the MSigDB Hallmark Hypoxia gene set. Points are colored and sized as in D-F.

**Figure S22, related to Figure 5:**
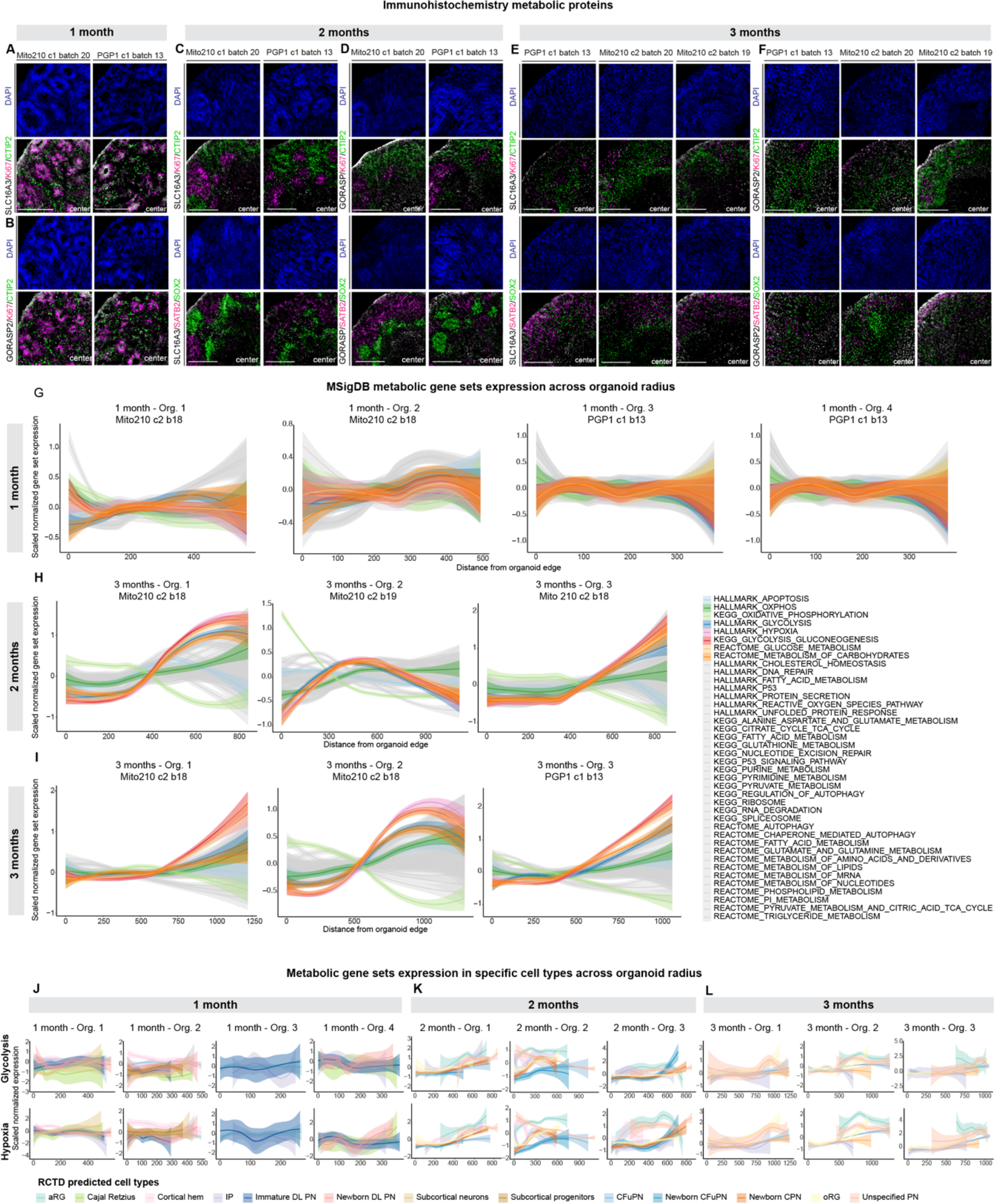
Spatial expression of metabolism-related genes and proteins. (A-F) Immunohistochemistry of dorsal forebrain progenitors (SOX2), cycling progenitors (Ki67), CFuPN (CTIP2), CPN (SATB2) markers, as well as SLC16A3 (A, C, and E) and GORASP2 (B, D, and F) in 1- (A and B), 2- (C and D), and 3-month (E and F) organoids. Scale bar 200 μm (A-B), 500 μm (C-F). (G-I) Distribution of the Slide-seqV2 beads’ scaled expression for each gene set compared to their distance from the edge of the organoid. Solid line shows the smoothed conditional mean values across beads, with transparent backgrounds showing the 95% confidence interval. Distributions are shown separately for each organoid at 1 month (G), 2 months (H), and 3 months (I). The plots for 2-month Organoid 1 and 3-month Organoid 1 are repeated from Figure 5 for ease of comparison. (J-L) Distributions as in G for the Hallmark Glycolysis (top) and Hallmark Hypoxia (bottom) gene sets, graphed separately for beads assigned to each cell type. Cell types assigned to more than 8 beads per organoid are shown. Distributions are shown separately for each organoid at 1 month (J), 2 months (K), and 3 months (L). The plots for 2-month Organoid 1 and 3-month Organoid 1 are repeated from Figure 5 for ease of comparison.

**Figure S23, related to Figure 5:**
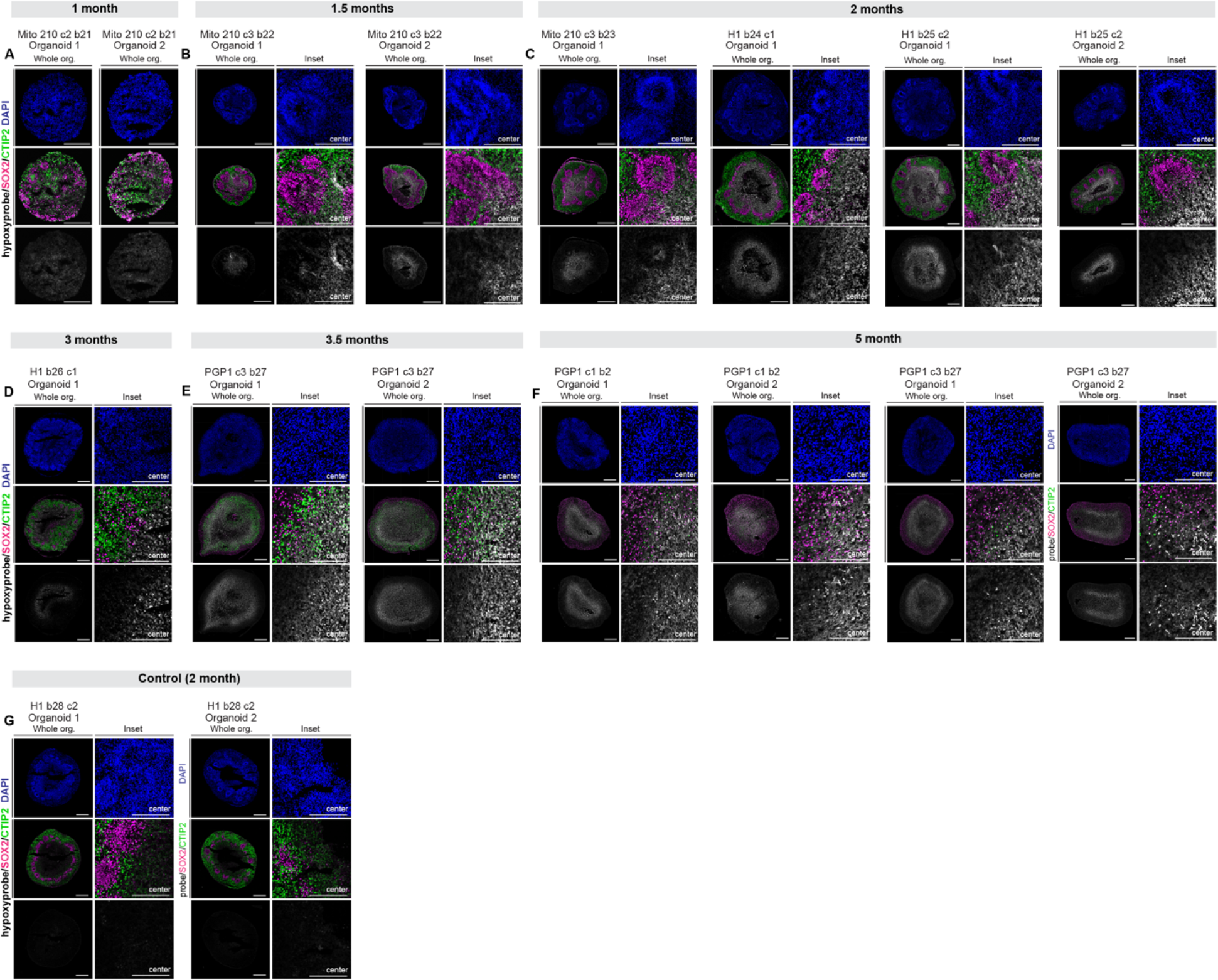
Spatial expression of hypoxia probe. (A-F) Immunohistochemistry of hypoxyprobe treated 1 month (A), 1.5 months (B), 2 months (C), 3 months (D), 3.5 months (E) and 5 months (F) organoids with markers for dorsal forebrain progenitors (SOX2), CFuPN (CTIP2), and hypoxyprobe. Scale bar whole organoids (org.) 200 μm 1 month, 500 μm 1.5 months onward. Scale bar insets 200 μm. (G) Immunohistochemistry for negative controls, performed on 2-month organoids. An immunohistochemistry using the primary antibody to detect the hypoxyprobe was performed in slices from organoids not incubated with the hypoxyprobe. Scale bars as in A-F.

**Figure S24, related to Figure 6:**
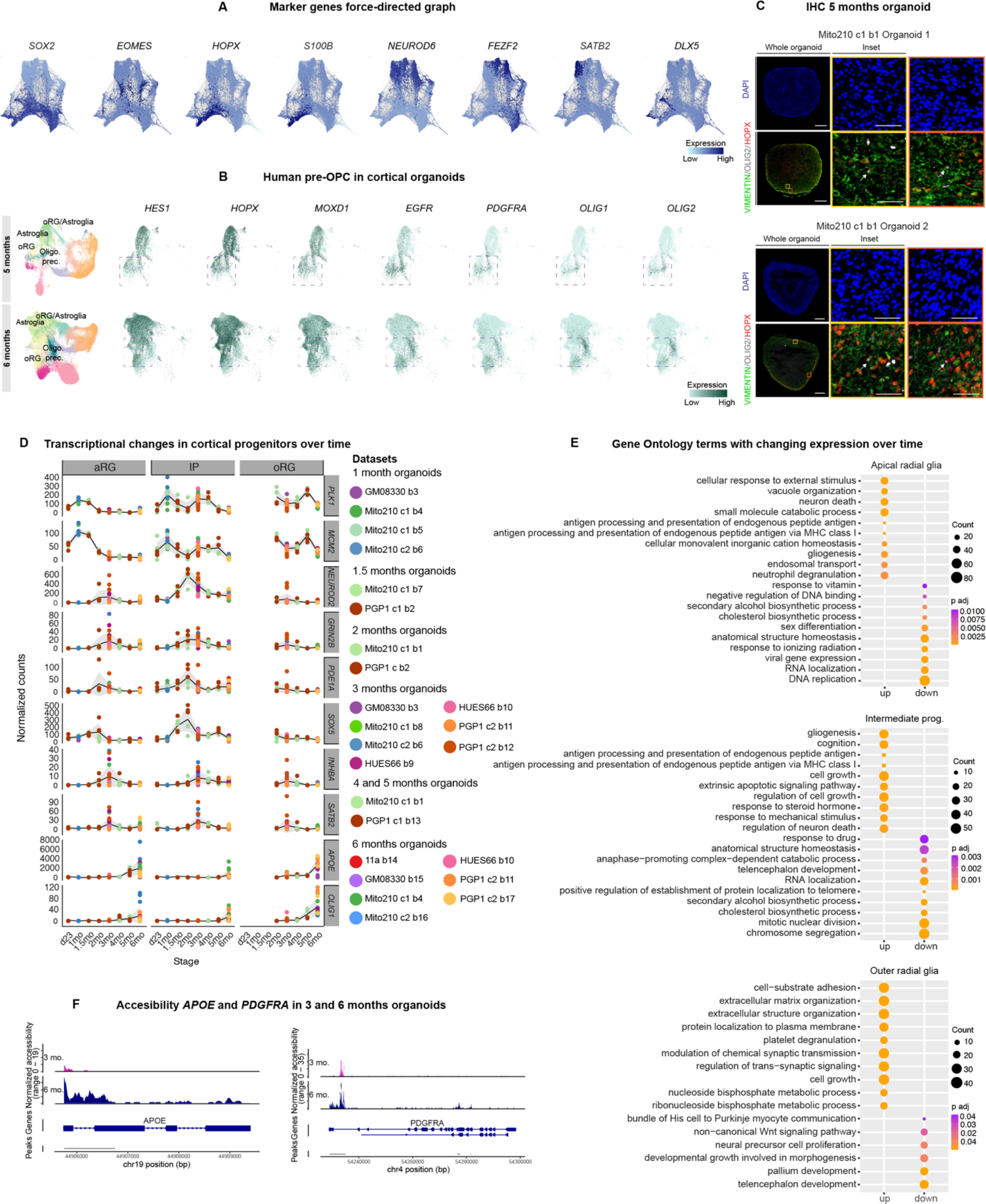
Developmental trajectories and progenitors’ transcriptional changes across time in human cortical organoids. (A) Feature plots showing normalized expression of marker genes of cortical cell types plotted on the same force-directed graph from Figure 6A: aRG and oRG (*SOX2*), IP (*EOMES*), oRG *(HOPX*), astroglia (*S100B*), dorsal excitatory neurons (*NEUROD6*), CFuPN (*FEZF2*), CPN (*SATB2*), interneurons (*DLX5*). (B) Left, scRNA-seq of organoids cultured for 5 and 6 months represented by UMAPs of the data after Harmony batch correction, repeated from Figure 1 for ease of comparison. Right, Feature plots showing normalized expression of marker genes of cortical pre-oligodendrocyte precursor cells in oRG, oRG/Astroglia, Astroglia and Oligodendrocyte precursors. Gray dotted boxes indicate regions of pre-OPC marker expression. (C) Immunohistochemistry of human pre-OPC markers (VIMENTIN, HOPX, OLIG2) in 5-month organoids. Whole organoids are displayed on the left, where colored squares refer to the insights on the right. White arrows indicate cells in which the three markers colocalize. Scale bar whole organoids 500 μm. Scale bar insets 50 μm. (D) Gene ontology terms associated with up-regulated or down-regulated genes across organoid development (1 to 6 months). Dot size indicates gene count and color indicates adjusted p value. (E) Normalized counts of marker genes for biological processes or cell types (Cell proliferation: *PLK1*, *MCM2*; Dorsal excitatory neurons: *NEUROD2*; Synapses: *GRIN2B*; CFuPN: *PDE1A*, *SOX5*; CPN: *INHBA*, *SATB2*; Astroglia: *APOE*; Oligodendrocyte precursors: *OLIG1*) across the different timepoints sampled through organoid development, d23 (day 23) to 6 mo (6 months). Each colored dot represents the normalized counts within a differentiation batch. (F) Chromatin accessibility (scATAC-seq read coverage) of *APOE* and *PDGFRA* at 3 months (pink) and 6 months (purple).

**Figure S25, related to Figure 6:**
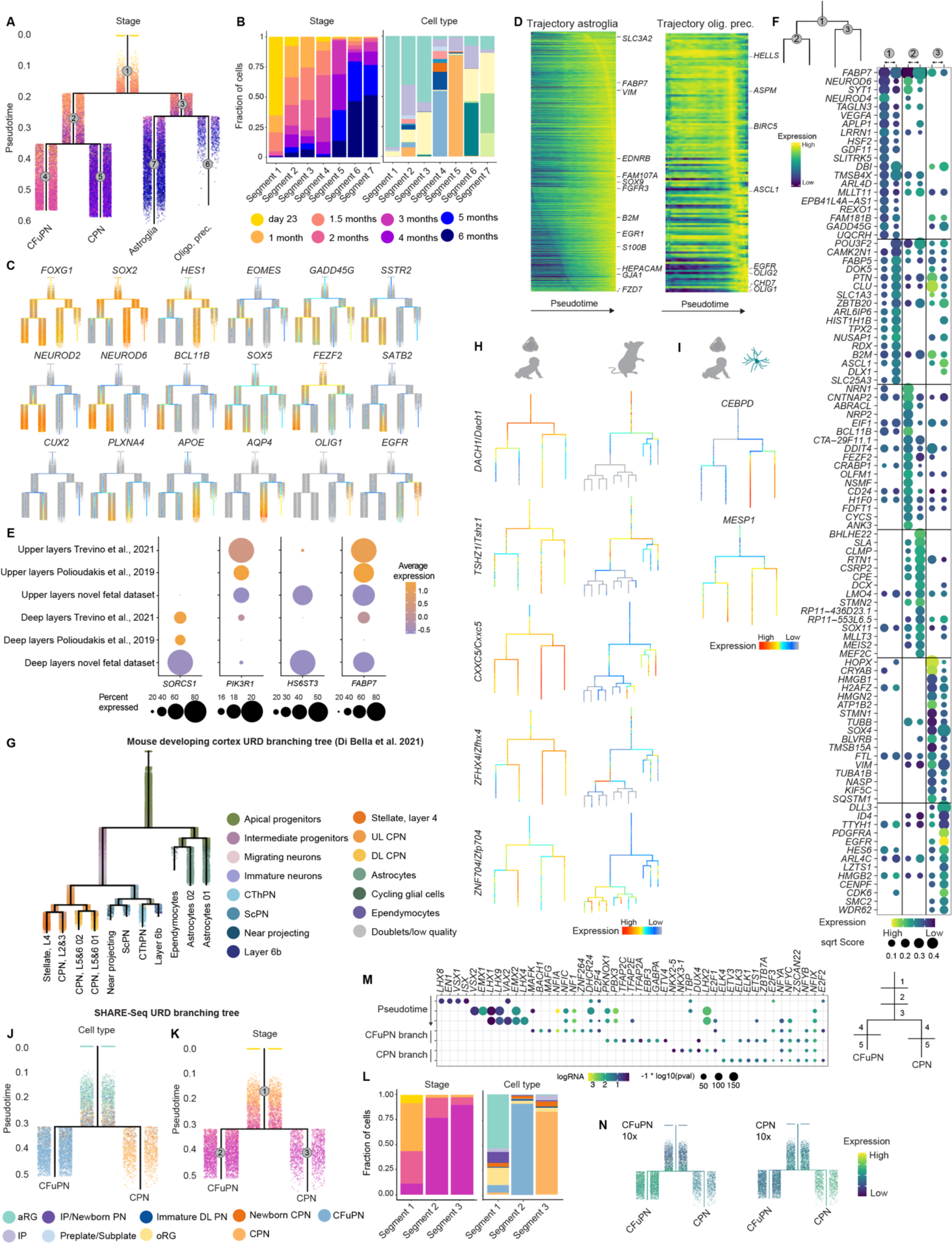
Characterization and gene expression in human cortical organoid branching tree. (A) URD branching tree. Cells are colored according to the organoid stage. Gray circles indicate segment number. (B) Left: Proportion of cells belonging to each organoid stage in each of the URD branching tree segments. Right: Proportion of cells belonging to each cell type in each of the URD branching tree segments. Cell types colored as in Figure 1. (C) Branching tree showing the expression of marker genes of the telencephalon (*FOXG1*), dorsal forebrain progenitors (*SOX2*, *HES1*), intermediate progenitors (*EOMES*), neuronal migration (*GADD45G*, *SSTR2*), dorsal excitatory neurons (*NEUROD2*, *NEUROD6*), CFuPN (*BCL11B*, *SOX5*, *FEZF2*), CPN (*SATB2*, *CUX2*, *PLXNA4*), astroglia (*APOE*, *AQP4*), oligodendrocyte precursors (*OLIG1*, *EGFR*). (D) Heatmap of lineage-specific gene cascades for astroglia and oligodendrocyte precursors. Gene expression in each row is scaled to maximum observed expression. Genes are ordered based on their onset and peak expression timings. Full cascade is shown in Table S15. (E) Expression of lineage-specific genes from the organoid pseudotime tree in human fetal datasets. Average expression is scaled per gene (F) Top 20 genes per branch predicted to be involved in cell type divergence. Points are sized by their score, defined as their feature importance in a gradient-boosting classifier distinguishing cells in each segment of the branch point, and are colored by their average expression in the corresponding cells (row-scaled). (G) URD branching tree of the mouse developing cortex (embryonic day 10.5 to postnatal day 4) adapted from (Di Bella *et al*., 2021). (H) Expression of transcription factors in simplified branching tree predicted as important in the first (*DACH1*, *TSHZ1*), second (*ZNF704*) and third branch points (*CXXC5*, *ZFHX4*). Left: expression on the human organoid URD branching tree, right: expression on the developing mouse cortex URD branching tree (adapted from (Di Bella *et al*., 2021)). (I) Expression of *CEBPD* and *MESP1* in branching tree. These genes are predicted as important in the third branch point. (J) SHARE-Seq URD branching tree. Cells are colored according to cell types. (K) SHARE-Seq URD branching tree. Cells are colored according to the organoid stage. Gray circles indicate segment number. (L) Left: Proportion of cells belonging to each organoid stage in each of the SHARE-Seq URD branching tree segments. Right: Proportion of cells belonging to each cell type in each of the SHARE-Seq URD branching tree segments. (M) Left, Transcription factor motifs enriched in DARs along segments of the SHARE-seq URD tree. Top three rows indicate 3 equal divisions of the parental segment, while CPN and CFuPN branch rows indicate two equal divisions of each of their segments. Right, schematic representation of segments in a simplified SHARE-Seq URD tree. (N) Module score expression of CFuPN and CPN lineage associated genes of 10x scRNA-seq branching tree in SHARE-Seq branching tree.

**Figure S26, related to Figure 7:**
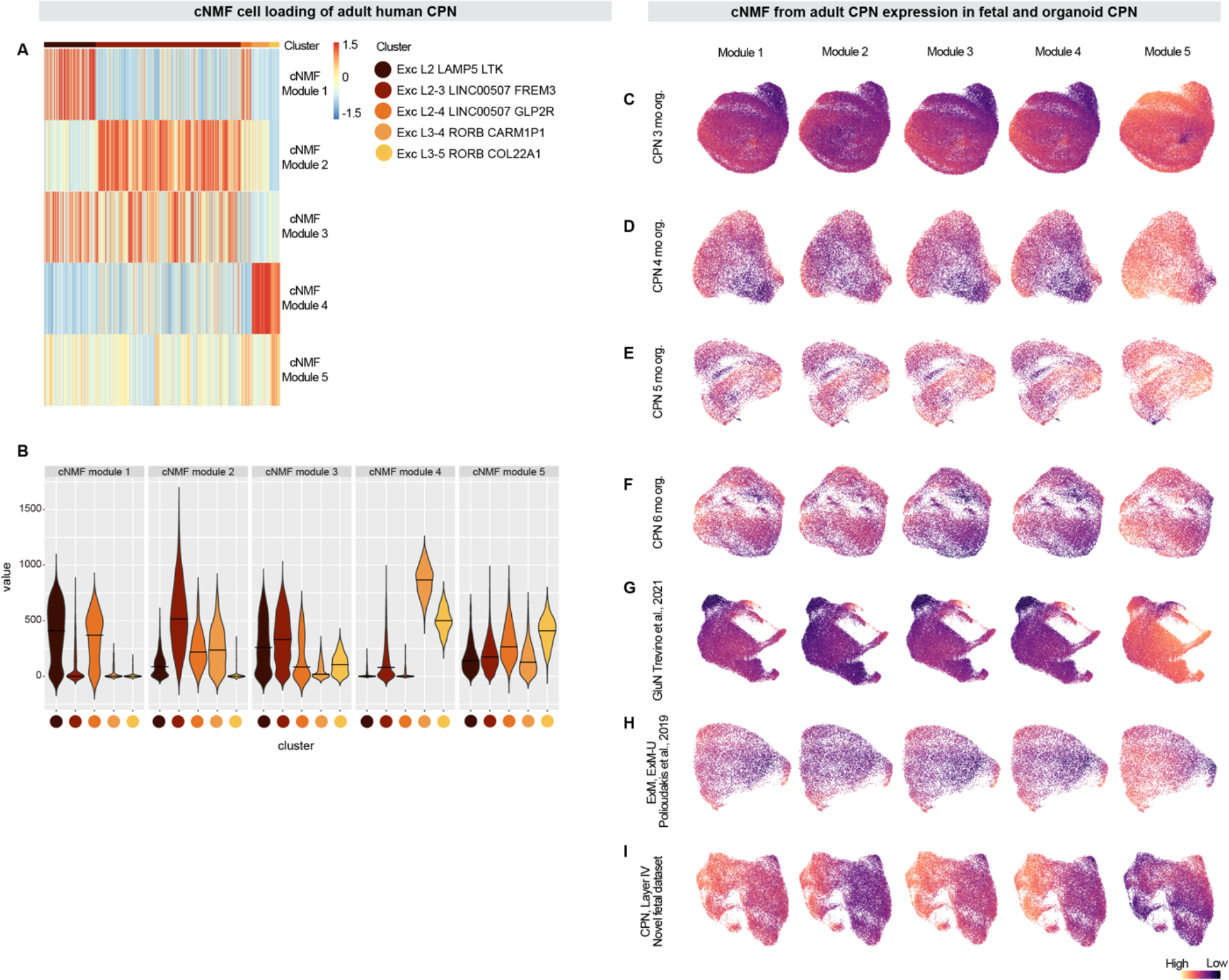
Gene modules in adult callosal projection neurons and expression in organoid and fetal cells. **(A-B)** Heatmap (A) and violin plot (B) showing the cell loadings of the five gene modules obtained from cNMF in adult CPN (Hodge *et al*., 2019). **(C-I)** UMAP of 3 month (C), 4 month (D), 5 month (E), 6 month (F) organoid (org.) callosal projection neurons (CPN) and human fetal GluN from (Trevino *et al*., 2021) (F), ExM and ExM-U from (Polioudakis *et al*., 2019) (H), and layer IV CPN from the novel fetal dataset (I) generated using as variable features the genes obtained in adult CPN cNMF analysis. The UMAPs are colored by the module scores of genes in each consensus Non-negative Matrix Factorization (cNMF) module.

## Supplementary Text

### Developmental-stage and cell-type-appropriate expression of known endogenous markers in scRNA-seq dataset

We determined gene expression signatures for all cortical cell types in organoids at each stage, using a pseudo-bulk analysis with a model that controls for variability between organoids (see Methods). We examined the top 15 DEGs for each cell type to confirm appropriate expression of known endogenous markers.

At 1 month, the molecular signature of neuronal progenitors (aRG and IP) contained genes related to their proliferative state as well as identity within the top 15 DEGs (e.g., *HMGA2*, *GLI3, SFRP1* for aRG; *ELAVL4*, *NHLH1*, *HES6* for IP; Figures S5A, Table S4). At 1.5 months, when newborn DL PN emerge in organoids, we observe canonical markers of neuronal commitment in their signatures (top 15 DEGs include *NHLH1*, *SOX11,* Figure S5H, Table S5). At 2 months, when organoid oRG appeared, their molecular signature reflected canonical oRG markers (top 15 DEGs include *FABP7, PTN, ETV5*; Figure S5C, Table S2), which were also consistently observed in their 3-month molecular signature (Figure 3A, Table S3). In 3-month organoids, when the main excitatory neuron types have emerged, the molecular signatures of organoid CFuPN and CPN populations contained genes characteristic of *bona fide* endogenous CFuPN and CPN. For example, among the top 15 DEGs, we detected genes such as *EFNA5, ETV1*, and *LMO7*, characteristic of CFuPN, and *CACNA2D1, DPYSL3*, and *PTPRK*, previously identified as highly expressed in CPN (Klingler *et al*., 2019) (Figure 3B, Table S3). Both at 3 and 4 months, when we observed the highest diversity of excitatory neuron subtypes in organoids, we found the molecular signature of CFuPN and CPN to be largely mutually exclusive, indicating that in organoids, these two main subclasses of excitatory neurons are molecularly distinct, as observed *in vivo* (Molyneaux *et al*., 2015) (Figures 3B and S5J).

At 4 months, the appearance of astroglia was reflected in the oRG and astroglia molecular signature, where the top 15 DEGs contained known astrocyte markers (e.g., *S100B, GFAP, APOE,* Figure S5D, Table S6). In 5-month organoids, when interneurons are an emerging and abundant population, genes known to mediate interneuron specification and migration were found within the signature for the immature IN class (*DLX5*, *ARX,* Figure S5K, Table S7). Finally, at 6 months, when oRG, astroglia, and interneuron populations have become more prominent, the signatures for these populations contained genes characteristic of their *in vivo* identity (e.g., *PEA15*, *F3*, *TNC* for oRG & astroglia; *KAT6B* for Immature IN) (Figure S5F, S5L, Table S8).

The expression of the 200 most differentially expressed genes of these signatures within their respective cell types was consistent across individual organoids; there was no statistically significant difference in the means of any cell-type signature gene set across all organoids assayed at each timepoint (Figures 3A-B, S5; one-way ANOVA, p > 0.05).

### Enrichment of developmental-stage and cell-type appropriate transcription factors

We used the epigenetic signatures of each cell type in the organoids to identify TFs associated with individual cell types. We searched for *de novo* motifs (see Methods; (Heinz *et al*., 2010)) that were over-represented in regions that were specifically accessible in each cell type within each timepoint assayed, assigned them to corresponding TFs by binding site similarity, and examined the expression of the corresponding TF mRNAs in the scRNA-seq data from the same cell line and timepoint (Figure 3C, Table S7). Cell types from early organoids (1 month) showed enrichment of putative binding sites for TFs known to regulate early steps of mouse forebrain development, such as *EMX1*, *EMX2*, *LHX2*, and *LHX9* (Yoshida *et al*., 1997; Matsunaga, Araki and Nakamura, 2000; Peukert *et al*., 2011; Chou and Tole, 2019) (Figure 3C, Table S5). At all timepoints analyzed, the neurogenic TF *NEUROG2* was enriched in the majority of neuronal types; as organoids matured (3 and 6 months), we observed widespread enrichment for TFs of the NFI (Nuclear factor one) family (*NFIX*, *NFIA*, *NFIC*), which are known to have broader roles in cortical progenitor proliferation, and neuronal differentiation (Piper *et al*., 2014), and gliogenesis (Kang *et al*., 2012). Likewise, at 3 and 6 months the *SOX9* motif was enriched in oRG and astroglia, in agreement with a role for this gene in oRG proliferation (Güven *et al*., 2020) and gliogenesis (Kang *et al*., 2012). Finally, concomitant with the late production of interneurons, we saw enrichment of DLX TFs at 6 months (Figure 3C).

To determine the accessibility dynamics of the target sites, we computed the difference between the normalized accessibility of a given TF motif across the genome and the normalized expression of the TF itself (see Methods). We found a variety of patterns, including cell populations where target site accessibility and TF expression were synchronized, cases where chromatin remained accessible after the TF was no longer expressed, and cases where the TF was expressed prior to target site opening (Figure S3J), indicating a range of regulatory strategies.

### Concordance between organoid and endogenous fetal cell-type chromatin accessibility profiles

To assess the similarity between the epigenomic landscape of cells in brain organoids and that of their endogenous counterparts, we compared scATAC-seq data from 1, 3, and 6 months *in vitro* organoids to a recently published dataset of scATAC-seq in human cortex at PCW 16-24 (Trevino *et al*., 2021). Both datasets were projected to a shared peak set, computed independently for the three organoid timepoints, and molecular signatures were calculated and compared using RRHO2, as for RNA-seq. We found similar cell type signatures between corresponding cell types (Figure S7E-G, Figure S15). For example, at 6 months, organoid oligodendrocyte precursors resemble fetal OPC/ologidendrocytes. Organoid IPs are in strong agreement with fetal IP/nIPC accessible regions across all timepoints. Organoid CPN agree with fetal GluN clusters expressing CPN markers (i.e., GluN2-7), both at 3 and 6 months *in vitro*. At 6 months, Immature IN and IN progenitors in organoids are in strong agreement with the fetal IN2-4 clusters (expressing markers of the CGE lineages). Indeed, when the datasets were analyzed at similar total number of reads (achieved by downsampling the (Trevino *et al*., 2021) dataset), few DARs were found between the human and the organoid samples (Table S10). Gene Ontology analyses on the sets of genes with accessible peaks unique to one dataset or the other showed only a few significantly enriched terms: “embryonic organ development” and “neural nucleus development”, both enriched in genes accessible in fetal cells only. This analysis demonstrates high concordance between cell-type specific chromatin accessibility profiles in organoid cell types and the corresponding endogenous human fetal cell populations.

### Domains of regulatory chromatin of *FEZF2*

SHARE-seq can directly link regions of chromatin accessibility to the expression of genes in *cis.* We used this method to identify domains of regulatory chromatin (DORCs, (Ma *et al*., 2020)); that is, regions with genes with greater than five associated peaks. Genes with a large number of significant peak-gene associations included several TFs of known importance during neurodevelopment such as *ZIC1*, *NEUROD2*, *SOX2*, and *LHX2*, indicating high precision epigenetic control of these genes (Figure S7H).

As a test case to validate mechanisms of gene regulation in organoids compared to endogenous processes, we chose to investigate *FEZF2,* a TF that regulates the fate specification of a subset of layer 5 CFuPN (Chen, Schaevitz and McConnell, 2005; Molyneaux *et al*., 2005; Eckler *et al*., 2014). In the mouse, *Fezf2* is expressed in aRG at early stages of corticogenesis, and its expression becomes restricted to CFuPN as development progresses (Chen, Schaevitz and McConnell, 2005; Eckler *et al*., 2014). In our cortical organoid transcriptional atlas, at 3 months in culture, expression of *FEZF2* and its neighboring gene *CAPDS* is largely restricted to the CFuPN population (Figure S7I). We generated a map of predicted cis-regulatory interactions by calculating co-accessible sites across single cells using the Cicero package (Pliner *et al*., 2018). This showed that while there are accessible chromatin regions at or near the *FEZF2* locus in all cell types at 3 months, there is a high-scoring putative *cis*-regulatory interaction between *FEZF2* and an upstream region within *CADPS*, which is only accessible in CFuPN (Figure S7I-K). Indeed, previous studies in both mouse and human fetal tissue have identified this region as a cell type-specific enhancer with a high degree of conservation across vertebrates, which is exclusively active in CFuPN (Eckler *et al*., 2014; Markenscoff-Papadimitriou *et al*., 2020). Using the SHARE-seq data, we observed that the *FEZF2* DORC is accessible in aRG and oRG progenitor cells at 1 and 2 months, before the onset of expression of the gene (Figure S7L), in accordance with prior work showing that accessibility often precedes gene expression (Ma *et al*., 2020).

### Epigenetic signatures of metabolic pathways in organoid cells

To assess epigenetic changes in cells with high metabolic gene expression, we leveraged our SHARE-seq dataset. We first assigned each cell a module score for the MSigDB Hallmark

Hypoxia and Glycolysis gene sets using the scRNA-seq data, as above. Then, we used the corresponding scATAC-seq data to find regions that were differentially accessible between cells with high and low hypoxia module scores. This resulted in 3,189 DARs, out of 216,877 regions tested (1.5% of detected peaks, Table S13). We matched these regions to genes using the gene-peak associations we had previously calculated for the SHARE-seq cells. There were 320 genes regulated by significantly downregulated DARs in the high-hypoxia-scored cells, but these genes were not enriched for any GO terms. On the other hand, the 249 genes regulated by significantly upregulated DARs were enriched for metabolic GO processes including “response to hypoxia” (FDR=0.02, including *DDIT4*, *VEGFA*, and *HIF1A*), as well as for neurodevelopmental terms including “forebrain development” (FDR=0.05, including *LHX2*, *NR2F2*, and *SLC7A11*). Next, we looked for TF motifs whose accessibility correlated with metabolism module scores in individual timepoints and cell types. This identified TFs such as LHX2 and SOX4, associated with hypoxia scores in certain cell types at the 2- and 3-month timepoints (Table S14, Figure S18D-E).

Taken together, this data indicates that upregulation of metabolic genes can also be seen on an epigenetic level.

### Differential expression of metabolic gene sets in other organoid models

We sought to assess if differences in metabolic gene expression across different cell types were observed in organoids grown with different protocols. We therefore generated whole-brain organoids using an established protocol (Quadrato *et al*., 2017), and profiled them by scRNA-seq after 3.5 months *in vitro* (69,333 cells) (Figure S19A-B). We first extracted cells expressing forebrain markers, and then applied a classifier trained on our cortical organoid dataset to assign cell-type identities (Figure S19C-D). Finally, we assessed expression of metabolic gene signatures across both cortical and non-cortical cell types within these whole-brain organoids. In agreement with our results in the cortical organoid model, we observed an enrichment of glycolysis and hypoxia genes in aRG and in a population identified as unspecified PN, compared to the other cortical cell types in these organoids (Figure S19E, F, H). The MSigDB oxidative phosphorylation signature did not show significant enrichment in any particular cortical cell type in this organoid model, similar to the dorsal organoids (Figure S19G). This shows that multiple organoid models have similarly restricted differences in metabolic gene expression across similar cell types.

### Comparison of metabolic differences across organoids and endogenous fetal cells

To examine the similarities in metabolic gene expression between organoid and endogenous fetal cells, we first applied Compass, an algorithm that uses flux balance analysis to model the metabolic state of single cells from scRNA-seq data (Wagner *et al*., 2021), to model the cells using “potential activity scores” for each metabolic reaction. Using these scores, cells from organoids after 3 months *in vitro* clustered together with the two published endogenous scRNA-seq datasets after principal component analysis (Figure 4F and S18F). The first principal component (PC1) correlated strongly with cell complexity (Figure 4G and S18G), as previously reported (Wagner *et al*., 2021). None of the top 20 Principal Components (PCs) captured differences between data sets (Figure 4G), indicating that organoid and endogenous cells cluster together well based on their overall metabolic flux. Reactions involved in glycolysis, such as the phosphofructokinase pathway, were upregulated in organoids, while some pathways involved in cellular transport were downregulated in organoids (Figure S18H).

To determine the effects of individual metabolic pathways in more detail, we examined 38 MSigDB gene sets spanning a broad range of metabolic and RNA-processing processes, including metabolic pathways that have been described as important for neuronal progenitor proliferation and cortical development (Khacho, Harris and Slack, 2019; Namba *et al*., 2020). We combined all cells derived from our cortical organoids at 3 months *in vitro* with cells from human fetal cortex (Polioudakis *et al*., 2019; Trevino *et al*., 2021), and scored each gene set expression (Figure S20A). Most gene sets showed a similar distribution of module scores across datasets, including glycolysis- and oxidative phosphorylation-related gene sets. Importantly, the expression of many of these metabolic gene sets showed a similar degree of variability when comparing different fetal datasets. Notably, in most cases where expression differences in metabolic pathways between organoid and fetal cells were observed, all cells were similarly affected, rather than showing cell type-dependence (Figure S20B).

Finally, we leveraged the RRHO2 analysis to identify processes enriched among the genes up-regulated in organoids but downregulated in fetal cells. Gene lists from the upper left quadrant (i.e., genes up-regulated in organoids) of the RRHO2 results from corresponding cell types were submitted to Gene Ontology analysis (Figure S20C). Synapse- and cell cycle-related terms were the most prominent biological processes found to differ between organoid and fetal cells, possibly indicating differences in the maturation of the cells being compared. Among the 38 MSigDB metabolic gene sets, only a few pathways, including oxidative phosphorylation and lipid metabolism, ranked high in organoids and low in human datasets (Figure S20D).

### Creation of a molecular developmental trajectory tree using the SHARE-seq dataset

In order to have a complete picture of the molecular logic underlying human cortical projection neuron specification, we built a new branched trajectory tree leveraging the SHARE-seq map, containing the transcriptome and epigenome from the same individual organoid cells. The URD algorithm was applied to the RNA-seq data as before, in order to recapitulate the beginning stages of our full tree, and then the epigenetic data for these cells was projected onto the tree. Here, cells also progressed in pseudotime according to both age and differentiation state (Figure S25J-L). The two branching points obtained in this new tree closely resembled branch 1 and 2 of the previous one, and the gene programs associated with each segment in the RNA-only tree were enriched in the expected segments of this new one (Figure S25N). We then sought to identify TFs that act in each of the projection neuron lineages represented in this tree, i.e., CFuPN and CPN. For this, we split each segment of the tree into sections, calculated enriched DARs in cells from each section, and found TF motifs enriched in those DARs. This analysis identified known and novel TFs that may act at different steps of neuronal development (Figure S25M). Early stages of the cascade, mainly composed by the earliest aRG sampled, displayed enrichment for motifs for TFs known to act in neural progenitors of several regions of the neural tube (e.g., *LHX8* ventral telencephalon; *VSX1* retina), likely indicating a higher chromatin accessibility of aRG at this earlier timepoints and consistent with the presence of cell types from multiple brain regions (Figure 1A-B, Figure S2). In later segments in which the tree is still mostly composed of cortical progenitor cells, there was enrichment for motifs of the cortical progenitor TFs *EMX1*, and *EMX2*. These segments were also enriched for motifs for the TFs *LHX1*, *LHX9* and *PBX3*, known to be expressed by the earliest neurons populating the cortical plate (Moreau *et al*., 2021). At later stages related to neuronal specification, there was an enrichment of motifs for families of TFs previously implicated in neuronal development (e.g., *E2F*, *NFI*) as well as others with no previously-described role in projection neuron diversification (e.g., *TFAP2*, *ELK*, *NFY*).

## Methods

### Materials availability

Cell lines used in this study are available from their respective repositories (see Experimental Model & Subject Details), except for Mito210. There are restrictions to the availability of the Mito210 cell line due to Materials Transfer Agreement conditions, requests will be handled by the Lead Contact. This study did not generate new unique reagents.

### Data and code availability

Count and meta data from scRNA-seq and scATAC-seq are available on the Single Cell Portal, at https://singlecell.broadinstitute.org/single_cell/study/SCP1756. Raw data will be deposited into a restricted access database before publication. Code used during data analysis is available at https://github.com/AmandaKedaigle/OrganoidAtlas.

### Pluripotent stem cell culture

The female HUES66 embryonic stem cell (ESC) line (Chen *et al*., 2009) was provided by the Harvard Stem Cell Institute; the male GM08330 iPSC line (aka GM8330-8) was provided by M. Talkowski Lab (MGH) and was originally from the Coriell Institute; the male PGP1 (Personal Genome Project 1) human iPSC line (a.k.a GM23338) was from the laboratory of G. Church (Church, 2005); the male 11a human iPSC cell line was from the Harvard Stem Cell Institute; the male Mito210 human iPSC line was provided by B. Cohen Lab (McLean Hospital); and the male H1 parental hESC line (a.k.a. WA01) was purchased from WiCell. Three clones of the Mito210 and PGP1 human iPSC lines were used, and two clones of the H1 hESC line were used. The PGP1 line was authenticated using STR analysis completed by TRIPath (2018); the HUES66 line was authenticated using STR completed by GlobalStem Inc (2008). For authentication of the 11a cell line refer to (Quadrato *et al*., 2017). The H1 and GM08330 lines were authenticated using STR analysis completed by WiCell (2021). The Mito210 lines were authenticated by genotyping analysis (Fluidigm FPV5 chip) performed by the Broad Institute Genomics Platform (2017). The GM08330 line has an interstitial duplication in the long (q) arm of chromosome 20. All cell lines tested negative for mycoplasma contamination.

Cell lines were cultured in feeder-free conditions on 1% Geltrex (Gibco) coated cell culture dishes (Corning) using either mTESR1 medium (StemCell Technologies) or mTESR+ medium (StemCell Technologies) with 100 U/mL penicillin and 100µg/ml streptomycin, at 37°C in 5% CO2. PSC cultures and colonies were dissociated with the Gentle Cell Dissociation Reagent (StemCell Technologies) or ReLeSR (StemCell Technologies). All human PSC cultures were maintained below passage 50, were negative for mycoplasma, and karyotypically normal (analysis performed by WiCell Research Institute). All of the research conducted for this publication was done in accordance with relevant laws and policies and with ethical oversight from the Harvard University Cambridge-area Institutional Review Board and the Harvard Embryonic Stem Cell Research Oversight Committee.

### Dorsal organoids protocol

Dorsally patterned forebrain organoids were generated as previously described in (Velasco *et al*., 2019; Velasco, Paulsen and Arlotta, 2019). Briefly, on day 0, human iPSC or ESC, were dissociated to single cells with Accutase (Gibco), and 9,000 cells per well were reaggregated in ultra-low cell-adhesion 96-well plates with V-bottomed conical wells (sBio PrimeSurface plate; Sumitomo Bakelite) in Cortical Differentiation Medium (CDM) I, containing Glasgow-MEM (Gibco), 20% Knockout Serum Replacement (Gibco), 0.1 mM Minimum Essential Medium non-essential amino acids (MEM-NEAA) (Gibco), 1 mM pyruvate (Gibco), 0.1 mM 2-mercaptoethanol (Gibco), 100 U/mL penicillin, and 100 μg/mL streptomycin (Corning). For the H1 line, Mito210 c3 and PGP1 c3, cells were plated and formed in the same pluripotent medium in which they were maintained for 1 day to better enable embryoid body formation. From day 0-6, ROCK inhibitor Y-27632 (Millipore) was added (final concentration of 20 μM). From day 0-18, Wnt inhibitor IWR1 (Calbiochem) and TGFβ inhibitor SB431542 (Stem Cell Technologies) were added (final concentration of 3 μM and 5 μM, respectively). From day 18, the aggregates were cultured in ultra-low attachment culture dishes (Corning) under orbital agitation in CDM II, containing DMEM/F12 medium (Gibco), 2mM Glutamax (Gibco), 1% N2 (Gibco), 1% Chemically Defined Lipid Concentrate (Gibco), 0.25 μg/mL fungizone (Gibco), 100 U/mL penicillin, and 100 μg/mL streptomycin. On day 35, cell aggregates were transferred to spinner-flask bioreactors (Corning) and maintained in CDM III (CDM II supplemented with 10% fetal bovine serum (FBS) (GE-Healthcare), 5 μg/mL heparin (Sigma) and 1% Matrigel (Corning)). From day 70, organoids were cultured in CDM IV (CDM III supplemented with B27 supplement (Gibco) and 2% Matrigel).

### Whole brain organoid protocol

Unpatterned brain organoids were generated as previously described (Quadrato *et al*., 2017; Quadrato, Sherwood and Arlotta, 2017). Briefly, on day 0, human iPSC or ESC, were dissociated to single cells with Accutase (Gibco), and 2,500 cells were plated in each well of a 96-well plate and cultured in embryoid body medium as previously described (Lancaster and Knoblich, 2014). On day 6, embryoid bodies were transferred to low attachment, flat-bottom, 24-well plates in 500 μl of intermediate induction medium consisting of DMEM/F12, Knockout Serum Replacement (Gibco), 0.9% FBS GE-Healthcare), 0.7% N2 (Gibco), Glutamax (Gibco), MEM-NEAA (Gibco), and 0.7 μg/ml heparin. On day 8, 500 μl of neural induction medium was added to each well. On day 10, organoids were embedded in Matrigel (Corning), and transferred to cerebral differentiation medium (as described in (Lancaster and Knoblich, 2014)). On day 14, organoids were transferred to spinner flasks. Medium was changed weekly. After one month in culture, 14 ng/ml of BDNF (Preprotech, 450-02B) was added to the medium.

### Fetal human brain tissue

Human fetal tissue samples were obtained from the Allen Institute for Brain Science in respect of IRB guidelines from Harvard University. Samples processed for single-nuclei RNAseq were obtained from 4 donors (14, 15, 16, and 18 post-conception weeks (PCW)). All of the research conducted for this publication was done in accordance with relevant laws and policies and with ethical oversight from the Harvard University Cambridge-area Institutional Review Board and the Harvard Embryonic Stem Cell Research Oversight Committee.

### Immunohistochemistry

Samples were fixed in 4% paraformaldehyde (PFA) (Electron Microscopy Services). Samples were washed with 1X phosphate buffered saline (PBS) (Gibco), cryoprotected in a 30% sucrose solution, embedded in optimum cutting temperature (OCT) compound (Tissue Tek) and cryosectioned at 12-18 µm thickness. Sections were washed with 0.1% Tween-20 (Sigma) in PBS, blocked for 1 hour at room temperature (RT) with 6% donkey serum (Sigma) + 0.3% Triton X-100 (Sigma) in PBS and incubated with primary antibodies overnight diluted with 2.5% donkey serum + 0.1% Triton X-100 in PBS. After washing, sections were incubated at room temperature with secondary antibodies diluted in the same solution as with primary antibodies (1:1000-1:1200; Key Resources Table) for 2 hours at room temperature, washed, and incubated with DAPI staining (1:10,000 in PBS + 0.1% Tween-20) for 15 minutes to visualize cell nuclei. Primary antibodies were diluted as specified in the Key Resources Table. The protocol was adapted for immunohistochemistry for SLC16A3, GORASP2, VIMENTIN, HOPX and OLIG2. Briefly, the first wash was followed by incubation of the sections in boiling temperature sodium citrate buffer pH6 (10 mM sodium citrate, 0.05% Tween-20) or 1x IHC Select Citrate Buffer pH 6.0 (Sigma, 21545) for 40 minutes. Then the sections were extensively washed with 0.1% Tween-20 and the protocol was followed as described above.

### Hypoxyprobe

The Hypoxyprobe (100 mg pimonidazole HCl plus 1.0 mL of 4.3.11.3 mouse Mab, Hypoxyprobe kit, HP1-100kit) was added to the medium at a concentration of 100 µM and organoids were incubated with the probe for 1.5h at 37C and 5% CO_2_. The organoids were then extensively washed with PBS and fixed in 4% PFA (Electron Microscopy Services) overnight. Immunohistochemistry was performed as described in the above section, including the antigen retrieval step. The primary antibody used at a 1:50 dilution was included in the kit.

### Microscopy

Immunofluorescence images were acquired with a Zeiss Axio Imager.Z2 and Lionheart™ FX Automated Microscope (BioTek Instruments). Images were analyzed and processed with the Gen5 (BioTek Instruments) and Zen Blue (Zeiss) image processing software, and ImageJ (Schneider, Rasband and Eliceiri, 2012).

### Dissociation of brain organoids and scRNA-seq

Individual brain organoids were dissociated into a single-cell suspension using the Worthington Papain Dissociation System kit (Worthington Biochemical). A detailed description of the dissociation protocol is available at Protocol Exchange, with adaptations depending on age and size (Quadrato, Sherwood and Arlotta, 2017; Velasco, Paulsen and Arlotta, 2019). We resuspended dissociated cells in ice-cold PBS containing 0.04% BSA (Sigma, PN-B8667), counted them with Countess II (Thermo Fisher Scientific), and then adjusted the volume to the final concentration of 1,000 cells/μl. We loaded cells onto a Chromium™ Single Cell 3’ Chip (10x Genomics, PN-120236), and processed them through the Chromium Controller to generate single cell GEMs (Gel Beads in Emulsion). scRNA-seq libraries were prepared with the Chromium™ Single Cell 3’ Library and Gel Bead Kit v3 and v3.1 (10x Genomics, PN-1000075, 1000121), with the exception of a few libraries in the earlier experiments that were prepared with a v2 kit (10x Genomics, PN-120236). See Table S17 for information on how many cells were estimated to be loaded and what kit was used. We pooled libraries from different samples based on molar concentrations and sequenced them on a NextSeq 500 or NovaSeq 6000 instrument (Illumina) with 28 bases for read 1 (26 bases for v2 libraries), 55 bases for read 2 (57 bases for v2 libraries), and 8 bases for Index 1. If necessary, after the first round of sequencing, we re-pooled libraries based on the actual number of cells in each and re-sequenced with the goal of producing an equal number of reads per cell for each sample.

### Single cell RNA-seq data pre-processing

All cortical brain organoid scRNA-seq datasets consisted of three individual organoids, except for the PGP1 c2 batch 17 at 6 months in which, as previously published (Velasco *et al*., 2019), one organoid showed a differentiation defect and was excluded. Whole-brain organoid single-cell datasets consisted of four (HUES66) and four (GM8330) individual organoids. ScRNA-seq reads were aligned to the GRCh38 human reference genome and the cell-by-gene count matrices were produced with the Cell Ranger pipeline (10x Genomics, (Satpathy *et al*., 2019), for version number, see Table S17). Default parameters were used, except for the ‘–cells’ argument. Data was analyzed using the Seurat R package v3.2.2 (Stuart *et al*., 2019) using R v3.6. Cells expressing a minimum of 500 genes were kept, and UMI counts were normalized for each cell by the total expression, multiplied by 106, and log-transformed.

In the case of GM08330 1-month organoids, cells were demultiplexed using genotype clustering from cells from a different experiment that were sequenced in the same lane. To demultiplex, SNPs were called from Cell Ranger BAM files with the cellSNP tool v0.1.5, and then the vireo function was used with default parameters and n_donor=2, from the cardelino R library v0.4.0 (Huang, McCarthy and Stegle, 2019) to assign cells to each genotype.

### Single cell RNA-seq dimensionality reduction and clustering

For each dataset of organoids, variable genes were found using the “mean.var.plot” method with default parameters, and the ScaleData function was used to regress out variation due to differences in total UMIs per cell and to cell cycle gene expression, following Seurat’s CellCycleScoring method. Principal component analysis (PCA) was performed on the scaled data for the variable genes, and the top 30 PCs were used for downstream analysis. Cells were clustered in PCA space using Seurat’s FindNeighbors on the top 30 PCs, followed by FindClusters with resolution=1.0, except for the Mito210 c1 23 days data set, in which FindClusters with resolution=2.0 was used. Cells were visualized by Uniform Manifold Approximation and Projection (UMAP) on the top 30 PCs.

### Single cell RNA-seq cluster annotations

For each dataset, upregulated genes in each cluster were identified using the VeniceMarker tool from the Signac package v0.0.7 from BioTuring (https://github.com/bioturing/signac). Cell types were assigned to each cluster by looking at the upregulated genes with lowest Bonferroni adjusted p values. In some cases, clusters were further subclustered to assign identities at higher resolution (See https://github.com/AmandaKedaigle/OrganoidAtlas/blob/master/Subclustering.R for specific clusters that were separated; for cluster numbers, refer to the metadata available for download from the Single Cell Portal).

Adjusted mutual information (Xuan Vinh, Epps and Bailey, 2010) was calculated between cell type assignments and individual organoids with the aricode R package v1.0.0 (Chiquet, Rigaill and Sundqvist, 2020).

### Single cell RNA-seq data integration

For visualization purposes and downstream analyses within each of the timepoints (1, 1.5, 2, 3-3.5, 4, 5 and 5.5-6 months), cortical organoid datasets were merged using Seurat, re-normalized and scaled as above, batch-corrected using Harmony v1.0 with default parameters (Korsunsky *et al*., 2019), and visualized using UMAP on the first 30 Harmony dimensions.

### Single cell ATAC-seq

Prior to processing, organoids were frozen in Recovery cell culture freezing medium (Thermo Fisher Scientific) at −80°C. Nuclei were extracted from 1-, 3-, and 6-month organoids derived from the Mito210 c1 line using two types of procedures, due to their size differences. For the 1-month organoids, nuclei were extracted following a 10x Genomics-provided protocol (Genomics, 2019) to minimize material loss. For the 3- and 6-month organoids, we used a sucrose-based nuclei isolation protocol (Corces *et al*., 2017) to better remove debris. ScATAC-seq libraries were prepared with the Chromium™ Single Cell ATAC Library & Gel Bead Kit (10x Genomics, PN-1000110). Approximately 15,300 nuclei were loaded in each channel (to give an estimated recovery of 10,000 nuclei per channel). Libraries were pooled from different samples based on molar concentrations and sequenced with 1% PhiX spike-in on a NextSeq 500 instrument (Illumina) with 33 bases each for read 1 and read 2, 8 bases for Index 1, and 16 bases for Index 2. See Table S17 for sequencing information.

### Single cell ATAC-seq data pre-processing

Reads from scATAC-seq libraries were aligned to the GRCh38 human reference genome and the cell-by-peak count matrices were produced with the Cell Ranger ATAC pipeline v1.2.0 (10x Genomics). Default parameters were used, except with ‘–force-cells 5000’ in one organoid (ATAC-Org5) at 3 months. Data were analyzed using the Signac R package v1.1.0 (Stuart *et al*., 2021) using R v3.6. Annotations from the EnsDb.Hsapiens.v86 genome annotation package (Rainer, 2017) were added to the object. After consideration of the QC metrics recommended as part of Signac, cells were retained that had 1,500-20,000 fragments in peak regions, at least 40% of reads in peaks, less than 5% of reads in blacklisted regions, a nucleosome signal of less than 4, and a TSS Enrichment score greater than 2. Latent semantic indexing (LSI) was performed to reduce the dimensionality of the data (counts were normalized using term frequency inverse document frequency, all features were set as top features, and singular value decomposition (SVD) was performed). The top LSI component was discarded as it correlated strongly with sequencing depth, and components 2-30 were used for downstream analysis.

To overlap accessible regions with annotated regions of the genome, peaks called in each scATAC-seq timepoint were overlapped with promoters and gene bodies (defined using the method in Signac’s GeneActivity function) and with human enhancer regions from Genehancer v4.4 (Fishilevich *et al*., 2017), using the “overlapsAny’’ function from the GenomicRanges R package.

### Single cell ATAC-seq clustering and cluster annotation

Cells were clustered using Seurat’s FindNeighbors on the top 30 PCs, followed by FindClusters with the smart local moving (SLM) algorithm and otherwise default parameters. Variation in the cells was visualized by UMAP.

Cells were annotated by two parallel methods. In one method, scATAC-seq data were integrated with scRNA-seq data from the corresponding Mito210 c1 dataset for each timepoint, using Seurat’s TransferData to predict cell type labels for the ATAC cells. Separately, differentially accessible regions (DARs) were called per scATAC-seq cluster using FindMarkers with the logistic regression framework, with the number of fragments in peak regions as a latent variable. These DARs were mapped to the closest genes. Top genes per cluster were used to confirm and refine cluster cell type assignments from those based on transferring labels from scRNA-seq.

Adjusted mutual information (Xuan Vinh, Epps and Bailey, 2010) was calculated between cell type assignments and individual organoids with the aricode R package v1.0.0 (Chiquet, Rigaill and Sundqvist, 2020).

### Identification of co-accessible regions from single cell ATAC-seq

To identify co-accessible sites across each scATAC-seq dataset, the Cicero package was employed (Pliner *et al*., 2018). A Cicero CellDataSet object was created from the Signac object and run_cicero was run with sample_num=100.

### Single cell ATAC-seq motif enrichment

Motif enrichment analysis was performed by first getting a set of differentially accessible peaks per cell type, using FindMarkers as above. For each cell type, peaks with increased accessibility with an adjusted p value (Bonferroni correction) less than 0.1 were then supplied to the HOMER software v4.11.1 (Heinz *et al*., 2010), using a 300 bp fragment size and masking repeats. The top 5 de novo motifs per cell type found by HOMER, with a p value <= 1e-10, are reported in Table S7, along with all TFs whose known binding sites match that motif with a score >= 0.59.

### SHARE-seq and bulk ATAC-seq

Nuclei were isolated from fresh-frozen organoids in HEPES Lysis Buffer (10 mM HEPES pH 7.3, 10 mM NaCl, 3 mM MgCl2, 0.1% NP-40) for 5 min on ice. Organoids from day 90 were dissociated using spring scissors (Roboz, RS-5650), while all others were dissociated by gentle pipetting. Samples were then diluted in 1 mL HDT Buffer (10 mM HEPES pH 7.3, 10 mM NaCl, 3 mM MgCl2, 0.1% Tween-20, 0.01% digitonin). Nuclei were pelleted at 800xg for 3 min at 4 C and resuspended in room-temperature HDT Buffer at a concentration of 1 million nuclei/ml. Formaldehyde was added to a 0.2% final concentration and nuclei were fixed for 5 min at RT. Fixation was quenched with glycine (140 mM final concentration), Tris pH 8.0 (50 mM final concentration), and BSA (0.1% final concentration) for 5 min on ice. Nuclei were washed two additional times with HDT Buffer, split into single use aliquots, pelleted and stored at −80C. All buffers were supplemented with Enzymatics (Qiagen, Y9240L) and SUPERase (Thermo Fisher Scientific, AM2694) RNase inhibitors.

SHARE-seq was performed as described in (Ma et al. 2020) with the following modifications: NIDT Buffer (10 mM Tris pH 7.5, 10 mM NaCl, 3 mM MgCl2, 0.1% Tween-20, 0.01% digitonin) was used instead of NIB. For reverse transcription and template switching, 5X Smart-seq3 Buffer (40 mM DTT, 125 mM Tris pH 8.0, 5 mM GTP, 150 mM NaCl, 12.5 mM MgCl2) was used instead of 5X Maxima RT Buffer.

Bulk ATAC seq was performed on day 23 organoids (4 separate organoids, Mito210 cell line, clone 2, batch 20) by isolating nuclei as described above from four fresh-frozen organoids followed by transposition reaction (without fixation) using the same reaction conditions as described above. Following transposition, DNA was purified using NucleoSpin Gel and PCR Clean-up kits (Takara, 740609.250). Libraries were prepared identically to SHARE-seq ATAC libraries.

Both scRNA-seq and scATAC-seq raw sequencing reads were processed as described in (Ma *et al*., 2020) (pipeline available at https://github.com/masai1116/SHARE-seq-alignment).

### SHARE-seq scRNA-seq dimensionality reduction & clustering

Cell barcodes with a minimum of 1,000 UMIs were retained. Two rounds of filtering were performed to computationally remove doublets and aggregated cells. First, the UMI count matrix was normalized to the average number of UMIs detected across all cells. The 5,000 genes with the highest variance were log2+1 transformed and PCA was performed on this set of genes. The top 20 components were used to find the 20 k-nearest neighbors (KNN) and the estimated library size for each cell barcode smoothened across KNN. Cell barcodes with disproportionately high smoothened library size (identified as outlier from the distribution of library sizes per experiment) were identified as “clumps” and computationally removed. Next, UMI counts for the remaining cell barcodes were processed using Scrublet (Wolock, Lopez and Klein, 2019) and the top scoring barcodes were identified as “doublets” and removed. The top 20 principal components were used for UMAP visualization, identifying KNN and Louvain clustering.

### SHARE-seq scATAC-seq dimensionality reduction and clustering

Fragment files from all single cell sequencing runs were combined and peaks were called using MACS2 (Zhang *et al*., 2008). For bulk ATAC on day 23 samples, data was processed as described in (Buenrostro *et al*., 2015). The peak sets were combined and filtered as described in (Lareau *et al*., 2019). Briefly, +/- 400 bp windows were extended from each peak summit and sorted according to MACS2 significance scores. Starting with the most significant peak window, all other overlapping windows with lower significance were removed. This was repeated until no overlapping windows were left. The resulting peaks were resized to 300 bp and these intervals were used for subsequent analysis.

Reads in peaks were counted using Signac (Stuart *et al*., 2021) and cell barcodes were filtered for a minimum fraction of reads in peaks of 0.2, and estimated library size of at least 2000 reads. Cell “clumps” were identified similarly to what described for scRNA-seq; topics were identified using cisTopic (Bravo González-Blas *et al*., 2019) and the top 20 topics were used for finding KNN. Library sizes were smoothened over these KNN and the top barcodes were removed. Cell barcodes identified as “doublets’’ in the scRNA-seq analysis above were also removed. Topics were identified on the remaining cells and the top 20 were used for UMAP visualization, KNN identification and Louvain clustering.

### SHARE-seq motif enrichment

Motif scores were computed on all scATAC cell barcodes, filtered as described above, using chromVAR (Schep *et al*., 2017). For subsequent analysis, data was then subsetted to cell barcodes shared between the RNA and ATAC datasets. For each TF, the RNA-motif correlation was calculated as the Spearman correlation coefficient across all cells between the TF’s motif scores and the same TF’s gene expression values.

### SHARE-seq gene-peak associations and DORC scores

Gene-peak associations were found on the set of all filtered cells shared between the scATAC-seq and scRNA-seq datasets using the method described in (Ma *et al*., 2020). Briefly, for each gene, peaks within +/- 50 kb of their TSS were considered for possible associations (median 9 peaks per gene). Spearman correlations were calculated between normalized scATAC counts and normalized scRNA UMI counts. A set of 50 background peaks per tested region was selected using chromVAR, controlling for GC content and accessibility across all cells. The mean and standard deviation of this background set was used to calculate a Z-score and p-value for the observed association. Significant associations were called as those with p-value less than 0.05.

Domains of regulatory chromatin (DORCs) were called as genes with greater than 5 associated peaks and DORC scores were the total number of reads within these associated peaks. For visualization, DORC scores were smoothened across the KNN identified in the total scATAC dataset and normalized RNA expression values were smoothened across KNN identified in the total scRNA-seq dataset. After subsetting to cells shared in both datasets, both DORC scores and RNA expression values were quantile normalized prior to visualization. The residual was defined as the (normalized smoothened DORC score) – (normalized smoothened RNA expression value).

### Slide-seqV2

For Slide-seqV2, organoids were embedded in OCT (Tissue Tek) and cryosectioned at 10 µm thickness. Sections were transferred to a custom-made array of densely packed barcoded beads (termed ‘pucks’) for Slide-seqV2 experiments. Library construction was performed as described (Stickels *et al*., 2021). Briefly, first-strand synthesis was performed by incubating the puck with tissue sections in the reverse-transcription solution followed by tissue digestion, second-strand synthesis, library amplification, and purification. The Slide-seqV2 libraries were sequenced with a NextSeq500 with an estimated ∼100 million reads per puck. See Table S17 for sequencing information.

### Slide-seqV2 data pre-processing

Sequencing reads were aligned to the GRCh38 human reference genome and followed by the Slide-seq pipeline (https://github.com/MacoskoLab/slideseq-tools) and as previously described (Stickels *et al*., 2021). Since organoid sizes were much smaller than the Slide-seq pucks, images were manually cropped to the edges of the organoids and beads outside of the images were excluded.

### Slide-seqV2 cell type decomposition

Cell type decomposition was performed by RCTD (Cable *et al*., 2021). Briefly, cell type profiles learned from scRNA-seq data corresponding to the same timepoints and cell line were used to decompose mixtures from the Slide-seqV2 data. First, for each timepoint, a reference was made by using the age-matched scRNA-seq dataset. Next, the RTCD package fits a statistical model to estimate the mixture and cell type identities at each bead of the Slide-seqV2 data. We restricted our analysis to beads with more than 200 UMIs. Finally, gene set expression of the top 50 genes from each cell type signature, as previously calculated from scRNA-seq data (Tables S1-S3), was visualized by plotting 100x the sum of the reads for all genes in the gene set that appeared in the Slide-seqV2 data, normalized by the number of UMI for each bead.

### Cell type signature analysis

To calculate lists of genes defining cell subtype identity from scRNA-seq data, we partitioned each dataset into a group containing cortical neurons and progenitors separately: for 1 month organoids, neurons (newborn DL PN and immature DL PN), and progenitors (aRG and IP); for 2 months organoids, excitatory neurons (i.e., CFuPN, CPN, and ‘unspecified PN’; Newborn CFuPN and CFuPN were merged into one cell type for this analysis), and progenitors (i.e. aRG, IP and oRG); for 3 months organoids, excitatory neurons (i.e., CFuPN, CPN, and ‘unspecified PN’) and progenitors (i.e., aRG, IP, and oRG); for 4 months organoids, excitatory neurons (i.e., CFuPN, CPN, and ‘unspecified PN’) and progenitors (i.e., aRG, IP, and oRG); for 5 months organoids, neurons (CPN, ‘unspecified PN’, and immature IN) and progenitors-glia (aRG, IP, oRG & Astroglia, oligodendrocyte precursors, and IN progenitors); for 6 months organoids, neurons (CPN, ‘unspecified PN’, and immature IN) and progenitors-glia (aRG, IP, oRG & Astroglia, oligodendrocyte precursors, and IN progenitors), the very few oRG II cells were excluded from 6 months analysis. For the purpose of plotting signatures of all cell types present in the organoid in Slide-seq data, the one-month signatures were recalculated including Cajal Retzius, Cortical hem and subcortical cells (Table S1). To examine processes enriched among genes up-regulated in organoids in the RRHO2 analysis, the neuron signatures from (Trevino *et al*., 2021) were re-calculated, combining all GluN cell types.

Next, in order to control for variation in the datasets, final signatures were calculated using a paired analysis, pairing cell type clusters occurring in the same individual organoids. Reads were summed across cells in each cell type in each organoid in all cell lines. Genes with less than 10 total UMI were excluded, and then DESeq2 (Love, Huber and Anders, 2014) was used to calculate DEGs between each cell type and all other cell types in its group. The DESeq2 design formula was “∼ organoid + CellType”, so that DESeq2 found differentially expressed genes between cell types (defined as the cell type of interest vs Other for each analysis), controlling for the effect of the individual organoid.

To examine whether expression of signature genes was consistent across individual organoids, module scores for the top 200 upregulated genes in each signature were calculated using Seurat’s AddModuleScore function on the same Seurat objects used as input to the DESeq2 method. For each cell type signature, datasets were split into the cell type of interest and remaining cells, and the average module score across each individual organoid was calculated. These averages were then submitted to a one-way ANOVA using the base R function aov, using the formula AverageValue ∼ organoid. None of these tests were significant for an overall difference of means (significance reported as Pr(>F) > 0.05).

### Analysis of previously published human fetal data

The dataset from (Polioudakis *et al*., 2019) was used, which contains Drop-seq data for 33,986 cells from human fetal cortex samples obtained at gestational week 17 and 18, and that had been aligned to the Ensembl release 87 Homo sapiens genome. Raw count data and associated metadata were downloaded from the CoDEx viewer (http://solo.bmap.ucla.edu/shiny/webapp/). Count data from all cells were loaded into a Seurat object and normalized using the same methods as described above for organoid data. To integrate with organoid data, objects were merged in Seurat, and batch-corrected using Harmony with default parameters.

The dataset from (Trevino *et al*., 2021) was also used. This dataset contains 57,868 single-cell transcriptomes and 31,304 single-cell epigenomes (ATAC-seq) from human fetal cortex samples from gestational week 16, 20, 21 and 24, generated using the 10x Genomics Chromium platform and aligned to the Cell Ranger GRCh38 genome. ScRNA-seq count data and metadata were downloaded from the authors’ GitHub (https://github.com/GreenleafLab/brainchromatin/blob/main/links.txt). Count data was normalized and integrated with organoid data as above. ScATAC-seq counts data and metadata were downloaded from GEO (GSE162170), and fragments files were downloaded from the authors’ GitHub. Cell types that would not be expected to be present in organoids (endothelial cells, pericytes, and microglia) were removed. MACS2 was used to rerun peak calling on the remaining cells via Signac’s CallPeaks function, grouped by cell type, yielding 326,515 peaks. A shared peak space was generated for each organoid timepoint using the reduce function from the GenomicRanges R package (Lawrence *et al*., 2013) to merge peaks called in the two datasets. This yielded 379,920 shared peaks with 1-month organoids, 351,207 shared peaks with 3-month organoids, and 358,664 shared peaks with 6-month organoids. Integration of fetal and organoid scATAC-seq data was performed using Signac.

### Fetal human brain tissue dissection and single nucleus RNA seq

Cryostat coronal sections (70-100 microns) were produced from human fetal somatosensory cortices and deposited on non-chargeable slides. The cortical plate (CP) was further dissected using single-use micro stab knifes (F.S.T. #72-2201). Following dissection, CP sections were gathered with a p1000 pipette in cold EZ-lysis buffer (Sigma, #NUC-101). For snRNA-seq, nuclei were isolated following the (Habib *et al*., 2017) protocol. In brief, tissue was transferred to a chilled dounce tube containing 2ml of ice-cold EZ-lysis buffer. Gently, tissue was homogenized with 20 strokes of pestle A followed by pestle B. The suspension was transferred to a chilled conical tube and incubated on ice for 5 minutes. Tissue suspension was then centrifuged at 500xg, 4 °C for 5 min and resuspended in 1ml of Nuclei Suspension Buffer (NSB): RNAse-free molecular biology grade PBS, 0.1%BSA (NEB #B9000S), 0.2U/ml Recombinant ribonuclease inhibitor (Takara, #2313B). A second centrifugation (same conditions) was performed before final resuspension in 500 ml chilled NSB. Nuclei yield was quantified by Trypan blue staining using disposable Hemacytometer chambers (NCYTO C-Chip™, SKC - DHCN015) prior to Chromium Single Cell 10x V3 assay.

### Fetal human brain tissue single nucleus RNA-seq pre-processing

SnRNA-seq reads were aligned to the GRCh38 human reference genome and the cell-by-gene count matrices were produced with the Cell Ranger pipeline version 3.0.2 using default parameters. Data was analyzed using the Seurat R package v3.2.0 using R v3.6.0. Cells expressing a minimum of 500 genes were kept, and UMI counts were normalized and scaled using the SCTransfrom function from Seurat (with vars.to.regress parameter set to the number of genes, number of UMIs, and percent mitochondrial gene expression). This function internally computes variable genes and scales the data on these genes.

### Fetal brain tissue single nucleus RNA-seq dimensionality reduction and clustering

PCA was performed on the scaled data of the ‘SCT’ assay created during the SCTransform step. Cells were clustered in PCA space using Seurat’s FindNeighbors on the top 30 PCs, followed by FindClusters with resolution=0.3. Cells were visualized by UMAP on the top 30 PCs. Upregulated genes in each cluster were identified using the VeniceMarker tool from the Signac package v0.0.7 from BioTuring (https://github.com/bioturing/signac). Cell types were assigned to each cluster by looking at the upregulated genes with lowest Bonferroni adjusted p values. In some cases, clusters were further sub-clustered to assign identities at higher resolution (See https://github.com/AmandaKedaigle/OrganoidAtlas/blob/master/Subclustering.R for specific clusters that were split; for cluster numbers, refer to the metadata available for download from the Single Cell Portal).

### Matching of human organoid and fetal cells

To assign organoid cells to fetal cell categories, as defined in (Polioudakis *et al*., 2019; Trevino *et al*., 2021), we used the fetal categories to train a random forest (RF) classifier to distinguish between cell types. This was done with the tuneRF function in the randomForest R package (Liaw, A and Wiener, 2002), with doBest = T, after removing cell types not found in organoids (microglia, endothelial cells, and pericytes), and downsampling the data to have an equal number of cells per cell type. Variable genes from each fetal cell dataset were used to train the model. For validation, 20% of the cells were reserved as a cross-validation set, and compared to a null model prepared by shuffling the cell type labels on the training data set. The classifier was then applied to the organoid cell profiles to assign organoid cells to one of the human cortex cell types. Organoid cells at 1 and 1.5 months were subsetted to exclude cells which are not expected to have corresponding cells in the fetal data (Cajal Retzius cells, cortical hem, and subcortical cells).

To classify fetal cells to organoid cell categories, we trained a RF classifier using organoid data and applied it to classify human fetal cells. The data were then downsampled to have an equal number of cells per cell type, and the same cell types were excluded at 1 month. Variable genes from the organoid dataset were used to train the model. For validation, RF models trained on organoid data were tested on a cross validation held-out set of organoid cells (20% of total dataset), compared to null models.

### Matching of accessible chromatin in human organoids and fetal cortex

Shared peaks between fetal (Trevino *et al*., 2021) scATAC-seq fragments and fragments from each organoid timepoint (1, 3, and 6 months) were derived using the reduce function from the GenomicRanges R package (see above). Fetal scATAC-seq counts matrices were recalculated using the shared feature spaces. LSI and SVD were performed on processed fetal scATAC-seq counts matrices, and SVD components 2-30 were used to calculate UMAP embeddings. Seurat’s FindTransferAnchors function (with parameters k.anchor=7, max.features=250, and k.filter=NA) and MapQuery functions were used to transfer fetal cell type labels onto organoid cells, and to project organoid scATAC-seq data into the fetal UMAP space. Cajal Retzius, Cortical hem, and subcortical cells, which are not expected to have corresponding cells in the fetal dataset, were excluded from the 1-month organoid.

Fetal cells were down-sampled to a maximum of 450 cells per cluster, and DARs were calculated for each fetal cell type using Seurat’s FindMarkers with a logistic regression framework, with the number of fragments in peak regions as a latent variable. Organoid DARs for each cell type were recalculated at each organoid timepoint (1, 3, and 6 months) using the new shared peaks calculated for organoid and fetal cells. DAR signatures were ranked by degree of differentiation [-log10(p value) * sign(effect)] and then compared using the improved rank-rank hypergeometric overlap test as described above. Cajal Retzius, Cortical hem, and subcortical cells were once again excluded from the 1-month organoid.

### Comparison of cell type signatures between human fetal cells and cortical brain organoids

Organoid DEG signatures (above) were compared to fetal DEG signatures, calculated using the same DESeq2-based method as for organoids. Cells originally labeled as cycling progenitors in the human datasets were reassigned to reflect their cell types (oRG and IP, in (Polioudakis *et al*., 2019), Early_RG, nIPC, mGPC and IN progenitors in (Trevino *et al*., 2021)) rather than proliferative state, by using FindMarkers with default parameters to examine upregulated genes in clusters containing these cells. In the human fetal dataset presented here, progenitor cells were directly classified based on their identity rather than cell cycle state. DEG signatures were ranked by degree of differentiation [-log10(p value) * sign(effect)] and then compared using the improved rank-rank hypergeometric overlap test [RRHO2 R package v1.0; (Plaisier *et al*., 2010; Cahill *et al*., 2018)].

To identify processes enriched in genes that differ between signatures, overlaps between the discordant genes as identified by the RRHO2 test and published gene sets were evaluated. The metabolic genes lists used in the section below, “Impact of metabolic genes on cell classification”, as well as 42 gene sets relevant to neurobiology (see Fig. S20C) were downloaded from the Molecular Signatures Database (MSigDB; (Subramanian *et al*., 2005; Liberzon *et al*., 2011), gsea-msigdb.org), and were compared to the discordant genes. The statistical significance of gene overlap between two gene lists was calculated by hypergeometric test, and p values were adjusted using the Bonferroni correction.

### Weighted Gene Correlation Network Analysis

In order to learn patterns of coordinated gene regulation across the cortical organoid scRNA-seq datasets, WGCNA (Langfelder and Horvath, 2008) was applied to each dataset with multiple batches: 1, 1.5, 2, 3, 4, 5 and 6 months. The Seurat objects were downsampled to have an equal number of cells per organoid prior to applying WGCNA. Normalized gene expression data were further filtered to remove outlying genes, mitochondrial, and ribosomal genes. Outliers were identified by setting the upper (>9) and lower (<0.15) thresholds to the average normalized expression per gene. After processing, blockwiseModules function from the WGCNA v1.69 library was performed in R with the parameters networkType=”signed”, minModuleSize=4, corType=“Bicor”, maxPOutliers=0.1, deepSplit=3,trapErrors=T, and randomSeed=59069. Other than power, the remaining parameters were left as the default setting. The power parameter that determines the resolution of gene module output was chosen separately for each dataset. To pick an adequate parameter, we used the pickSoftThreshold function from WGCNA to test values from 1 to 30. Final resolution was determined by studying the output modules and choosing the resolution that captured the most variation in the fewest total number of modules - this resulted in a power of 8 for the 1 month harmonized dataset, 9 for the 1.5 months harmonized dataset, 14 for the 2 months harmonized dataset, 4 for the 3 months harmonized dataset, 6 for the 4 months harmonized dataset, 9 for the 5 months harmonized dataset, and 5 for the 6 months harmonized dataset.

### Module scores

Expression of gene sets identified in MSigDB (Subramanian *et al*., 2005; Liberzon *et al*., 2011) compared to random gene lists with similar average expression, was evaluated using Seurat’s AddModuleScore function using the downsampled objects used in WGCNA. The distribution of module scores between cell types was visualized using Seurat’s VlnPlot function. To assess the differences between module scores across cell types at each timepoint, the average of module scores was calculated in each cell type per individual organoid. These averages were then submitted to a one-way ANOVA using the base R function aov, with the formula AverageValue ∼ CellType. Since all tests were significant for an overall difference of means (significance Pr(>F) <0.05), post-hoc comparisons were done with the TukeyHSD function. Results from pairwise comparisons were visualized with a compact letter display using the multcompLetters function from the multcompView R package v0.1-8 (Graves, Piepho and Selzer, 2019) with default parameters.

For module score analysis of MSigDB metabolic pathways in human fetal cortical cells (Polioudakis *et al*., 2019), cell types that do not appear in the organoids were excluded from the analysis: microglia (48 cells), pericytes (117), and endothelial cells (237). For module score expression of MSigDB metabolic pathways in 3 months organoids cells, Cajal Retzius cells were excluded from the analysis (99 cells). Remaining cells from each dataset were grouped into broad cell type categories: Cells called “CPN”, “GluN1”, “GluN2”, “GluN3”, “GluN4”, “GluN5”, “GluN6”, “GluN7”, “GluN8”, “ExN”, “ExM”, and “ExM-U” were all labeled “UL Neurons.” Cells called “CFuPN”, “SP”, “ExDp”, “ExDp1”, and “ExDp2” were all labeled “DL Neurons.” Cells called “IP” and “nIPC” were labeled “IPs.” Cells called “CGE_IN”, “MGE_IN”, “Immature IN”, “InMGE”, and “InCGE” were labeled “INs.” Cells called “tRG”, “Early_RG”, “Late_RG”, “Cyc_Prog”, “aRG”, “oRG”, “PgS”, “PgG2M”, “vRG” were labeled “RG.” Unspecified PN from the organoids retained their unique label.

### Evaluation of metabolic activity with Compass

Activity scores for metabolic reactions in the RECON2 database (Swainston *et al*., 2016) were assigned to each cell from the organoid 3-month dataset, and two human fetal datasets (Polioudakis *et al*., 2019; Trevino *et al*., 2021), using the Compass tool (Wagner *et al*., 2021).

To calculate the PCA, datasets were each down-sampled to 1000 cells, and gene counts were normalized to Counts Per Million. This matrix was used as an input to Compass using default parameters and the homo_sapiens species. Downstream analysis was done as in the original Compass paper (Wagner *et al*., 2021). Briefly, the output penalties were set to zero if they were below 0.0004, and reactions which had a total score of 0 across all cells or that showed a total range across all cells within 0.001 were eliminated from the analysis. Hierarchical clustering was then performed on the penalty matrix, based on Spearman correlation, and reactions were clustered into metareactions based on cutting the hierarchical clustering tree at a Spearman correlation of 0.95. Metareactions were given penalty scores as the mean of their component reactions. The metareaction penalty matrix was then transformed by log1p. PCA was performed on this transformed data. The top 20 principal components were correlated with the number of UMI per cell, the cell type labels of each cell, and the dataset each cell derived from, using linear regression, with the lm R function. The R squared value from the resulting function was reported.

For differential activity analysis, cells from 3-month organoids and the (Trevino *et al*., 2021) fetal dataset were grouped into broad cell type categories as in the “Module Scores” section above. Datasets were then downsampled to have 50 cells from each broad cell type category. Only genes involved in one or more metabolic reactions (found using Compass’ –list-genes function) were retained, and then gene counts were normalized to Counts Per Million. Normalizing only metabolic-related genes should remove variability due to overall up- or down-regulation of metabolism across the board, emphasizing variable use of specific pathways. Counts were then submitted to Compass as above, and resulting penalty scores were processed as above, without clustering into metareactions. As in the original Compass original publication, reactions were then submitted to a two-sided Wilcoxon test, using the datasets as the two groups. False discovery rate was calculated using the Benjamini Yekutieli method (Yekutieli and Benjamini, 1999).

### Comparison of cortical and whole-brain organoids

To assign high-resolution cell types to cortical cells from whole-brain organoids, cortical cells were extracted from those datasets, i.e. clusters with high expression of *FOXG1* and *EMX1*, and submitted to a RF classifier created from 3- and 5-month cortical organoid datasets, using the same process as used above when classifying fetal cells. Module scores for metabolic gene sets were calculated using the process described in the Module scores section above.

### Impact of metabolic genes on cell classification

In order to evaluate the effect of metabolic genes on fetal cell classification to organoid cell types and vice versa, 38 gene sets were downloaded from the MSigDB v7.2 (Liberzon *et al*., 2011). For each gene set, genes in the set were excluded from the list of genes used to build the RF model. The proportion of fetal cell types assigned to each organoid cell type was then compared to those from the model using all organoid variable genes. To assess which changes were higher than may be expected by chance, a background distribution was built. For each timepoint and pathway, for 10,000 repetitions, a randomly selected list of genes was removed from the variable genes in the RF model and the model was re-built. The background sets of genes were the same size as the MSigDB gene set, and were expression matched, using the same method as used by Seurat’s AddModuleScore function to choose background gene sets (briefly, genes are grouped into 24 bins by average expression in the organoid dataset, and for each gene in the MSigDB gene set, a randomly selected gene from the same bin is added to the background set). P values for each fetal to organoid cell type pair were calculated by evaluating the fraction of times the change in cell type proportions in the background distribution were higher than or equal to the change induced by removing metabolic pathway genes. False discovery rate was calculated using the Benjamini Yekutieli method (Yekutieli and Benjamini, 1999).

To evaluate the expression of metabolism related gene sets across topological location, gene set expression of the MSigDB gene lists was evaluated in the Slide-seqV2 data by plotting 100x the sum of the reads for all genes in the gene set that appeared in the data, normalized by the number of UMI for each bead. Expression scores for each bead were then plotted against distance to the edge of the organoid. The edge distance was produced by calculating the convex hull of the set of beads using the chull R function and for each bead, calculating the minimum distance to the convex hull. This was done by calculating the perpendicular distance from the bead to each line segment of the convex hull and taking the minimum distance of this set. Finally, the relationship of edge distance to expression scores was plotted with geom_smooth from R’s ggplot2 library v3.3.3 (Wickham, 2016), using the Loess smoothing function and default parameters.

### Epigenetic changes associated with metabolism

To determine epigenetic changes in cells with high expression of metabolic gene sets, we use the SHARE-seq data. Module scores for the Hallmark Glycolysis and Hypoxia gene sets were assigned to cells based on RNA expression, as above. For differential metabolic peak analysis, high scoring cells were identified as those with a hypoxia score > 0.2. For each peak, the test statistic was defined as (fraction accessible in high-hypoxia cells) – (fraction accessible in low-hypoxia cells). A null distribution was calculated by permuting cell labels 1,000 times. The mean and standard deviation from this were used to calculate a Z-score and p value for each peak. Significant peaks were called those with an FDR-adjusted p-value less than 0.1. Metabolic TF correlation was calculated separately for each cell type at each timepoint by finding the Spearman correlation between each TF and each metabolic score.

### Trajectory inference across time

Cells of non-dorsal lineage (i.e., subcortical cells and cortical hem, in addition to unspecified PN) were removed from the Seurat objects of each timepoint; all objects were merged, and then we performed PCA on the resulting object. The first 15 PCs were then used as an input for the FLOWMAP algorithm (Ko *et al*., 2020) to build a force-directed graph of the datasets, by first clustering cells from each timepoint into 3,000 clusters and then building a graph connecting these clusters, using the “single” mode, the k-means clustering algorithm, Euclidean distance between clusters, and allotting 2-5 edges during each edge-building step of the algorithm. FLOWMAP only allows edges between clusters belonging to the same or consecutive timepoints. The graph, with node coordinates calculated by FLOWMAP, was visualized in Cytoscape v.3.8.2 (Shannon *et al*., 2003).

### Differential expression over time

Because ambient RNA could differ between timepoints and contribute to differential expression results, ambient RNA was estimated and removed from organoids using the decontX function from the Celda R package, version 1.6.1 (Yang *et al*., 2020), with each organoid treated as its own sequencing batch. For each cell type, DESeq2 was used to identify genes that are differentially expressed across organoid age, in days. Significant genes (those with FDR-adjusted p values of less than 0.05) for each cell type were split into two lists based on whether their log-fold-change was positive or negative. In cases where these lists included more than 1000 significant genes, they were truncated to the 1000 genes with the lowest p values. GO term enrichment was calculated for these lists using the enrichGO function from the clusterProfiler R package. In order to visualize terms with minimal overlaps, significant GO terms were run through the simplify function with a similarity cutoff of 0.4.

### Lineage branching tree trajectory

Developmental trajectories were built in organoids in a form of a branching tree using R package URD v1.1.1 (Farrell *et al*., 2018). The diffusion map was computed using the R package Destiny v2.14.0 (Angerer *et al*., 2016) by invoking calcDM function from URD on normalized counts from the organoid datasets (KNN=100 and sigma.use=30 for 10x data, and KNN=100 and sigma.use=30 for SHARE-seq data). The root cells were assigned to be the aRG cells from our youngest sample (i.e., 23 days aRG). Simulating diffusion from the root to each cell, cells were ordered in pseudotime by floodPseudotime and floodPseudotimeProcess functions with the default parameters. Terminal states of the tree (“tips”) were manually selected leveraging biological knowledge. Astroglia, oligodendrocyte precursors, CPN and CFuPN (from all timepoints at which they were present) were selected as tips for scRNAseq data, and CPN and CFuPN for SHARE-seq data. To recover the developmental trajectories in the data, we performed biased random walks starting from each tip. This walk is biased because the transitions are only permitted for cells with younger or similar pseudotimes. Parameters of the logistic function used to bias the transition probabilities were determined using the pseudotimeDetermineLogistic function (optimal.cells.forward = 40 and max.cells.back = 80). A biased transition matrix was obtained from the pseudotimeWeightTransitionMatrix function. Ten thousand simulated random walks were performed per tip using simulateRandomWalksFromTips function. Walks were then processed into visitation frequencies using processRandomWalksFromTips function. Finally trees were built using the buildTree function with the following parameters for both trees: visit.threshold = 0.7, minimum.visits = 10, bins.per.pseudotime.window = 5, cells.per.pseudotime.bin = 80, divergence.method = “preference”, p.thresh = 0.05.

### Gene trajectories from the branching tree

To identify marker genes for each trajectory, we used the aucprTestAlongTree function from the URD package. Starting from a tip, this function compares cells in each segment pairwise with cells from each of that segment’s siblings and children (cropped to the same pseudotime limits as the segment under consideration). Genes were considered differentially expressed if they were expressed in at least 10% of cells in the trajectory segment under consideration (frac.must.express = 0.1), their mean expression was upregulated 1.5x compared to the sibling and the gene was 1.25x better than a random classifier for the population as determined by the area under a precision-recall curve. Genes were considered part of a population’s cascade if, at any given branch point, they were considered differentially expressed against at least 50% of their siblings (must.beat.sibs = 0.5), and they were not differentially upregulated in a different trajectory downstream of the branch point. To determine gene expression onset and offset, we used geneSmoothFit function. Looking at the averages gene expression for a group of genes and cells in a moving window through pseudotime (moving.window = 5, cells.per.window = 25), it applied smoothing algorithms (method=spline) to describe the expression of each gene. Genes were then ordered by the pseudotime value at which they enter and then leave ‘peak’ expression (expression 50% higher than minimum value), and enter and then leave ‘expression’ (expression 20% higher than minimum value), in that order.

### Genes associated with tree branching points

To define branch-point-associated genes, we selected cells adjacent to the branch points (0.04 pseudotime units before and after) and calculated differentially expressed genes between parent and sibling branches (FindAllMarkers function from Seurat R package, min.pct = 0.1, logfc = 0.25, test.use=wilcox). To select for the genes that vary in pseudotime, we first identified variable genes in the data (Seurat’s FindVariableFeatures function) and then fitted a Lasso regression model using glmnet function from the R package glmnet 2.0-16. (Friedman, Hastie and Tibshirani, 2010) (family = “gaussian”, type.measure = “mse”, nfolds = 10). The best lambda value that minimizes mean squared error (MSE) was obtained using cv.glmnet function. To find the top distinguishing features at a given branch point, a Gradient Boosting Classifier was trained using scikit-learn 0.22.1 (Pedregosa *et al*., 2011), with the union of genes from the differential expression and regression analyses. A grid search was performed with tenfold cross-validation to optimize the maximum depth (3, 4, 5) and number of estimators (25, 50, 75, 100), which resulted in the best depth of 4 and the best number of estimators 100. Feature importance score was calculated on the basis of maximal estimated improvement by splitting on the feature under consideration against not-splitting (measured in terms of MSE), using the default option in sklearn, “friedman_mse”. The expected amount of improvement is summed over all internal nodes (where splitting occurs) of a single tree, and then summed over all trees in the gradient boosted tree model to get a single number per gene. TFs were identified using the CIS-BP database (Weirauch *et al*., 2014).

### Branching tree trajectory from SHARE-seq data

For the SHARE-seq tree, a URD tree was constructed as above using the RNA gene count data from cells that contained both RNA and ATAC data. After constructing the tree, the underlying gene count data was substituted with the ATAC peak data, while maintaining the RNA tree embedding. To find enriched TF motifs along pseudotime, cells were cut into seven sections along the tree. The parental branch was cut into three equal divisions (pseudotime from 0 to 0.1057075, then to 0.211415, then to 0.3171224), the CPN branch into two equal divisions (from 0.3171224 to 0.44005, then to 0.5629775) and the CFuPN branch into two equal divisions (from 0.3171224 to 0.4151327, then to 0.5131430). Cells from each of these sections were grouped, and DARs were calculated for each group using Signac’s FindMarkers function, with the logistic regression framework, with the number of fragments in peak regions as a latent variable, and with min.pct = 0.05. Enriched motifs per group were calculated using Homer as in the “ScATAC-seq motif enrichment” section above.

### Human callosal projection neuron diversity

Data from adult CPN from (Hodge *et al*., 2019) were downloaded, and the exon count matrix was used to create a Seurat object, along with their metadata. Counts were normalized using the same method as for the organoid scRNA-seq data. The five CPN populations were used to define gene modules that are specific to each CPN subtype using consensus NMF (cNMF) (Kotliar *et al*., 2019). The k parameter of cNMF was set to 5 since there are five different CPN clusters in the original dataset. With five cNMF modules identified, the specificity of each module to a CPN subtype was visualized by plotting the cell loading (H matrix). Gene modules were then curated by taking the top 100 genes with the highest feature loading (W matrix) for each cNMF module. To assess CPN diversity in the organoid and fetal datasets, CPN were isolated from each dataset and a UMAP was calculated for each of them, using the module genes as the variable genes. The expression of the gene modules was visualized using Seurat’s AddModuleScore function. The cNMF algorithm (originally written in Python) was implemented in R by Sean K. Simmons and can be found in our GitHub.

